# A detailed molecular picture of protein folding during active translation

**DOI:** 10.64898/2026.07.03.736445

**Authors:** Amir Bitran, Carlos Bustamante, Susan Marqusee

## Abstract

All proteins can begin to fold on the ribosome, and many rely on co-translational folding to attain their native conformation. This process is not accounted for using structure prediction algorithms such as AlphaFold and its molecular details remain largely unknown. Here, we develop a hydrogen-deuterium pulse-labeling approach which reveals nascent polypeptide folding at a high level of structural detail and its kinetic coupling with translation. Two proteins exhibit hierarchical folding of structures smaller than a domain during translation. A third protein, however, does not have time to conformationally equilibrate on the ribosome, instead becoming kinetically trapped. This subsequently biases post-translational folding to avoid an aggregation-prone intermediate populated during refolding from denaturant. Our results reveal diverse strategies that promote robust protein folding during non-equilibrium translation.

## Main Text

The specific molecular mechanisms that ensure robust protein folding in the cell, and their associated breakdown in misfolding disease, remain enigmatic (*1,2*). These protein-folding dynamics are not accounted for by native-structure prediction tools such as AlphaFold (*3,4*). All proteins have the opportunity to begin folding during their biosynthesis on the ribosome, where a nascent polypeptide chain experiences a dynamically evolving energy landscape as elongation proceeds and amino acids are added to its C terminus. Indeed, this directional, non-equilibrium process of co-translational folding has been shown to be critical for ensuring robust folding of certain model proteins that do not efficiently refold once fully synthesized (*5–10*). These benefits of co-translational folding likely extend broadly given the appreciable fraction of proteins across proteomes that cannot fold autonomously (*11–14*). In addition, co-translational folding is directly linked to proteostasis, as nascent chains sequentially engage chaperones (*15–21*) and nucleate interactions with quaternary assembly partners (*10,22–26*) at precise points in translation. Local variations in translation elongation rate have evolved to orchestrate these processes, as evidenced by conserved rare codons at putative folding intermediates (*27–32*) and enhanced ribosome pausing events correlated with chaperone recruitment (*15,33,34*).

Despite its critical importance, we do not yet understand the detailed molecular mechanisms underlying co-translational protein folding. Recent experimental studies have succeeded in monitoring the compaction of a nascent chain during real-time elongation at high temporal resolution using optical tweezers(*35,36*) and intramolecular FRET or PET (*37–40*). But a high-resolution structural description of the process is still lacking. In contrast, refolding from denaturant can be probed at near residue-level resolution using pulse-labeling hydrogen deuterium exchange (HDX) coupled to mass spectrometry (MS) or NMR (*41–45*). Recent studies have managed to extend HDX-MS methodology to stalled ribosomal nascent chains (RNCs), providing an intricate structural and dynamic characterization of RNC complexes at equilibrium including the detailed effect of chaperone interactions (*10,17–19*). But co-translational folding is intrinsically a non-equilibrium process, and thus attaining a complete picture of the process necessitates probing time-resolved folding during elongation. This requires solving two major technical hurdles: 1) monitoring the kinetic coupling between active elongation and folding, and 2) detecting the nascent chain deuteration (or the analogous structural readout in other biophysical techniques) amidst the complex background of ribosomal proteins, elongation factors, and nucleic acid components involved in the reaction. Here, we address these challenges by developing a novel application of pulse-labeling HDX-MS and provide a high-resolution time-resolved description of the co-translational folding pathways of three proteins from distinct structural families. These studies reveal a diversity of behaviors governed by both nascent chain folding propensities and the kinetic coupling between folding and translation elongation. We further reveal how non-equilibrium folding on the ribosome can introduce long-lasting hysteresis into the folding pathway that ultimately affects the folding yield.

## Monitoring time-resolved protein folding during active translation by HDX-MS

To resolve the challenges associated with probing folding during active translation, we developed an approach to perform synchronized, single-turnover *in* vitro translation using the using PURExpress® system (*46*) (Fig. 1A-B, fig. S1A). At various translation times, we pulse-label an aliquot of the translation reaction with deuterium. This allows exposed amides along the polypeptide backbone to exchange their protons for deuterons, while buried or stably hydrogen-bonded amides remain protonated, creating a chemical “snapshot” of solvent accessibility.

**Figure 1:**
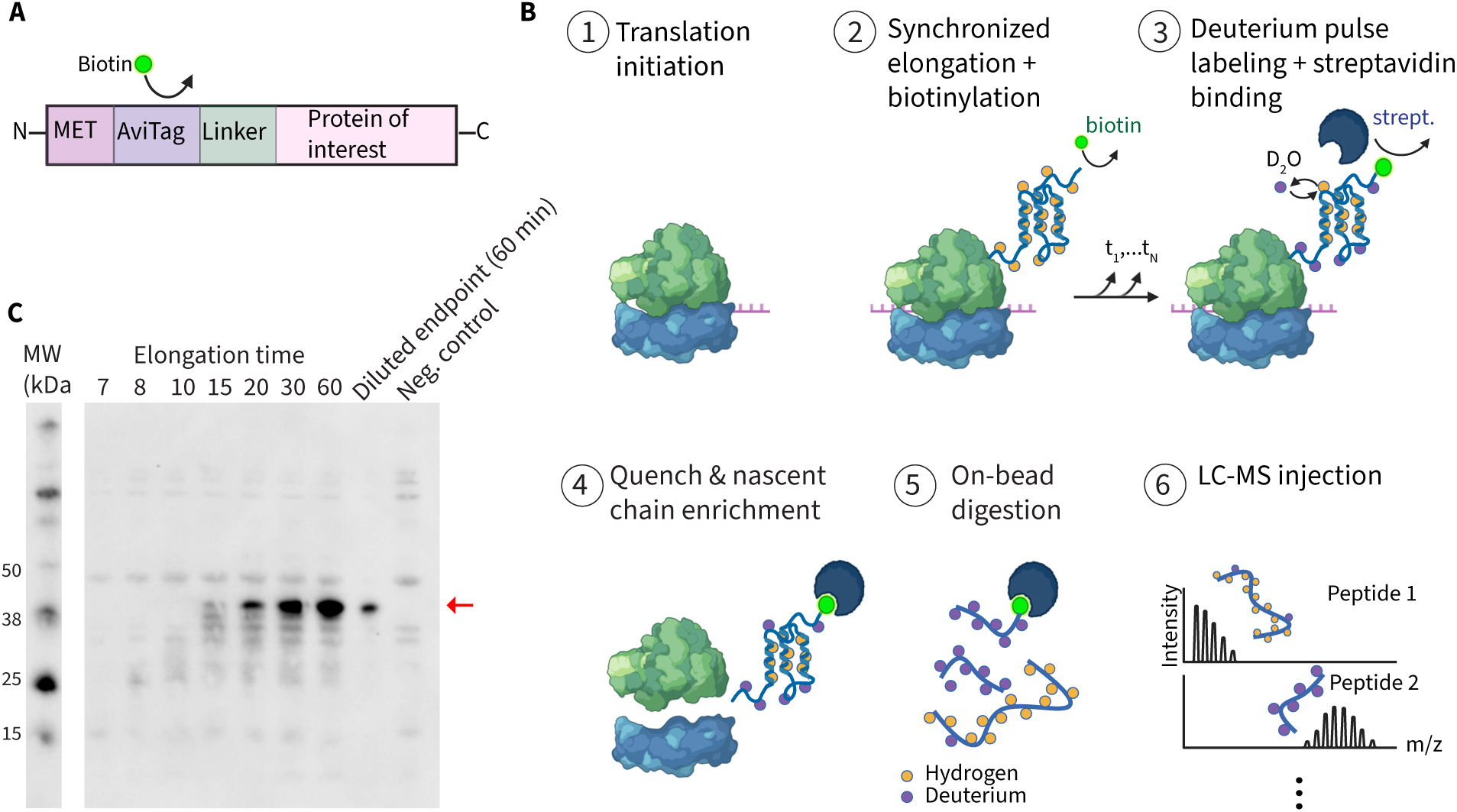
Experimental pipeline to monitor nascent chain (NC) folding during synchronized translation via hydrogen-deuterium exchange (HDX) (**A**) Map of constructs used for experiments. An N-terminal AviTag is biotinylated by the enzyme BirA. (**B**) Summary of experimental design. Synchronized translation is achieved by pre-initiating ribosomes on mRNA (step 1), followed by addition of complete amino acid mix and pre-charged tRNA to begin elongation (step 2), coupled to nascent-chain biotinylation. At various elongation times, an aliquot is pulsed with deuterium to label exposed amides and simultaneously bound to streptavidin beads (step 3). The HDX reaction is quenched via acidification, which also dissociates ribosomes while nascent chains (NCs) remain bound to beads (step 4). NCs are digested into peptides via addition of pepsin to the beads (step 5), then analyzed for deuteration via liquid-chromatography mass spectrometry (step 6). For details, see Materials and Methods. (**C**) Example anti-biotin western blot using streptavidin HRP to monitor progress of synchronized elongation of *E. coli* alpha Tryptophan synthase. Red arrow indicates full-length protein. The 60 min timepoint is diluted 5-fold in the penultimate lane, and the final lane shows a negative control translation reaction without genetic template, revealing bands resulting from non-specific streptavidin HRP binding.

Crucially, our hydrogen-deuterium exchange (HDX) reaction is coupled to biochemical enrichment of the nascent chain. To achieve this, we translate the protein of interest with an N-terminal AviTag (Fig. 1A), which is co-translationally biotinylated by the enzyme BirA. Then, during the pulse-labeling reaction, the biotinylated nascent protein is pulled down with streptavidin beads that have been pre-equilibrated in deuterated buffer. Bead binding does not disrupt RNC integrity, nor globally alter the protein conformation (fig. S1B-E). Following a ten-second labeling and binding period, we quench both the exchange and translation reactions by diluting the sample into ice-cold, low-pH buffer. All translation components are then separated from bead-bound nascent chains, and the nascent chains are then digested with protease, releasing the peptides from the beads. The peptides are then subjected to liquid-chromatography mass spectrometry (LC-MS) for detection of the amount of deuteration in the different regions in the nascent chain. This workflow successfully monitors the on-ribosome population during elongation (fig. S1F and S2, see also Supplementary Note 1) and yields high peptide coverage for all our proteins of interest (fig. S2B), indicating that we can monitor time-resolved folding at multiple regions in a protein during synchronized translation.

## Protein subdomains acquire stable structure during translation

We applied this method to monitor the co-translational folding of various proteins that have been previously predicted to fold co-translationally, beginning with *E. coli* cytidine monophosphate kinase, CMPK, (Fig. 2A) from the P-loop structural family. All-atom simulations predict that CMPK folding begins with the NMP subdomain after roughly 130 AAs have emerged from the ribosome, while core folding is delayed until after translation is complete (*29*)(fig. S3). We carried out our HDX-MS approach on a synchronized CMPK translation reaction and followed the mass spectra for multiple peptides (Fig. 2A, replicate reaction in fig. S4—see also Supplementary Note 2: Reproducibility analysis) spanning all three subdomains of this protein (spectra in Fig. 2B and fig. S4A).

**Figure 2:**
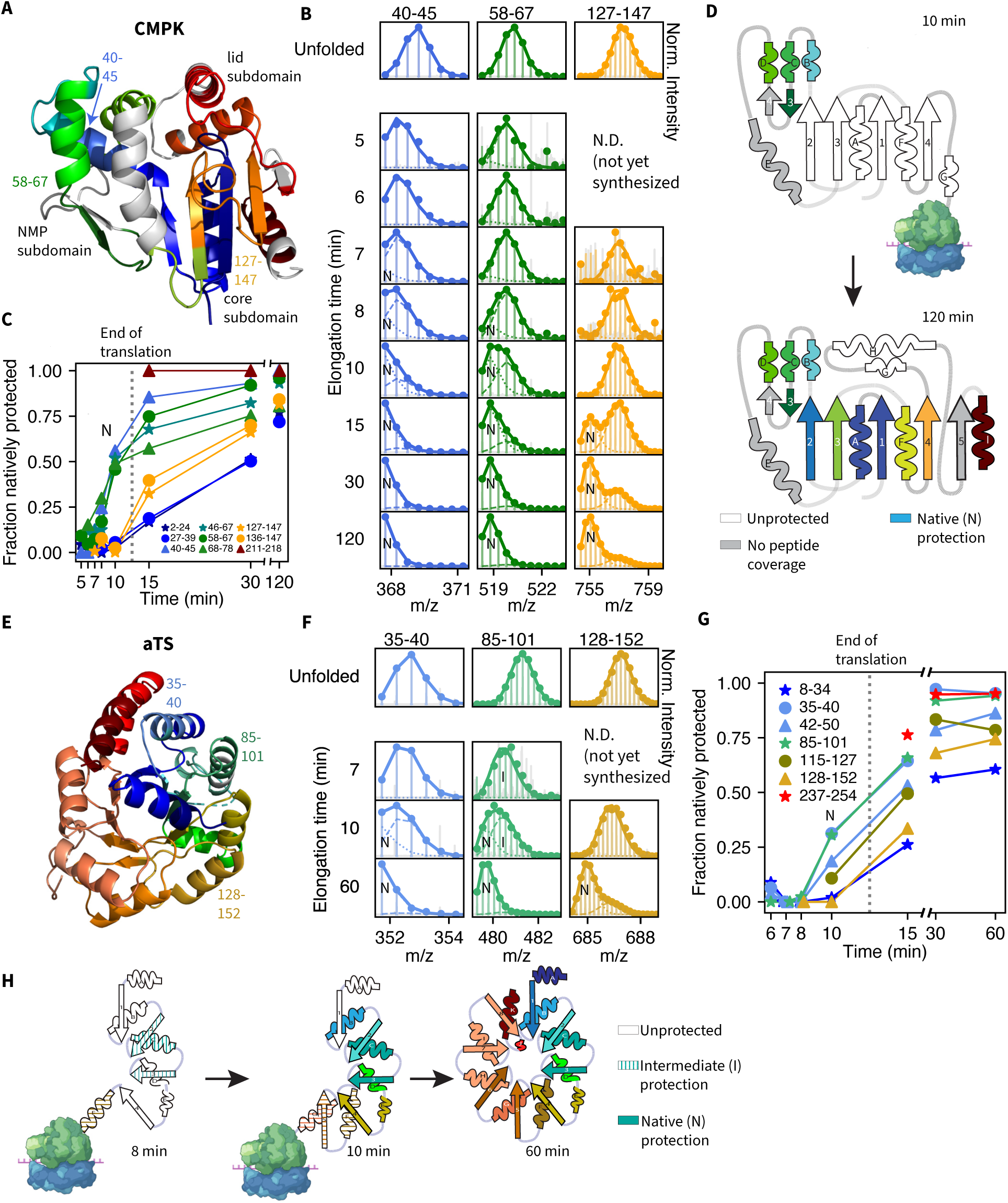
Subdomains begin folding during active translation. **(A)** Structure of *E. coli* CMPK (PDB ID *2CMK*) with colored HDX-MS reporter peptides, three of which are labeled. **(B)** Mass spectra (translucent solid lines) vs. elongation time of example deuterated peptides, alongside maximally deuterated spectra (“Unfolded”). Round markers denote isotopic peaks, opaque solid lines indicate fits, and dashed lines show individual mode contributions (N=natively protected mode). **(C)** Fractional population associated with natively protected mode vs. elongation time for indicated peptides. **(D)** Schematic of CMPK co- translational folding. Structures spanned by peptides showing >40% natively protected population at the indicated times are colored solid, provided the native mode is ≤67% deuterated. All other peptides are shown in white, and gray denotes structures lacking peptide coverage. **(E)** Structure of *E. coli* aTS (PDB ID *1WQ5*) with color-coded reporter peptides. **(F)** Mass spectra for indicated aTS peptides as in (B) **(G)** Same as (C) for reporter peptides from aTS. **(H)** Same as (D) for aTS. In this case, solid colors refer to peptides with >10% natively protected population, while dashed colors denote peptides showing statistically significant, albeit partial intermediate protection (see Materials and Methods).

To elucidate the CMPK folding trajectory, we globally fit the mass spectra associated with each peptide to multiple modes reflecting different levels of deuteration (Fig. 2B-C). For most CMPK peptides, two modes are required, a protected and an unprotected mode (Fig. 2D), as determined by a Bayesian information criterion approach (Materials and Methods, fig. S5). The pattern of deuteration for the protected mode correlates with the expected native secondary structure of CMPK (fig. S4D). Thus, as a proxy for native-like folding, we monitored the fractional population associated with this protected mode as a function of elongation time across all peptides (Fig. 2C and fig. S4). Peptides from the NMP subdomain show a concerted uptick in native protection after 8 mins of elongation, at which point this subdomain has fully emerged from the exit tunnel in the fastest-translating ribosomes (fig. S4B). This result strongly suggests that this subdomain folds co-translationally. In contrast, the core subdomain does not fold until after synthesis is complete (15 mins timepoint), consistent with the core beta-sheet topology involving long-range contacts between the N-and-C termini. We note that for some peptides, a minor population remains unprotected, likely due to a small fraction of prematurely stalled ribosomes that do not complete synthesis (fig. S4E). Peptides from the lid subdomain show only partial protection even in the native state (fig. S4A-C), consistent with the known flexibility in its structural homolog adenylate kinase (*47*). Thus, CMPK folds through a sequence of intermediates beginning with early co-translational folding of the N-terminal NMP subdomain, followed by the slower rearrangement of the core subdomain, consistent with simulation predictions (Fig. 2D and fig. S3).

We next turned to the TIM barrel family, hypothesizing that we may observe sequential co-translational folding given that the N-terminal halves of some TIM barrels stably and rapidly fold in isolation (*48,49*). We performed pulse-labeling HDX-MS on a synchronized translation reaction of the well-characterized protein, *E. coli* alpha Tryptophan Synthase (aTS, Fig. 2E, replicate reaction in fig. S5). Indeed, we find that aTS folds co-translationally (Fig. 2F-H), beginning with early acquisition of partial protection of N-terminal structures, particularly the third alpha helix and beta sheet (peptide 85-101, Fig. 2F mode “I”). This is followed by an uptick in native protection from the second alpha helix through the fourth beta strand, and finally, post-translational protection of the C-terminal half barrel and the first beta strand, which must pair with the final strand to close the barrel (Fig. 2G-H). The co-translational folding intermediate seen here closely resembles an intermediate observed upon refolding from denaturant (*50*) (fig. S6G), indicating that aTS folds via a similar mechanism on and off the ribosome.

## Translation alters the folding pathway of aggregation-prone HaloTag upon release from ribosome

The protein HaloTag (containing an α/β hydrolase fold) achieves a higher yield of natively folded protein during *in vitro* translation compared to refolding from a denatured state (*9*). To uncover the molecular basis underlying this observation, we performed pulse-labeling HDX-MS on a synchronized HaloTag translation reaction (Fig. 3, replicate reaction in fig. S7)— HaloTag acquires its native structure at the end of translation as shown by its ability to covalently bind the fluorescent ligand TMR (fig. S7C) (*9*). Pulse-HDX reveals that across the entire protein, emergence of native-like protection is delayed until after translation is complete at which point folding begins with the C-terminus of the core. The N-terminus of the core and lid subdomains fold much later (Fig. 3B, C and F). This sequence of native-structure acquisition contrasts with HaloTag’s folding pathway upon dilution from denaturant (*43*) where the first detectable intermediate involves both the N- and C-terminal parts of the core (Fig. 3E and G and fig S8). The delayed N-terminal folding relative to the C-terminus in this translation-based experiment is reproducible across replicate translation reactions (fig. S7) and cannot be accounted for by lagging ribosomes (Supplementary Note 3). Instead, this delay indicates that translation induces an N-terminal conformation that is kinetically trapped. The slow resolution of this trap biases post-translational folding to proceed through an alternative pathway, starting from the C-terminus. In support of this model, peptides arising from various N-terminal regions exhibit subtle, but significant nonnative protection early in translation (Fig. 3D) distinct from the burst-phase protection pattern seen upon dilution from denaturant (fig. S8B-D). This is consistent with a translation-induced conformational trap. In summary, translation biases HaloTag’s folding pathway to proceed via a different pathway than upon dilution from denaturant and circumvents the refolding intermediate populated during refolding that has previously been implicated in aggregation (*9,43*). This likely accounts for the observation that HaloTag achieves a higher folding yield upon translation than upon refolding from denaturant.

**Figure 3:**
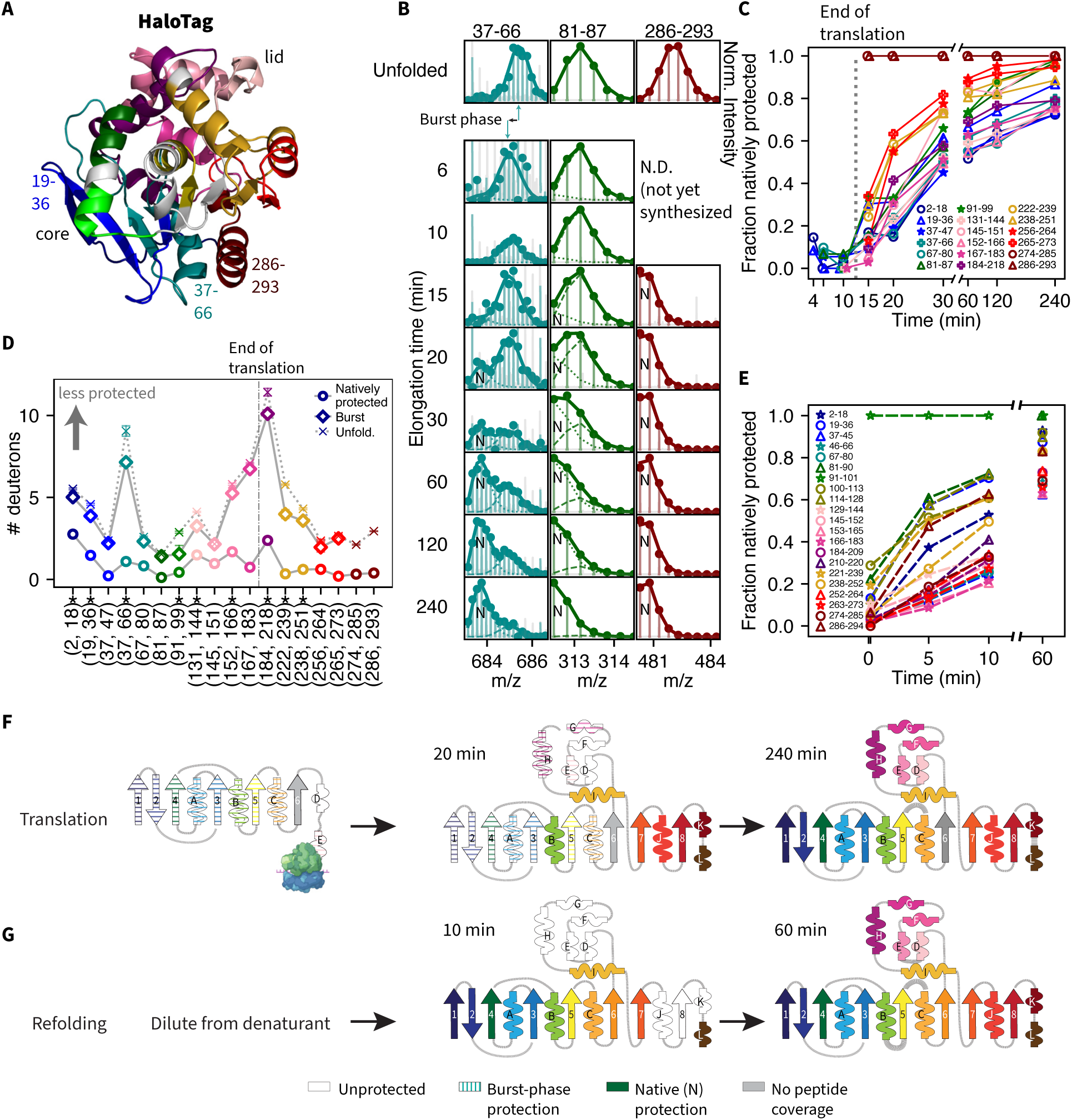
Translation biases HaloTag folding to circumvent aggregation-prone intermediate. **(A)** Structure of HaloTag (PDB ID 5Y2Y) with peptides used in HDX-MS analysis colored and mapped onto the structure and subdomains indicated. **(B)** Deuterated peptide mass spectra vs. elongation time for indicated peptides alongside fully deuterated control as in Fig. 2B, highlighting burst-phase protection for 37-66. **(C)** Native population vs. elongation time as in Fig. 2C. **(D)** Number of deuterons associated with natively protected mode (circles), burst-phase mode (diamonds) and unfolded state (Xs) for each peptide. Error bars represent 95% confidence intervals from bootstrapping and asterisks above labels denote peptides exhibiting statistically significant burst-phase protection (Materials and Methods, illustrated in Fig, 2B). Peptides to the right of the dashed line are only detected after 10 mins (post-translation). **(E)** Same as (C) for HaloTag refolding from 7.5M urea, with data re-analyzed from reference (***43***). **(F)** Schematic of HaloTag co-and-post translational folding with solid colors highlighting structures spanned by at least one peptide showing at least 50% natively protected fraction at the indicated times, striped colors indicating significant burst-phase protection, white indicating no protection, and gray indicating no peptide coverage. **(G)** Same as (F) now depicting Halo Tag refolding from 7.5M urea.

## Nascent HaloTag can fold on the ribosome given enough time to reach equilibrium,

HaloTag folds on a significantly slower timescale than that of translation itself (Fig. 3C), suggesting that nascent chains may not have time to reach conformational equilibrium during active elongation. To test this, we realized the equilibrium scenario by generating stalled ribosomal nascent chains (RNCs), which we incubated for four hours before isolation via sucrose cushion ultracentrifugation and then interrogated their folding status via pulse-labeling HDX as was done for actively translating NCs. These RNCs contained HaloTag N-terminal fragments of various lengths coupled to a strong SecM stalling sequence (*51*) and an N-terminal AviTag (Fig.4A-B). Our second-longest length, HaloTag 280, recapitulates the scenario where HaloTag is fully translated but not yet released from the ribosome, while our longest length (F.L. + 34 AA) contains an additional C-terminal linker to fully span the ribosomal exit tunnel, such that the native HaloTag sequence has fully emerged from the ribosome. We confirmed that our HDX pulse-labeling reports on RNCs that are stably attached to the ribosome (Fig. 4C) (*52*).

**Figure 4:**
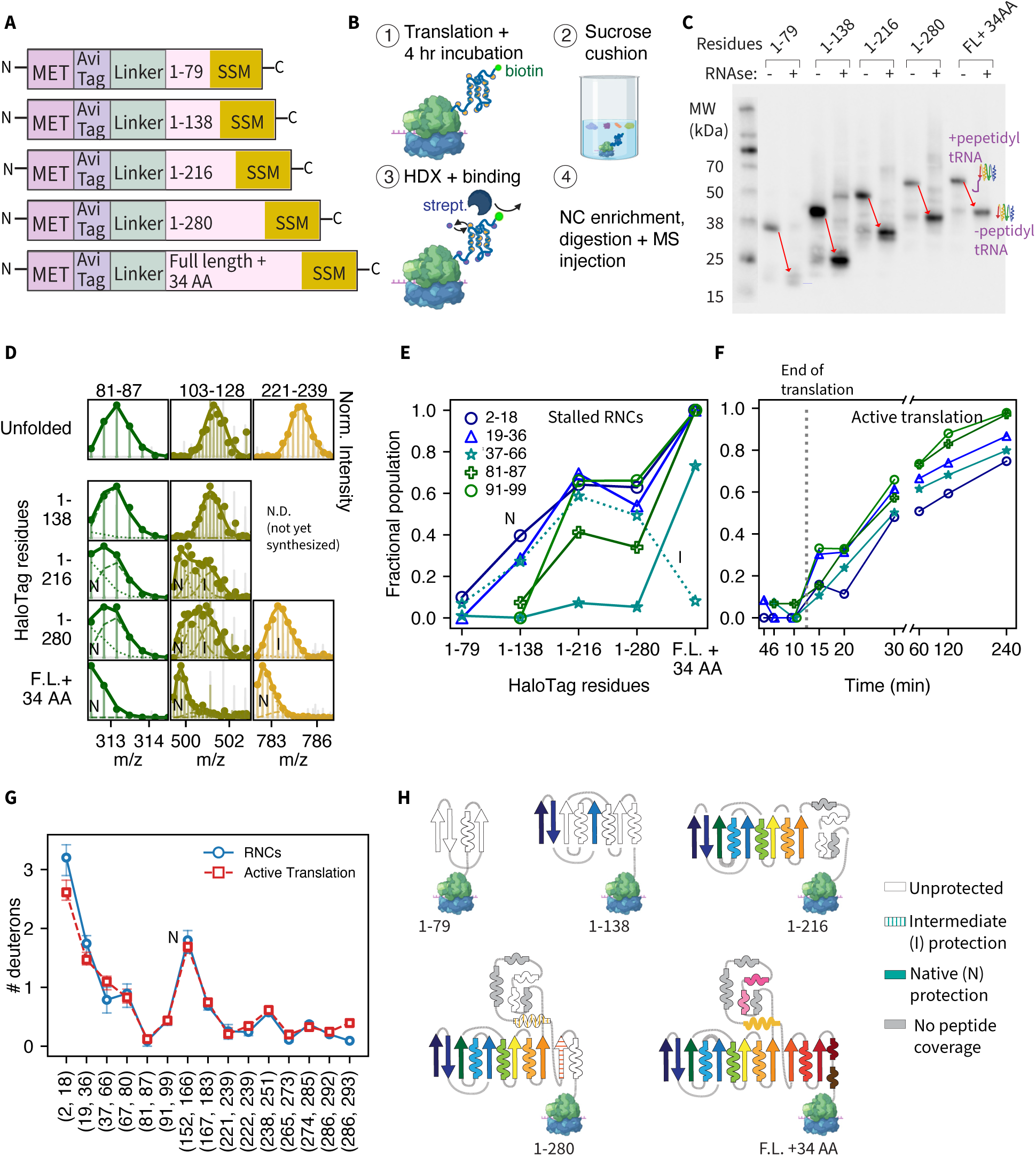
HaloTag nascent chains do not reach equilibrium during active elongation. **(A)** Diagrams of constructs used for stalled ribosomal nascent chain (RNC) experiments including strong SecM (SSM) stalling sequence. **(B)** Experimental workflow for RNC preparation and analysis by HDX-MS--see Materials and Methods. **(C)** Streptavidin-HRP western blots showing RNAse-dependent migration shift in RNCs due to peptidyl tRNA attachment (red arrows). **(D)** Deuterated mass spectra for representative peptides from equilibrated RNC constructs alongside maximally deuterated condition (“Unfolded”). **(E)** Fitted fractional populations associated with natively protected mode for indicated peptides from each RNC construct (opaque markers), and intermediate mode for peptide 37-66 (empty markers) obtained from two HDX technical replicates of each sample. **(F)** Same as Fig. 3C for subset of peptides shown in E. **(G)** Number of deuterons associated with natively protected mode (N) for each peptide detected in both HaloTag RNC and active translation experiments. Error bars represent 95% confidence intervals from bootstrapping. **(H)** At each length, secondary structures are shown with solid colors if spanned by at least one peptide showing ≥25% natively protected population, with dashed colors if partially protected, in white if unprotected, and in gray if the region lacks peptide coverage.

In contrast to actively translating HaloTag, which remains predominantly unprotected on the ribosome (Fig. 3B), equilibrated RNCs show a length-dependent increase in protection across multiple structural regions (Fig. 4D and fig. S9). Various peptides show bi-or-trimodal mass spectra indicative of multiple slowly interconverting populations. We thus quantify length-dependent protection by globally fitting all mass spectra to a combination of modes reflective of these populations (Fig. 4E). At short lengths, N-terminal peptides show minimal, albeit significant protection relative to the unfolded state (Fig. 4D, E, and fig. S9C), resembling the nonnative protection seen early during active translation of HaloTag (fig. S9D and E). However, at increasing RNC lengths, the N-terminus of the core populates increasingly protected states in a length-dependent manner (Fig. 4D, E and H), although this relationship with length is not perfectly monotonic, potentially due to complex stability effects related to distance from the ribosome (*53–55*). Interestingly, the regions that become protected closely match those protected in the denaturant refolding intermediate (Fig. 4H and 3G), suggesting a similar structure may form in equilibrated RNCs. Moreover, the native-mode protection in RNCs nearly perfectly correlates with that observed at the end of active translation (Fig. 4G, fig. S9D and E), strongly suggesting the protected RNC intermediates are native-like. But crucially, while equilibrated RNCs show length-dependent native protection of various core structures, these same regions do not show comparable protection during active translation until well after synthesis is complete (Fig. 4F). Thus, actively translating HaloTag remains out of equilibrium, as nascent chains do not have time to natively fold while on the ribosome despite their ability to do so.

## Discussion

We developed and implemented an HDX-MS approach to monitor co-translational folding across the nascent chain during active elongation. The ability to follow folding at a high level of structural detail during active elongation provides important insights inaccessible to high-resolution structural studies on stalled nascent chains (*10,17–19,53–55*) or time-resolved studies of compaction during elongation (*35–40*). We demonstrate that co-translational folding is shaped by both the thermodynamic drive to fold and the coupling between the relative kinetics of elongation and folding. This coupling results in diverse behavior that is protein dependent. For CMPK and aTS (from the P-loop family and TIM barrel family, respectively), co-translational folding proceeds through a hierarchical series of intermediates involving N-terminal structural elements smaller than a complete domain (Fig. 5, top pathway). Although subdomain-level folding during translation has long been postulated (*56*), including for TIM barrel proteins (*57*), our work provides direct experimental evidence of this process during active elongation.

**Figure 5:**
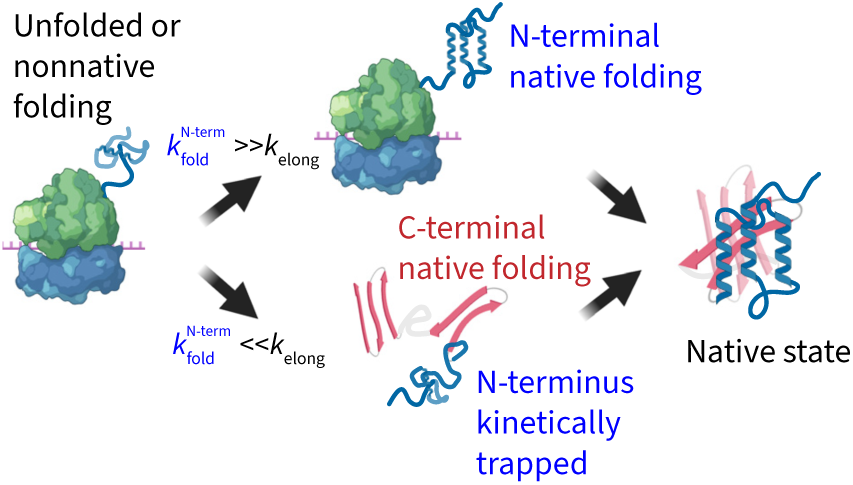
Nascent chain folding depends on kinetic competition between translation and folding rates. Cartoon depicting kinetic competition between elongation and folding during active translation. For details, see discussion.

Atomistic simulations of CMPK using the DBFOLD algorithm (*58*) recapitulate our observed folding hierarchy and generate experimentally consistent structures of nascent chain intermediates (fig. S3), further highlighting the power of this and related approaches to predict and interpret co-translational folding mechanisms (*40*). For aTS, the co-translational intermediate we observe closely resembles an obligatory N-terminal denaturant refolding intermediate previously observed by HDX-MS (fig. S6G), suggesting the two folding mechanisms are similar (*50*). But, for the large number of TIM barrels that do not refold efficiently from denaturant (*11,13,14*), sequential co-translational folding as observed here may be critical for ensuring robust folding of this topologically complex structure. More broadly, this strategy may enable proteins to circumvent deep post-translational folding traps (*5–8,29,30,59*).

In contrast, HaloTag does not begin exhibiting native-like folding until after its release from the ribosome, even though the stalled RNC experiments demonstrate that nascent HaloTag is capable of folding on the ribosome. The timescale required for it to do so is much longer than the timescale of translation elongation. Instead, actively translating HaloTag shows minimal protection from HDX on the ribosome, but counterintuitively begins folding from the C-terminus upon its subsequent release from the ribosome (Fig. 5 bottom pathway). This indicates that the N-terminus becomes kinetically trapped during elongation and thereby doesn’t sample the aggregation-prone intermediate populated during refolding from denaturant which involves both termini. This intermediate resembles the conformation formed by stalled HaloTag RNCs that have been given enough time to reach equilibrium (Fig. 3G and 4H). Thus, the non-equilibrium nature of co-translational folding is likely crucial in allowing the protein to avoid this problematic intermediate through a chaperone-independent mechanism and thus attain a high folding yield (*9*).

The structural nature of the kinetic trap during active HaloTag translation remains to be fully elucidated. Nevertheless, our observation of significant, albeit partial, nonnative protection at early translation times are consistent with either 1) a single conformation with a small subset of native hydrogen bonds formed, or 2) a dynamic ensemble of nonnative conformations that readily exchange with each other, akin to that observed on the ribosome for the protein HemK (*37–40*). In either case, the barrier to escape this state must be large enough to trap the N-terminus until well after translation is complete, thus biasing the folding pathway. More broadly, our results underscore the importance of assays like the one developed here to monitor folding during active elongation, as non-equilibrium folding during translation can differ starkly from that seen in equilibrated RNCs, as noted in single molecule mechanical studies on calerythrin (*36*).

It is important to note that the rate of translation elongation in PURE-based *in vitro* translation assays such as ours (∼0.5-1 AA/sec, consistent with previously measured rates PURE system (*9*)) is significantly slower than the *in vivo* elongation rate in bacterial systems such as *E. coli*, from which PURE components are derived (∼10-20 AA/sec). However, for the case of HaloTag, we expect that a faster rate of translation would only *further* heighten the discrepancy between equilibrium and real-time co-translational folding, provided the kinetic trap we observe forms on rapid timescales. In contrast, aTS very rapidly refolds from denaturant (<1 sec timscales) into an obligatory N-terminal refolding intermediate closely resembling the co-translational intermediate observed here (*50*) suggesting it may also form under a faster elongation rate. Closing the gap between the *in vitro* and *in vivo* elongation rates will enable detailed investigations on the effects of the relative rates of translation. This includes the role of conserved rare codons such as those in the CMPK sequence shortly after the NMP-binding subdomain (*27,29*) which folds co-translationally in our assay.

In summary, our work reveals a diversity of strategies that are deployed by the ribosome to promote robust protein folding, from promoting sequential folding of subdomain elements to *delaying* folding to avoid problematic intermediates. Over the course of evolution, a protein’s fitness may depend on its ability to exploit these folding mechanisms, and future work is needed to extend equilibrium biophysical fitness models (*60–62*) to account for non-equilibrium co-translational folding. Viral proteins, in particular, rely heavily on such mechanisms to fold robustly during infection (*63*). In the cell, co-translational folding is further influenced by chaperones and quaternary assembly partners that interact with nascent chains. In the future, the methodological platform presented here can be extended to account for these more complex mechanisms and reveal possible links between co-translational (mis)folding and disease. Finally, these findings can be exploited to enhance the efficiency of *de novo* protein design and transgenic protein expression by optimizing for kinetic folding yield.

## Acknowledgments

We thank Hossein Amiri for assistance with tRNA pre-charging and David Balchin for detailed technical advice on experimental optimization and troubleshooting. We thank Marqusee lab members Vrishab Aravind for assistance with early methods-development experiments, Shikha Kathrani, Mark Petersen, Haley Moran, and Joshua Morse for hands-on assistance with synchronized translation experiments, as well as Darren Kahan, Mark Petersen, and Sophie Shoemaker for assistance with LC-MS method development and instrument troubleshooting. We thank all current and former members of the Marqusee and Bustamante labs for helpful discussions and feedback. We thank Shawn Costello, Mark Petersen and Darren Kahan for carefully reading and providing thoughtful feedback. Figures 1A and 2D and H, 3F, 4B and H, and 5 were created with assistance from BioRender.com. S.M. is a Chan Zuckerberg Biohub investigator. C.B. is Howard Hughes Medical Institute investigator.

## Funding

This work was supported by the Jane Coffin Childs Memorial Fund for Postdoctoral Research (to A.B.) and the National Institutes of Health grant R35GM149319 (to S.M.).

## Author contributions

Conceptualization: AB, SM Methodology: AB, SM Investigation: AB Visualization: AB

Funding acquisition: SM Project administration: SM, CB Supervision: SM, CB

Writing – original draft: AB

Writing – review & editing: AB, SM, CB

## Competing interests

Authors declare that they have no competing interests.

## Data, code, and materials availability

All data (raw peptide mass spectra, fitting parameters, unedited western blot images), as well as Python code used to generate the figures in this text, are available on Dryad at https://datadryad.org/share/LINK_NOT_FOR_PUBLICATION/iAhpdnuyJjdeCEWNpFPp52pa9 <u>WgKLhZZ8I-9m_axRwI.</u> All DNA constructs are available upon request.

## Materials and Methods

### Cloning and PCR

Genes encoding proteins of interest used for synchronized translation reactions were cloned via Gibson assembly into a plasmid encoding an ampicillin resistance marker. The gene was inserted downstream of an AviTag as well as a linker region containing a threonine, a stretch of serines and glycines (41 residues for HaloTag, 54 for aTS and CMPK), and three valines. The *cmk* and *TrpA* genes encoding the proteins CMPK and aTS were amplified directly from the *E. coli* K-12 chromosome via PCR, using primers directed against the 5’ and 3’ ends of the gene prior to Gibson assembly. The identities of the inserts on the respective plasmids were verified by Sanger sequencing.

As a template for transcription and synchronized translation reactions, the genes of interest were reamplified from their harboring plasmids using a forward primer directed upstream of the T7 promoter and a reverse primer downstream of the stop codon. The resulting PCR product was DpnI-digested to remove the template plasmid, followed by PCR purification on a silica gel column and elution using molecular biology grade water (Corning 46-000). Purity and yield were assessed via NanoDrop. Any PCR products showing an *A*_260_/*A*_230_ absorbance ratio less than 1.8 were further purified using a 0.5 mL Zeba spin desalt column (Thermo Scientific 89883) to remove residual salts.

For stalled ribosome-nascent chain (RNC) experiments, the HaloTag gene, alongside an N-terminal AviTag and linker (as described above), was cloned into a plasmid containing a 17 AA C-terminal linker (GSGSGSGSGSEFLPYRQ) followed by the 17 AA strong SecM stalling sequence (FSTPVWIWWWPRIRGPP) using around-the-horn (ATH) PCR. In total, this C-terminal region (linker + SecM) contains 34 amino acids. This plasmid was then used as a template for subsequent ATH PCRs to generate plasmids consisting of various HaloTag C-terminal truncations (HaloTag 1–79, 1–138, 1–216, and 1–280) all followed by the strong SecM stalling sequence (but no linker). Prior to *in vitro* translation, the regions spanning the T7 promoter through the strong SecM stalling sequence were amplified from each plasmid via PCR, and the resulting PCR fragments were purified as described in the previous paragraph. In addition to the truncations, we generated a PCR fragment to encode an RNC harboring full-length HaloTag sequence and the 34 AA C-terminal region (linker + strong SecM) to fully span the ribosomal exit tunnel, thus ensuring that HaloTag is fully emerged from the ribosome.

### Synchronized Translation Reactions

Synchronized translation reactions are performed in two steps: (1) mRNA transcription and initiation, followed by (2) single-turnover elongation. The transcription and initiation steps are expected to be slow; thus, it is advantageous to complete them first to maximize subsequent synchrony.

For the transcription and initiation step, reactions are prepared with methionine as the sole provided amino acid to allow mRNA to be transcribed and initiation complexes to form without elongating. This is achieved by preparing two to four PUREexpress reaction equivalents, depending on the number of timepoints to be taken for HDX-MS. Each HDX-MS timepoint requires 6 μL of material, with an additional 2 μL for western blot samples. Each reaction equivalent consists of:

- 5 μL of Solution A –aa, tRNA (New England Biolabs B6841AVIAL from E6840S)
- 7.5 μL Solution B –RF123 (NEB P6854AVIAL)
- 1 μL Murine RNAse inhibitor (NEB M0314L)
- 1.25 μL of 20 μM BirA
- 2.5 μL of *E. coli* tRNA mix (NEB N6842AVIAL)
- 0.5 ìL of 30 mM methionine
- 0.5 μL of 1 mM biotin (20 μM final)
- 260 ng of purified PCR product encoding the gene of interest downstream of an AviTag + glycine-serine linker (see previous section) or an equivalent volume of molecular biology grade water (for negative control reactions)
- Molecular biology grade water to bring the total volume to 30 μL.

The optimal biotin concentration of 20 µM was determined by a streptavidin bead binding experiment shown in fig. S1H. A formyl donor for the formation of fMet-tRNAs is present in Solution A, and the addition of tRNAs at this step provides a source of methionine tRNA. In certain control experiments, we added 1 μL of FluoroTect GreenLys (Promega L5001)—a lysine tRNA charged with a BODIPY-FL labeled lysine—to facilitate in-gel visualization of the translated product without requiring western blotting; however, FluoroTect is excluded from reactions destined for HDX-MS analysis. This reaction is incubated for 30 min at 37°C to allow transcription to occur and initiation complexes to form. Where indicated, a HaloTag TMR Ligand (Promega G825A) was added to a 10 μM final concentration at this stage.

During the initiation period, a working elongation mix is prepared by mixing 2.5 μL of complete amino acid mix (NEB N6843AVIAL); 0.5 μL each of release factor 1 (NEB P6851AVIAL), release factor 2 (P6852AVIAL), and release factor 3 (P6853AVIAL); at least 2.5 μL of pre-charged tRNA for a final concentration of >20 μM pre-charged tRNA (>1 μM each); and 0.95 μL of 2.37 mM aurintricarboxylic acid (ATCA, MilliporeSigma 18-940-0100MG) for a final working concentration of 75 μM per reaction equivalent. ATCA is used to prevent the reinitiation of ribosomes and ensure single-turnover elongation. We confirmed that translation is inhibited when ATCA is added during the initiation step, but not when added at the elongation step (fig. S1A). The ATCA solution is prepared fresh from powder on the day of use, stored on ice, and protected from light. The working elongation mix is prepared under RNAse-free conditions and pre-warmed at 37°C for 10 min prior to the onset of elongation.

After 30 min of initiation, 7.45 μL of elongation mix is added per reaction equivalent of initiation mix to begin elongation. The N-terminal AviTag in the nascent chains becomes rapidly biotinylated after its emergence, with significant biotinylation already occurring after 3 minutes as shown in fig. S1G. At the desired elongation timepoints, a 6 μL aliquot of the synchronized elongation reaction is drawn to perform pulse HDX and nascent chain enrichment (see section *Pulse HDX-MS, Maximally Deuterated Sample Prep, and Nascent Chain Enrichment*). In addition, at these same timepoints, 1.8 μL of the reaction is drawn and mixed with 3.6 μL of 2 mM puromycin (1.33 mM final) to generate samples for western blot analysis. These samples are kept at room temperature for at least 30 s and then placed on ice. They are subsequently treated with RNAse A (Thermo Scientific EN0531, 0.5 mg/mL final) for 5 min at 37°C, followed by the addition of 5.4 μL of 2 × Laemmli sample buffer prior to SDS-PAGE and/or western blot analysis. After collecting the final timepoint, 6 μL of the sample was bound to streptavidin beads pre-equilibrated in H_2_O for the preparation of an undeuterated control, and two 6 μL aliquots were used to prepare maximally deuterated samples.

Although the main text figures show one biological replicate for each active translation reaction, we performed at least two total replicates for each protein; additional replicates are shown in the supplementary figures. We verified that our main results—particularly the order of folding of distinct structural elements before and after the end of translation—reproduce across biological replicates. The precise times at which events occur vary slightly between replicates due to ribosomal batch variations that lead to minor variability in translation rates.

### Preparation of Pre-charged tRNA

Total *E. coli* tRNA (6.25 mg, Roche MRE600) was weighed and resuspended in 280 μL of molecular-biology grade water. This was mixed with 50 μL of a complete amino acid mixture containing roughly 2.2 mM of each amino acid at near-neutral pH, 20 μL of 100 mM ATP, 50 μL of 10 × charging buffer (final 1 × composition: 50 mM K-HEPES pH 7.5, 50 mM KCl, and 5 mM DTT), and 100 μL of DEAE-purified S-100 extract (*64*). This reaction was incubated for 1 h at 37°C, followed by the addition of 80 μL of NaOAc or KOAc pH 5.3 (final concentration of 300 mM) to acidify the reaction and maintain ester bond stability.

Protein components were then removed by phenol-chloroform extraction using cold, saturated phenol (pH 5.3). The tRNA was precipitated overnight at -80°C with 100% ethanol, followed by centrifugation for 2 h at 14,000 × *g* at 4°C. The supernatant was removed and the pellet was air-dried for 5–10 min, followed by resuspension in water (pH 5.3) and desalting through an illustra MicroSpin G-25 spin column (Cytiva 27-5325-01) pre-equilibrated in 3 mM KOAc. Following this stage, absorbance at 260 nm was measured using a NanoDrop to determine tRNA concentration. Pre-charged tRNA was aliquoted, flash-frozen in liquid nitrogen, and stored at -80°C.

### Preparation of Stalled Ribosomal Nascent Chain Samples

A translation master mix was prepared under RNAse-free conditions, consisting of:

- 15 μL Solution A –aa, tRNA (New England Biolabs B6841AVIAL from E6840S)
- 22.5 μL Solution B –RF123 (NEB P6854AVIAL)
- 3 μL Murine RNAse inhibitor (NEB M0314L)
- 3.75 μL of 20 μM BirA (0.8 μM final)
- 7.5 μL of *E. coli* tRNA mix (NEB N6842AVIAL)
- 7.5 μL of complete amino acid mix (NEB N6843AVIAL)
- 1.5 μL of 1 mM biotin (17 μM final)

This master mix was then split into 6 equal aliquots corresponding to the five HaloTag RNC constructs to be generated, with duplicate reactions for the L34 construct (which contains the 34 AA combined C-terminal linker + strong SecM sequence to fully span the exit tunnel, see *Cloning and PCR*) to ensure sufficient volume for undeuterated and maximally deuterated controls. To each aliquot, 130 ng of PCR product encoding the respective gene construct (including the N-terminal AviTag + linker and C-terminal strong SecM stalling sequence) was added, along with molecular biology grade water to a total volume of 15 μL. The reactions were incubated for 4 h at 37°C to allow for complete translation and equilibration of nascent chains.

Subsequently, 14 μL of each reaction was layered atop a 94 μL sucrose cushion consisting of 1 M sucrose in HKMT buffer (25 mM HEPES, 15 mM Mg(OAc)_2_, 150 mM KCl, 0.1 mM TCEP, pH 7.5) in polycarbonate centrifuge tubes (Beckman Coulter 343775). These samples were ultracentrifuged in a Beckman Coulter Optima MAX-XP ultracentrifuge for 80 min at 200,000 × *g* and 4°C using a pre-chilled Beckman Coulter TLA-100 fixed-angle rotor. Following the removal of the supernatant, the ribosomal pellets were resuspended in 20 μL of HKMT buffer, and the two resuspended pellets corresponding to the L34 construct were pooled.

All resuspended pellets were re-equilibrated to 37°C for 15 min, followed by pulse-labeling HDX and streptavidin bead binding as described in the subsequent section. Two technical replicates of the pulse-labeling reaction were performed for each sample. The L34 construct was also used to generate undeuterated and maximally deuterated control samples.

Following pulse-HDX, the integrity of the nascent chains (NCs) was assessed by mixing 2 μL of resuspended pellet with 10 μL of low-pH sample buffer for a final composition of 7.5% glycerol, 62.5 mM Bis-Tris pH 5.7, 0.01% bromophenol blue (wt/vol), 0.2% DTT (wt/vol), and 2% SDS (wt/vol) (*65*). These samples were analyzed by SDS-PAGE and western blotting (see section *Anti-biotin Western Blotting*) to ensure homogeneity of each NC construct and to verify a migration pattern approximately 20–25 kDa larger than expected due to peptidyl-tRNA attachment. To generate “released” samples, a separate aliquot of each construct was treated with RNAse A (final concentration of 0.2 mg/mL, Thermo Scientific EN0531) and incubated for 5 min at **37°C** prior to the addition of low-pH loading buffer and SDS-PAGE analysis.

### TEV Elution Experiment

To confirm that streptavidin bead binding does not disrupt nascent chain (NC) integrity, we designed an experiment in which the AviTag is separated from the NC by a linker ending with a TEV cleavage loop; following bead binding, NCs were eluted via the addition of TEV protease. The TEV cleavage loop (ENLYFQ) was inserted at the end of the linker separating the AviTag from the nascent protein of interest (in this case, aTS), and a PCR fragment for *in vitro* transcription and translation was produced as described in subsection *Cloning and PCR*. This PCR fragment was designed to contain a 35 amino acid C-terminal linker downstream of aTS and lacked a stop codon, thereby promoting ribosomal stalling at the end of this linker.

A translation master mix was prepared consisting of:

- 5 μL Solution A –AA, tRNA
- 2.5 ìL of 10 × AA mix
- 2.5 μL tRNA
- 7.5 μL of Solution B-RF123
- 1 μL Murine RNAse inhibitor
- 1 μL of FluoroTect GreenLys
- 1.25 μL of 20 μM BirA
- 270 ng PCR fragment

This master mix was split into three samples (A, B, and C). Samples A and B were supplemented with biotin to a 20 μM final concentration, while sample C was supplemented with an equivalent volume of water. All three samples were incubated at 37°C for 1 h to allow translation to occur. After 1 h, sample B was treated with puromycin to a final concentration of 68 μM to release nascent chains. At this stage, a 2 μL “input” aliquot was drawn from all three samples, mixed with 8 μL of HKMT buffer and low-pH loading dye (to a 1 × working concentration), and kept on ice.

A 6 μL aliquot from each sample was then incubated for 10 s at 37°C with a 54 μL streptavidin bead aliquot pre-equilibrated in D_2_O (see previous sections). After 10 s, the samples were placed on a magnetic stand and the supernatant was removed. The beads were resuspended in 30 μL of a protease mix containing 3 μM TEV protease purified in-house and 1 μL of murine RNAse inhibitor in HKMT buffer, and then incubated for 30 min at room temperature to allow TEV cleavage and nascent chain elution. Samples were placed on a magnetic stand, and 27 μL of the supernatant was recovered and mixed with 9 μL of 4 × low-pH loading dye. All samples were loaded onto a NuPAGE gel, which was run in a cold room (4°C), and the gel was imaged using a Typhoon system as described in *SDS-PAGE and Anti-biotin Western Blotting*.

### SDS-PAGE and Anti-biotin Western Blotting

Samples in 1 × Laemmli sample buffer (for active translation reactions) or low-pH sample buffer (for stalled RNCs or other samples requiring preservation of peptidyl-tRNA integrity) were loaded onto a NuPAGE 4–12%, 1.5 mm, 15-well gel. We included 2 μL of WesternSure Pre-stained Chemiluminescent Protein Ladder (LI-COR, 926-98000) for western blot samples, or 0.5 μL of PageRuler Plus Prestained Protein Ladder (Thermo Scientific 26619) for samples destined for fluorescence imaging. Gels were run at 125–130 V at room temperature, or at 4°C (for samples requiring preservation of peptidyl-tRNA integrity), for at least 1 h in MES running buffer. If gel samples were fluorescently labeled with FluoroTect, the gel was imaged using a Typhoon FLA9500 (GE Healthcare) with a 488 nm laser and a 510LP filter. If labeled with TMR, a 532 nm excitation laser and a 580 nm BP emission filter were used.

Prior to western blot transfer, gels were rinsed in deionized water and equilibrated for 10 min in 1 × Towbin transfer buffer containing 20% methanol, alongside blotting paper and a 0.2 μm polyvinylidene difluoride (PVDF) membrane (Bio-Rad, 1620174) that had been pre-activated in 100% methanol. Transfer was performed at 25 V for 42 min using a Trans-Blot Turbo apparatus. The membrane was then rinsed in deionized water and incubated with 10 mL EveryBlot Blocking Buffer (Bio-Rad, 12010020) on an orbital shaker for 5 min at room temperature, or overnight at 4°C. The blocking buffer was decanted, and the membrane was incubated in a streptavidin solution consisting of 0.5 μL of 2 mg/mL Streptavidin-horseradish peroxidase (HRP) conjugate (Invitrogen S911A) in 10 mL EveryBlot Blocking Buffer for 1 h at room temperature with orbital shaking. The membrane was then washed five times for 5 min each with Tris-buffered saline solution containing Tween-20 (TBST). Prior to chemiluminescent imaging, the membrane was rinsed in deionized water and blotted dry. The blot was imaged via the addition of either 5 mL of each solution of Clarity Western ECL substrate (Bio-Rad 1705061), 6 mL of each solution of SuperSignal West Pico PLUS Chemiluminescent Substrate (Bio-Rad, 34580), or 2 mL of each solution of SuperSignal West Femto substrate (Thermo 34094). The membrane was imaged on a ChemiDoc system. Where indicated, an Alexa Fluor 488 Streptavidin conjugate (Invitrogen A32361) was used in lieu of Streptavidin-HRP, and imaging was carried out using an FLA9500 (GE Healthcare) with a 488 nm laser.

### Pulse HDX-MS, Maximally Deuterated Sample Prep, and Nascent Chain Enrichment

Pierce streptavidin magnetic beads (Thermo Scientific 88816) were equilibrated to room temperature and resuspended, and 60 μL of bead slurry was drawn per translation timepoint. The beads were placed on a magnetic stand to remove the storage solution and subsequently washed with 1 mL of nascent chain labeling buffer (10 mM MES, 30 mM KOAc, 12 mM Mg(OAc)_2_, pH 6.0) in H_2_O. Beads used for deuterated samples were additionally washed in 0.5 mL of nascent chain labeling buffer in prepared in 99.8% D_2_O (TCI W002 or Thermo 426931000). The beads were then resuspended in 54 μL of nascent chain labeling buffer in D_2_O (for deuterated samples) or H_2_O (for undeuterated controls) per timepoint, and 54.3 μL aliquots corresponding to the respective timepoints of interest were transferred into 0.5 mL Protein LoBind Tubes (Eppendorf 022431064).

These aliquots were pre-warmed for 10 min at 37°C prior to use in the pulse HDX-MS reaction. A pD value of 6.0 (as read directly by a pH meter) was chosen for pulse HDX to ensure that a 10 s labeling pulse would label solvent-exposed amides and protect regions that are folded with some level of stability. predominantly label solvent-exposed amides. We calculated that under EX2 conditions, this pH and pulse duration would prevent labeling of any structure formed with a stability greater than approximately 2 kcal/mol at 37°C (*66*).

At indicated elongation timepoints, nascent chains were pulse-labeled and bound to streptavidin beads. This was achieved by drawing 6 μL of elongation reaction (see section *Synchronized Translation Reactions*) and thoroughly mixing it with the respective bead aliquot in deuterated or undeuterated buffer (prepared as described in the previous paragraph). The sample was incubated for 10 s at 37°C to allow hydrogen-deuterium exchange (for deuterated samples) and nascent chain binding to the streptavidin beads, followed by the addition of 60 μL of 2 × ice-cold quench buffer (3.5 M GdmCl, 1.5 M glycine, and 0.5 M TCEP, pH 2.4) and rapid mixing. Samples were then flash-frozen in liquid nitrogen and stored at -80°C until LC-MS injection.

Maximally deuterated control samples were prepared by mixing 6 μL of nascent chains with a 54.3 μL streptavidin bead aliquot pre-equilibrated in D_2_O, followed by a 15 min incubation at room temperature on a rotary mixer to allow complete binding of nascent chains to the beads. The samples were placed on a magnetic stand and the supernatant was removed. The beads were then resuspended in 60 μL of 8 M urea + NC labeling buffer in 90% D_2_O. This buffer was prepared by dissolving urea to 9 M in NC labeling buffer in H_2_O, followed by three rounds of lyophilization and resuspension in 90% D_2_O; the final urea concentration was adjusted to 8 M (as measured by refractometry) using NC labeling buffer in 90% D_2_O. Beads were incubated overnight at 37°C in this deuterated urea buffer, followed by the addition of 60 μL of ice-cold 2 × quench buffer. Samples were then flash-frozen in liquid nitrogen and stored at - 80°C until LC-MS injection. For the first CMPK biological replicate, maximally deuterated samples were incubated at 37°C for 3 h rather than overnight; we confirmed this duration is sufficient to produce maximal deuteration. Maximally deuterated controls were prepared in duplicate for each sample.

Prior to LC-MS injection, samples of all types were thawed by rapid flicking and passed through a Corning Costar Spin-X 0.22 μm filter (Costar 8161) by centrifuging for 30 s at 12,000 × *g* to separate the quenched labeling solution from the beads, which are retained in the filter basket. The beads were then resuspended in 1 × quench buffer (2 × quench buffer diluted in NC labeling buffer in H_2_O) containing 0.2 mg/mL of porcine pepsin (Sigma P6887) to initiate digestion. A total digestion time of 4 min was found to yield optimal peptide coverage for HaloTag samples, while 5.5 min was found to be optimal for CMPK and aTS samples.Immediately after resuspension in the pepsin-containing 1 × quench, the beads were vortexed for 5 s and placed on ice. Two minutes prior to the end of digestion, the bead-containing centrifugal filter was centrifuged again for 30 s at 12,000 × *g*, and the flow-through containing the peptides was collected in a clean 1.5 mL LowBind tube and returned to ice until the end of the digestion period. At this point, the liquid sample was injected into the LC-MS system.

### LC-MS Runs

Peptide-containing samples were injected into a cooled valve system (Trajan LEAP) in line with a Thermo Ultimate 3000 LC. Peptides were desalted on a C8 pre-column (Waters 186003978) for 4 min, and subsequently bound to a Waters Acquity C18 analytical column (Waters 86002344); both columns were maintained at 2°C. The bound peptides were gradient-eluted (5–40% acetonitrile w/v and 0.1% w/v formic acid) across the analytical column at a flow rate of 40 μL/min over 14 min, followed by a gradient from 40% to 90% buffer B over 30 s. The analytical and trap columns were then sawtooth-washed and equilibrated at 5% buffer B. Eluted peptides were analyzed directly using a high-resolution Orbitrap mass spectrometer (Q Exactive, Thermo Fisher) operating in positive mode [resolution: 140,000; automatic gain control (AGC) target: 3 × 10^6^; maximum injection time: 200 ms; scan range: 300 to 1500 mass/charge ratio (*m*/*z*)]. For each construct, we performed a tandem mass spectrometry (MS/MS) experiment on an undeuterated sample [full MS resolution: 70,000; AGC target: 3 × 10^6^; maximum injection time: 50 ms; scan range: 300 to 1500 *m*/*z*; dd-MS2 settings: resolution: 17,500; AGC target: 2 × 10^5^; maximum injection time: 100 ms; loop count: 10; isolation window: 2.0 *m*/*z*; normalized collision energy: 28; charge states 1 and ≥ 8 excluded; dynamic exclusion: 15 s].

Between injections, the sample loop and trap column were manually washed at 200 μL/min with 50% acetonitrile + 0.1% formic acid for 4 min, followed by one wash of the protease column with 100 μL of 1.6 M GdmCl and 0.1% formic acid, and three sawtooth washes of the trap and analytical columns (5–90% acetonitrile gradient each).

### Mass Spectrometry Data Analysis

Our basic data analysis pipeline involves the following steps:

1. **Peptide identification:** Observed MS1 and MS2 spectra from undeuterated control samples are assigned to peptides derived either from the protein of interest or from other protein components in the translation and enrichment pipeline.
2. **Manual inspection and curation** of peptide mass spectra from deuterated samples.
3. **Fitting of mass spectra** to multimodal distributions and identification of the optimal number of modes.
4. **Bootstrapping** error analysis.

### Peptide Identification

RAW files from undeuterated control injections, containing precursor and fragment mass spectra (MS1 and MS2, respectively) across all retention times, are loaded into Byonic (Protein Metrics) software. For peptide identification, we input a FASTA file containing sequences for all *in vitro* translation components present in PUREexpress reactions, as well as protein components from the nascent chain enrichment steps (namely, streptavidin and pepsin) and the translated protein of interest downstream of the AviTag and its respective linker.

The following peptide search parameters are used:

- Precursor mass tolerance: 6 ppm
- Fragmentation type: QTOF/HCD
- Fragment mass tolerance: 20 ppm
- Digestion specificity: Non-specific
- Maximum precursor mass: 10,000
- Number of precursors per MS2: 1
- Smoothing width (*m*/*z*): 0.01

We retain all peptide and charge state identifications associated with a Pep2D score <0.1, and export these spectra and associated retention times for subsequent analysis.

### Manual Inspection of Peptide Spectra

RAW files corresponding to all deuterated samples are loaded into HDExaminer 3 (Sierra Analytics) software, alongside the peptide lists and associated retention times from the previous step. Using HDExaminer, we manually inspect deuterated mass spectra assignments for all peptides to ensure that intensity peaks are centered about their expected *m*/*z* isotopic windows, and that the retention times at which the peptide spectra are observed differ from those in undeuterated spectra by no more than ∼ 0.2 min. If any retention time shifts are observed for a given peptide, we verify that such a shift is systematically observed across all peptides within that sample.

We additionally adjust the integrated retention time windows to ensure that these integration ranges are comparable across samples for each specific peptide. As an additional quality control step, we examine the corresponding *m*/*z* and retention time ranges in a negative control sample for each peptide to rule out interfering or carry-over signals. For each sample, we discard any peptide spectra for which the negative control signal in relevant *m*/*z* windows (namely, all isotopic windows ranging from the monoisotopic peak to the largest meaningful window containing signal in the maximally deuterated control) is more than one-fifth of the signal within the sample. This threshold indicates that the background is expected to contribute meaningfully to the observed signal. This criterion excludes interference from both background peptides whose spectra overlap with the target peptide and carryover; the latter is observed only in a small fraction of peptides. Following manual curation, all resulting high-confidence peptide mass spectra are exported and fit as described below. These high-quality peptides are shown as colored vertical lines in the coverage maps for each protein construct in Fig. S2.

### Peptide Spectra Fitting

During co-translational protein folding, we expect different structural regions to cooperatively fold and attain protection from deuterium exchange in a time-dependent manner. Because the pulse timescale (10 secs) is significantly shorter than the timescale of translation, we obtain a snapshot of the conformations populated at the time of the pulse. These conformations have different protection patterns, manifesting as peptide mass spectra with distinct peaks representing their respective level of protection.

For each peptide, we globally fit all mass spectra from distinct co-translational folding timepoints to a combination of basis spectra indexed by *m* corresponding to these distinct conformational states. We assume that the centroids themselves are constant across timepoints but allow the relative contribution of each basis (its weight) to vary. That is, for a peptide with charge state *z*, we assume the mass spectral intensity at mass-to-charge value *m*/*z* and translation time *t* is given by:

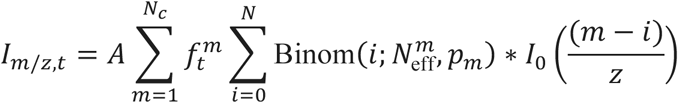

Where *A* describes the signal amplitude related to factors such as digestion and ionization efficiency, *N_c_* is the number of curves being fit, *f^m^* is the fraction of signal associated with mode *m* at time *t* (at each timepoint, these must sum to 1 over all *m*), *N* is the total number of exchangeable amides in the peptide, Binom(*i*; *N^m^*, *p_m_*) refers to a normalized binomial distribution evaluated at integer *i* with parameters *N^m^* and *p_m_* (the product of which is the average number of deuterons taken up for mode *m*), and *I*_0_(*m*/*z*) is the mass spectral intensity for the natural abundance isotopic distribution for the undeuterated peptide at m/z. In this formulation, we neglect the instrument-induced mass spectrometric peak width as well as any interference from other overlapping peptides.

To analyze data for a given peptide, we globally fit the above equation to the mass spectrometric signal at all isotopic peaks across all timepoints, optimized for parameters *A*, 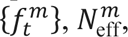, and *p_m_*. The 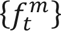 parameters reveal the population fractions associated with each mode at each translation time, while the product 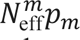 gives the centroid for mode *m*. Where technical replicates exist for a given condition (in the case of stalled RNCs), we required that both replicates be fit to the same parameters corresponding to that respective condition. Fitting is implemented in custom Python scripts using the SciPy package. Our approach and fitting algorithm, including our calculation of the natural isotopic distribution, are largely based on those in reference (*67*), but adapted to allow for global fitting across timepoints.

To decide on an optimal number of curves *N_c_*, we balance attaining a high-quality fit with the risk of overfitting. This is achieved in two steps. First, we compute the Bayesian information criterion (BIC) associated with the fit for each *N_c_* value ranging from 1 to 3, which we approximate as:

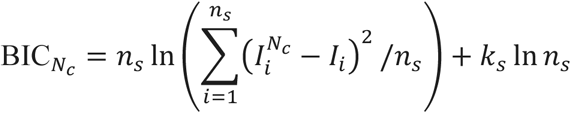

Where the numerator in the first logarithm is the residual sum of squares, obtained by summing the squared differences between the model fit 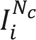 (given *N_c_* curves) and the experimental peak intensities *I_i_* over data points indexed by *i* spanning the total number of data points for the peptide (*n_s_*), and *k_s_* is the total number of model parameters (excluding the final fraction for each timepoint whose value is constrained by normalization). We note that intensities are normalized per-spectrum prior to being plugged into the above calculation so that timepoints with higher ion abundance are not weighted more heavily. We then determine the smallest number of curves *N_B_* for which the BIC(*N_B_*) is greater than the smallest BIC value (across all *N_c_* ranging from 1 to 3) by no more than 10 units. As a second method, we compute the goodness of fit for each configuration as:

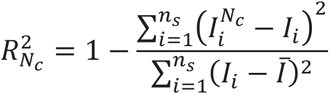

Where *̅I* is the average intensity over per-timepoint normalized spectra. From these values, we determine *N_R_*, which is the smallest value of *N_c_* required to achieve 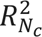 > 0.95. Finally, the optimal number of curves is chosen as:

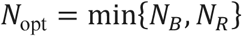

These combined criteria prevent overfitting in a few cases where the BIC calculation favors a larger number of curves than is needed to achieve a satisfactory fit. In the SI, we show plots of selected spectra fit to various numbers of curves with an indication of the optimal number per the above criteria, as well as the mathematical justification. In Fig. S5, we show fits for three representative peptides from aTS to one, two, or three curves as visual evidence that this approach yields a sensible *N*_opt_ value. The above criteria are applied to the majority of peptides and overwritten manually in a limited number of cases, primarily where interfering peaks from other peptides are spuriously fit by the algorithm—in this case, the values obtained by these calculations are not reliable. Maximally deuterated samples, in contrast, are fit to just one curve in all cases.

### Bootstrapping and Centroid Comparison

To obtain error estimates on fitting parameters, we bootstrap by resampling global fit residuals (relative to the fit of the initial data) with replacement. Residuals are freely shuffled across spectra from different timepoints and technical replicates (where present) for a given peptide; however, we only allow residual exchange between a given pair of data points if their respective intensities differ by no more than a factor of 5, accounting for the possibility that the noise distribution differs across intensity scales. We also enforce positivity for all resampled intensities.

These resampled data points are then globally re-fit to the same number of curves determined as per the previous section, and the resulting parameters are recorded. This process is repeated 2,000 times for each peptide to obtain distributions and confidence intervals for all parameters, including the number of deuterons for each mode *m*, computed for each bootstrap iteration as the product 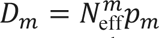. We discard any bootstrap iterations for which *D_m_* differs from the median value by more than three times the median absolute deviation, as these often represent scenarios where the fit is distorted by spurious peaks due to interfering peptides. We refer to the resulting bootstrapped distribution for the number of deuterons in mode *m* as *P*(*D_m_*). Separately, we bootstrap the two maximally deuterated control technical replicates for each sample to obtain a parameter distribution for that condition.

For certain analyses, we compare the deuteration of the least protected (highest *D_m_*) mode with *D_m_*_,max_, the number of deuterons in the unfolded state under the null hypothesis that the former is no smaller than the latter. Extracting *D_m_*_,*u*_ from our maximally deuterated control sample requires correcting for the fact that this sample was prepared via overnight incubation in 90% D_2_O, whereas pulse HDX-MS samples are deuterated for only 10 s. This cannot be trivially corrected experimentally because the presence of 8 M urea slows down the rate of exchange in the base-catalyzed regime (*68*). At a meter-read pH of 6, we find this leads to incomplete labeling of denatured proteins within 10 s. Instead, we calculate the fraction of deuterons that we expect to have exchanged during our 10 s pulse for each peptide and scale the number of deuterons taken up in our maximally deuterated control by this fraction, as a back exchange correction.

To achieve this, we use a Python implementation of the algorithm in reference (3) to compute the expected exchange rates *k_i_* for all backbone amides across the intact protein sequence. These calculations are performed assuming a temperature of 310 K and pH 5.6.

For a given peptide spanning residues indexed from *a* through *b*, we define the corrected exchange rates as follows:

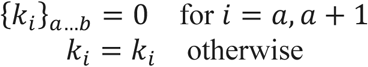

where the first two peptide residues have an effective exchange rate of zero because they back-exchange very quickly following digestion (the first residue, in fact, contains a free amine rather than an amide). We then compute the corrected deuteration for the unfolded state as:

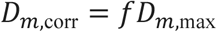

where:

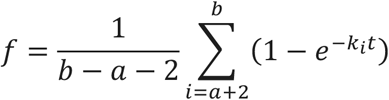

Where *t* = 10 s, assuming single-exponential exchange kinetics for each site, and *D_m_*_,max_ is the number of deuterons exchanged in the maximally deuterated control. For most peptides, we obtain *f* values in the range of ∼ 0.9 to 1. We likewise apply this correction to the centroid obtained for all bootstrap iterations performed on the maximally deuterated control to obtain a corrected *P*G*D_m_*_,corr_H for the expected deuteration of an unfolded polypeptide following 10 s of pulse labeling under our experimental conditions. Finally, we compute the p-value associated with the null hypothesis that *D_m_* ≥ *D_m_*_,corr_ as the fraction of bootstrap trials for which the observed centroid is at least as large as the corrected maximally deuterated centroid as an approximation to:

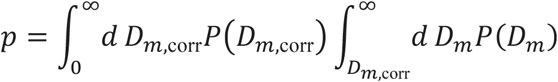

For any peptides for which *p* < 0.01 for the least protected mode and for which *D_m_*_,corr_ − *D_m_* ≥ 0.5, we reject the null hypothesis and conclude that the least protected mode is significantly more protected than would be expected if it reflected the unfolded state. This same definition of statistically significant burst-phase protection is also used to designate which peptides are shown with stripped shading in Fig. 2H, with the additional requirement that this partially protected mode show no more than 85% deuterium uptake in this specific panel. For fig. S9E which shows mode deuterations normalized by the maximally deuterated value, the error bars are obtained via element-wise divided samples from the bootstrapped distribution for the respective mode and that for the corrected maximally deuterated control, respectively and then extracting the 95% confidence intervals from the resulting distributions.

We note that, in main text figures showing fractional populations as a function of time and mode protection at each peptide, a subset of high-quality peptides showing poor global fits is excluded. In addition, in cases where multiple charge states exist for a given peptide, the main text figures present only the charge state showing the stronger signal and/or higher quality fitting. In the supplementary figures, we present these graphs with all high-quality peptides and redundant charge states included. We observe reproducible behavior across charge states for a given peptide, even though these are fit and analyzed independently (see supplementary figures).

### End of Translation Determination

As a lower-bound estimate for the first timepoint associated with the end of translation, we considered, for each timepoint *t*, the most intense amplitude *A_C_*_max,*t*_ associated with any peptide/charge state combination spanning the most C-terminal residue for which we have coverage, as well as the least intense amplitude *A_N_*_min,*t*_ associated with any peptide spanning the most N-terminal residue for which we have coverage, and computed their ratio, *R*_CN,*t*_. We then normalized by this intensity ratio for the exact same peptide/charge state pair at the final translation timepoint, *R*_CN,*t*final_, and determined the first timepoint *t* for which:

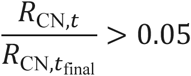

This threshold implies that enough ribosomes have completed translation that full length protein will appreciably contribute to the detectable mass spectrometric signal at N-terminal peptides. For all proteins, we additionally verified that this timepoint coincides with the appearance of appreciable full-length protein by Western blotting.

### Supplementary text

Supplementary Note 1: Evidence that our method detects folding on the ribosome

The control experiments shown in fig. S1D indicate that nascent chain (NC) binding to streptavidin beads under HDX conditions does not disrupt the integrity of NCs. But this experiment does not rule out detection bias in favor of released proteins resulting from subsequent steps in our sample preparation and injection pipeline, including quench or digestion-related biases. Here, we provide two lines of evidence pointing against such a bias. Together, these strongly indicate that our MS signal faithfully reports on NCs at timepoints when translation is still occurring.

### Evidence 1: Mass spectrometric intensities in CMPK are inconsistent with strong bias in favor of released proteins

Let us consider the alternative hypothesis that our mass spectrometric signal is exclusively sensitive to released proteins, namely either nascent chains that are released prematurely or fully synthesized, released protein. For CMPK, we observe that roughly ∼70% of ribosomes finish translation, as obtained by quantifying the intensity of the full-length band, relative to the total intensity in the 15 min lane in fig. S4E. Thus, under our **alternative hypothesis, the maximum mass spectrometric signal we will be able to detect from nascent chain peptides at co-translational timepoints will be ∼30% that at post-translational times**, assuming 1) negligible variability in total injected material (see next paragraph) and 2) all peptides whose signal is compared have been fully synthesized by 10 mins; this is likely true for N-terminal peptides on which we focus our attention.

In reality, however, we consistently observe that these two conditions show comparable signal at N-terminal peptides (Table S1). For example, for the core peptide 2-24, z=3, the overall intensities (quantified as the total amplitude associated with the global fits at the respective timepoints) at the 10 and 15 min timepoints nearly identical at 1.8E8 and 1.7E8, respectively.

The 10 and 15 min intensities are likewise nearly identical for the lid peptide 40-45 z=2, at 1.2E8 and 1.1E8, respectively. We note that this result holds true when corrected for injection-to-injection variability in total amount of injected material. As a proxy this total injected material, we quantify the intensities associated with peptides from two distinct ribosomal proteins, L5 and L23, as detected in these specific samples (although our biotin-streptavidin pulldown strategy appreciably enriches for the NC, some signal from ribosomal proteins is nonetheless detected due to nonspecific binding to streptavidin beads and/or co-association with NCs stemming from incomplete ribosomal dissociation during quench).

Although these analyses reveal that the amount of material in the 10 mins sample is slightly greater than that in the 15 mins sample by a factor of 1.5 using the proxy ribosomal protein L5 (Table S2) and a factor of 1.4 when using the proxy protein L23 (Table S3), these ratios are not sufficient to support the alternative hypothesis; the total injected material in the 10 min timepoint would have to be at least ∼3.3 times that in the 15 min timepoint in order for the two timepoints to show similar N-terminal CMPK intensities under the hypothesis that no more than 30% of the population (i.e. the spuriously released NCs) is detectable at the 10 min timepoint.

We additionally note that for lid peptide 40-45, ∼60% of this signal at 10 min is attributed to the protected mode, and the remaining 40% is attributed to a burst phase mode with significant protection (Fig. 2C and fig. S4). Let us consider the related hypothesis that this ∼60% detectable protected population consists entirely of spuriously released NCs, which comprise up to 30% of the population. Then this would require that the total amount of injected material associated with the 10 mins timepoint be at least ∼2x associated with the 15 mins sample, again larger than the **∼1.4x-1.5x factor observed.** This strongly supports that our method can detect co-translational folding in actively translating nascent chains.

We note that over long timescales (∼30 min to hours), the signal at all peptides, including N-terminal peptides steadily increases, which we attribute to multiphasic biotinylation kinetics (fig. S1G).

### Evidence 2: Mass spectrometric signal is unlikely to be skewed by nascent-chain-specific precipitation during HDX quench

As a second line of reasoning, we explicitly consider what we believe to be the most likely mechanistic origin for any possible detection bias against nascent chains: namely, selective precipitation of peptidyl-tRNA attached NCs during low-pH quench conditions (*52*), which could inhibit digestion of these species while potentially leaving released proteins accessible to digestion. As our quench step is applied to NCs that are physically attached to micron-sized streptavidin beads, we cannot simply perform a centrifugation-based pelleting assay to quantify the amount of precipitated NCs under our conditions. Instead, we address this hypothesis by systematically increasing the concentration of GdmCl in our quench buffer from 2M (the concentration used in most of our experiments) to 6M which should significantly reduce precipitation. Thus, if the signal under our standard 2M quench is heavily biased by aggregates, then this [GdmCl] increase should notably alter the mass spectra we observe. We note that in both cases we reduce the [GdmCl] to 2M while adding pepsin to ensure the protease can efficiently digest the sample—this complete buffer exchange is uniquely enabled by our bead setup.

Contrary to this expectation, we observe highly similar mass spectra for the model protein aTS between two simultaneously-performed pulse-labeling HDX replicates after 8 mins of synchronized elongation—these replicates differ only in their concentration of GdmCl in the quench as described above. For example, under both quench conditions, peptides 1-7 and 128- 152 show no protection at 8 mins, whereas peptides 35-40 and 85-101 show partial protection. As controls, we also confirm that the 60 mins (post-translational) and maximally deuterated conditions also show similar spectra across both quenches with slightly increased back-exchange when 6M GdmCl is used. These results strongly suggest that our signal is not notably biased against NCs due to acid-induced precipitation under our standard quench conditions.

We note that, formally, we cannot rule out the possibility that some on-bead aggregation occurs amongst bead-attached NCs in our 6M sample following buffer-exchange into the 2M buffer to enable digestion. However, this is extremely unlikely under single-turnover translation conditions where the maximal NC concentration equals the ribosome concentration, which is about 2 µM in PURE systems (*46*). We estimate that under our experimental conditions assuming this upper-bound concentration, the average spacing between NCs on the beads would be about 16 nm. This spatial isolation strictly constraints macro-aggregation, limiting any possible interactions to low-order oligomerization at most. Moreover, the concentration of NCs is almost certainly much less than 2 µM in our experiments, as per our observation that the total intensity due to translated protein on a Coomassie-stained gel is significantly less than that associated with ribosomal proteins (see fig. S7C for such a gel on HaloTag).

**Table S1.**
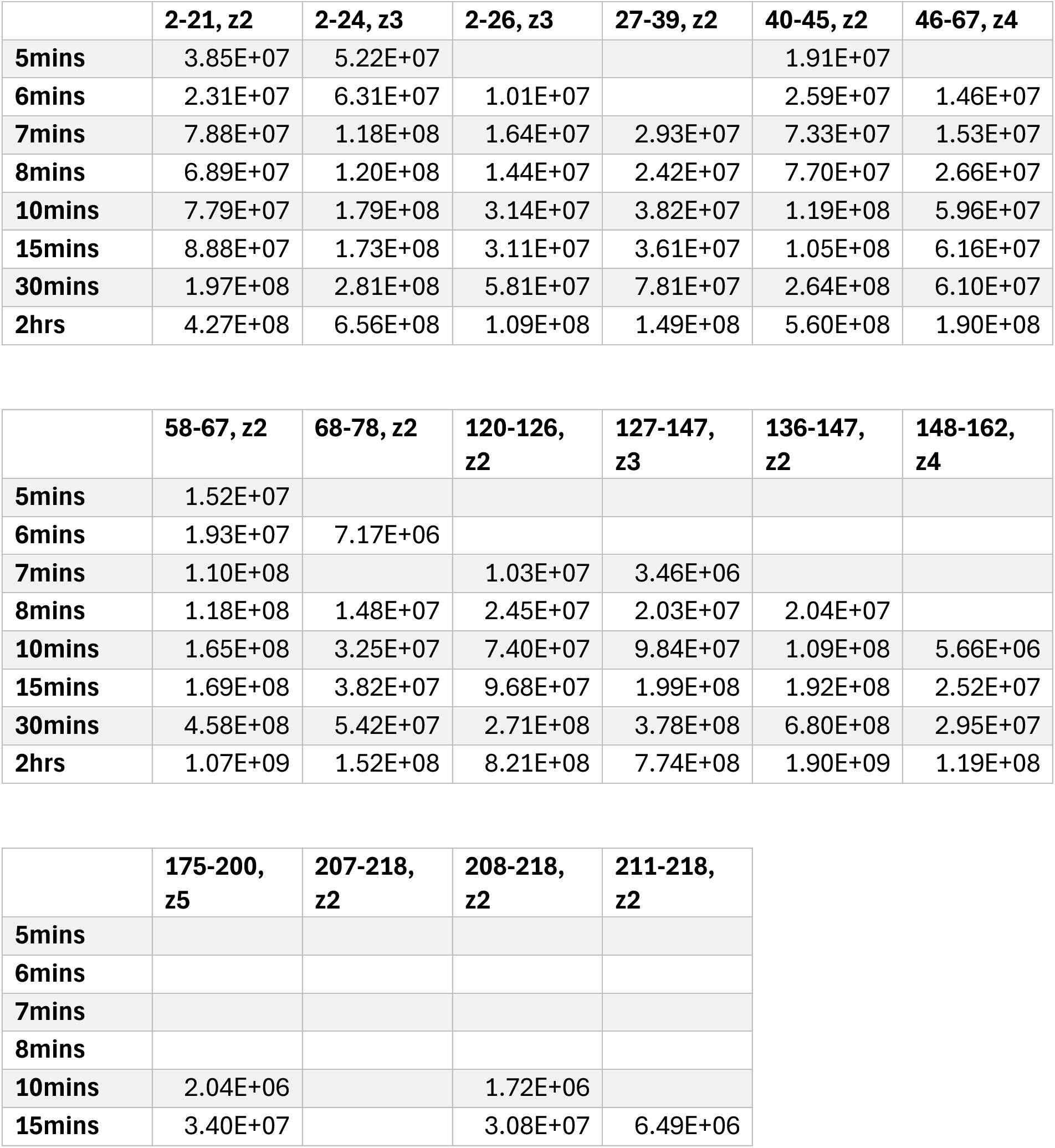
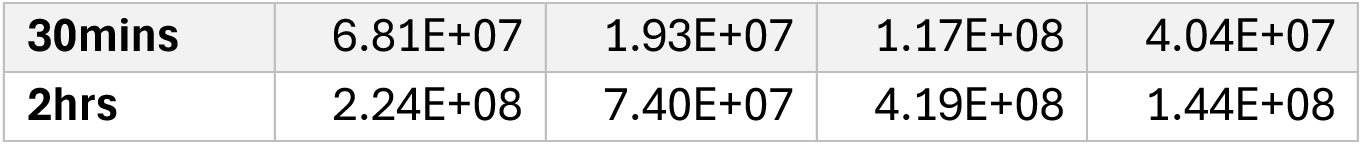
Global fit amplitude, a quantification of overall mass spectrometric intensity, as a function of translation time for spectra corresponding to indicated CMPK peptide and charge state combinations. No value indicates the peptide was not detected at the indicated time

**Table S2.**
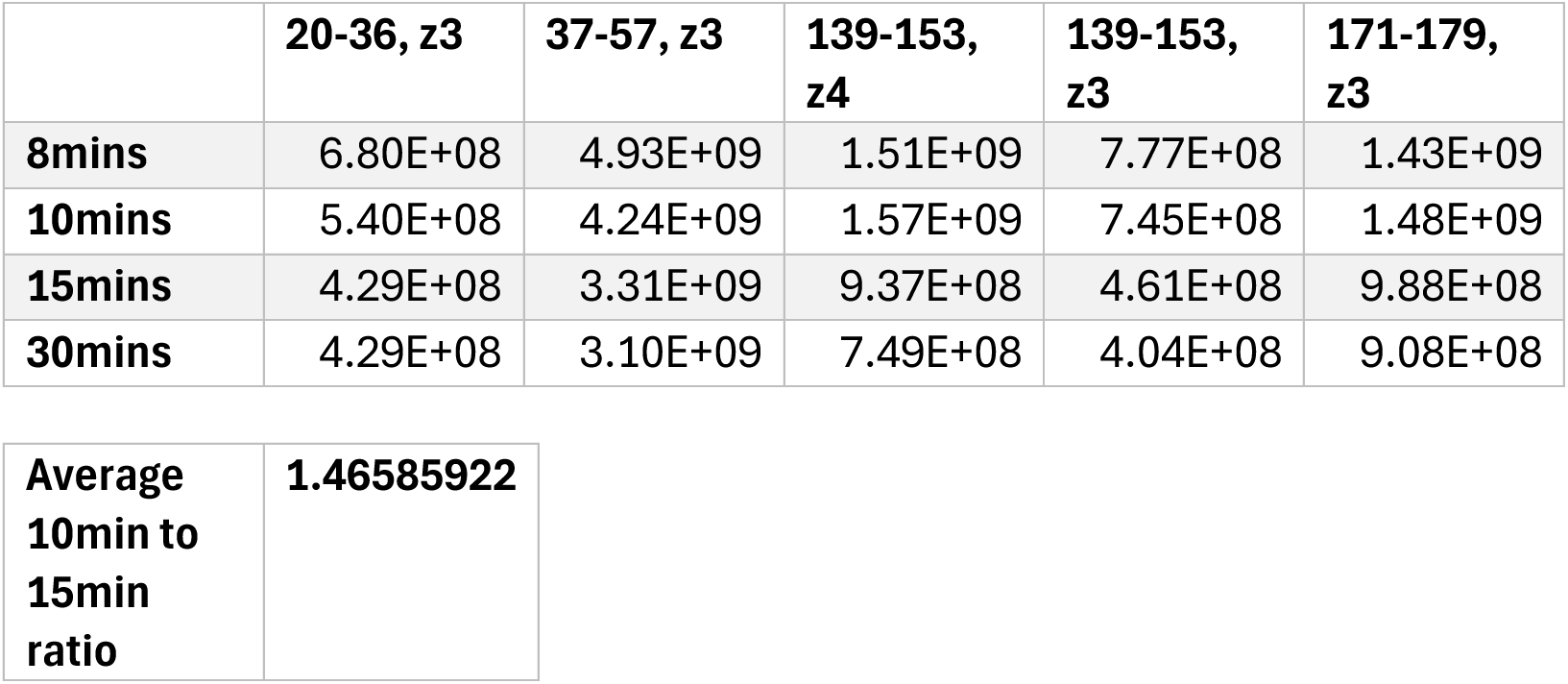
Global fit amplitude, a quantification of overall mass spectrometric intensity, as a function of translation time for spectra corresponding to indicated peptide and charge state combinations from ribosomal protein L5 during translation reaction of CMPK. Only a subset of translation times is shown. Final row shows average ratio of intensity at 10 mins to that at 15 mins over all peptides.

|  |  |
| --- | --- |
| **Average 10min to 15min ratio** | **1.46585922** |

**Table S3.**
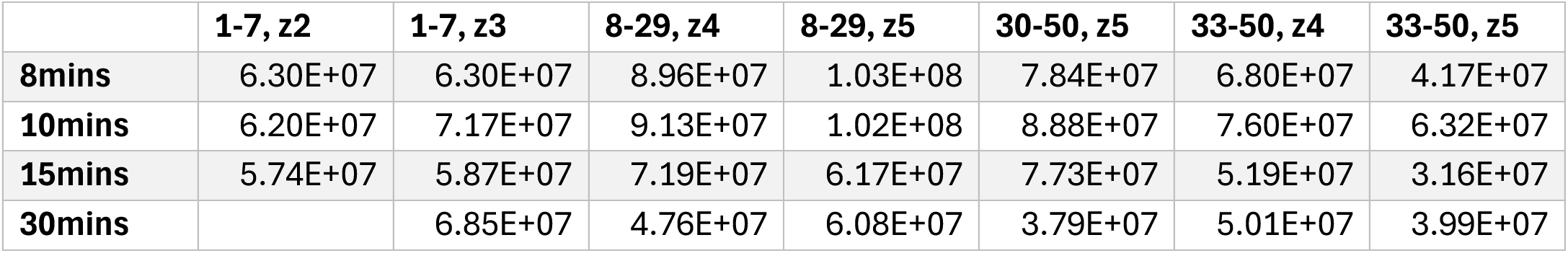
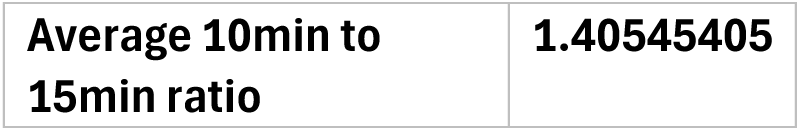
Global fit amplitude, a quantification of overall mass spectrometric intensity, as a function of translation time for spectra corresponding to indicated peptide and charge state combinations from ribosomal protein L23 during translation reaction of CMPK. Only a subset of translation times are shown. Final row shows average ratio of intensity at 10 mins to that at 15 mins over all peptides.

Supplementary Note 2: Reproducibility analysis

We analyze reproducibility in the order in which peptides become protected across biological replicates of our translation reactions by generating scatter plots showing the natively protected fractional population at a given timepoint in both replicates and computing the spearman correlation. These analyses are conducted at timepoints at which a subset of peptides has begun to show protection, while others do not, so as to most cleanly delineate the sequence of acquisition of protection. These plots are shown in fig. S4I-J, fig. S6H, and fig. S7H-I for CMPK, aTS, and HaloTag, respectively at the indicated timepoints. These analyses all show high Spearman correlation coefficients above 0.8 with associated p values less than 10^-5^, except for CMPK which has a reduced peptide coverage relative to the other proteins and p-values of approximately 10^-4^ to10^-3^. The following factors contribute to the slight remaining variability between replicates:

1. As the two replicates are fit independently, the centroids associated with the relevant modes may vary slightly between replicates. As a result, the associated population fractions at a given timepoint may vary. This is especially applicable for peptides exhibiting closely overlapping modes. This may be further confounded by the fact that, in some replicate reactions, certain timepoint were not collected.
2. Variations in translation speed imply that, at a given timepoint, different replicates may have progressed to different degrees in the folding reaction. This will lead to noise in the fitted population fractions, particularly when the fractions are close to 0 or 1 in one replicate, but not the other.

The observation of high and statistically significant correlation coefficients despite these limitations in the analysis strongly indicates a high degree of reproducibility.

Notably, applying this comparison of HaloTag co-translational folding and refolding at comparable timepoints did not yield a meaningful correlation (R=-0.21, p=0.55, fig. S8E), supporting that these proceed through different folding pathways.

Supplementary note 3: Altered HaloTag folding sequence cannot be accounted for by lagging ribosomes

We note that peptides from N-terminal regions will generally appear to fold more slowly in our assay than peptides from the C-terminus, even if the regions fold simultaneously in reality, as lagging ribosomes will present as a population of unfolded N-terminal peptides that is slow to vanish. However, this effect is not sufficient to account for our observation that newly-translated HaloTag folds starting from the C terminus, in contrast to refolding from denaturant where initial folding involves both termini. To show this, we consider the null hypothesis in which the folding sequence of newly-synthesized HaloTag is precisely the same as that observed during refolding, and show that even after accounting for the apparent delay in N-terminal folding due to lagging ribosomes, this model cannot explain our observed results .

To do this, we develop an analytical null model under minimal assumptions and set upper bounds on the ratio of the probability that C-terminal peptides appear folded to the probability that N-terminal peptides appear folded at various times under our model. We show that the upper bound on this ratio under our null model is lower than many of the values we observe.

### Null model assumptions

1. We assume native folding of all regions in a given HaloTag molecule does not begin until that molecule is fully synthesized and released from the ribosome, based on our observation of no native-like co-translational folding.
2. We let *f* denote the fraction of ribosomes that successfully finish translation, conditional on them having translated through the N-terminal residues of interest, and assume that the remaining fraction 1 *— f* of aborted nascent chains do not show any folding of residues that have been synthesized. In reality, our stalled RNC experiments demonstrate that these may fold given enough time, but allowing for this will only decrease the worst-case upper bound that our null model predicts for the C-terminal to N-terminal folded ratio.
3. While folding of neither N- nor C-terminal regions is permitted until after the C-terminus is complete, we assume that N-terminal peptides can be *detected* (in the unfolded state) immediately upon their translation.

### 1 The model

We define the following general functions:

- *g_N_* (*t*): The probability density function that a ribosome finishes translating the N-terminal residues of interest at time *t*.
- *g_C_*(*t*): The probability density function that a ribosome finishes translating the full protein at time *t*. Note that *g_C_*(*t*) *≤ g_N_* (*t*) for all *t*.
- *f_N_* (*t*): The probability that an N-terminal region of interest is folded in a given HaloTag molecule some time *t* after that molecule’s translation is complete.
- *f_C_*(*t*): The probability that a C-terminal region of interest is folded in a given HaloTag molecule some time *t* after that molecule’s translation is complete.

To begin, we consider pairs of N- and C-terminal regions where the former folds more rapidly than the latter during refolding from denaturant. Thus under our null assumption, *f_N_* (*t*) *≥ f_C_*(*t*) for all *t*.

Under these assumptions, the observed probability that an N-terminal region and a C-terminal region of interest appear folded at time *t* are given by, respectively:

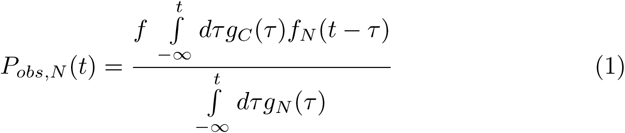

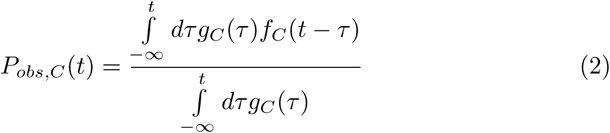

Using *f_N_* (*t*) *≥ f_C_*(*t*), we then have:

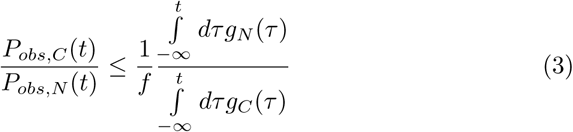

Let us further assume that all ribosomes complete synthesis of the N-terminal region before any ribosome finishes translation of the full protein, in which case the integral in the numerator of Equation 3 is 1 for all *t* of interest. Although this assumption may not be true in general, making this assumption allows us to set the strictest possible upper bound on *P_obs,C_*(*t*)*/P_obs,N_* (*t*). In this case, we have:

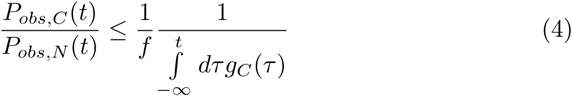

Or inverting, we have

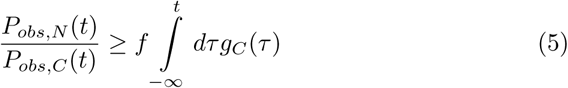

In other words, the observed ratio of the N-terminal folded fraction to the C-terminal folded fraction must be at least as large as the total fraction of ribosomes that has finished translated at time t.

### Null model predictions applied to experiment

We now show that *P_obs,N_* (*t*)*/P_obs,V_* (*t*) is smaller than allowed by Equation 5 at various timepoints *t*, in violation of our null model. In all cases we use a value of *f* = 0.7, under the assumption that all protein regions that have not folded at the final (240 min) timepoint come from aborted ribosomes—this yields an aborted fraction of 0.3. In reality, this assumption may be too strict as this unfolded fraction may also include a subset of fully synthesized molecules that fails to fold. We further reiterate that this bound reflects the extreme scenario where N and C termini fold simultaneously. In the following section (”Null model under additional assumptions”), we explicitly vary the ratio of folding rates and, under additional assumptions, can obtain a looser bound.

### 20 minute timepoint

At this time, the intensity ratio of the most intense C-terminal peptide to the least intense N-terminal peptide, normalized by this same ratio at the final timepoint, is *⇠* 0.65, indicating that approximately this fraction has finished translating. Notably, quantification of the 20 min lane in the western blot in Fig. S7B gives an even higher fraction of 0.8 (which itself may be an underestimate due to saturation of the full-length band). Using the more conservative value of 0.65, Equation 5 gives:

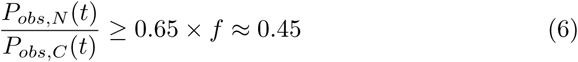

Again using *f* = 0.7. At the 20 min timepoint, the C-terminal peptide 274–285 is already 100% folded, so all regions that fold more rapidly than this C-terminal region during refolding must be at least 45% folded at this time under our null model. But as shown in Fig. 3C, many such peptides are notably less folded than this. For example, 19–36 is only about 30% folded. Peptide 2–18 is less than 20% folded—this is true of both charge states detected for this peptide (Fig. S7). Finally, peptide 81–87 is only about 35% folded (We note that during refolding, this region is reported on by an analogous peptide 81–90). Similarly, at 20 mins, peptide 252–264 is approximately 65% folded. Thus, all peptides that fold more rapidly than 252–264 during refolding must be at least 29.5% folded per our null model. But this is again violated by 2–18.

We note that even in the unlikely case that both our methods for quantifying the amount of fully translated protein yielded a significant overestimate and the true fully translated fraction were as low as *⇠* 30% (a highly unrealistic scenario given the substantial intensity for full length band on western blot and the high intensity for C-terminal peptides that matches expectation from the final timepoint assuming synthesis is complete), then the observed folded fraction for peptide 2–18 would still be too low in comparison with 274–285 to be accounted for by our null model, which would set a bound at 21% in this extreme case.

### 60 minute timepoint

At this time, the intensity ratio of the most intense C-terminal peptide to the least intense N-terminal peptide, normalized by this same ratio at the final timepoint, is *⇠* 1.14 times that at the final (240 min) timepoint, a strong indication that essentially all ribosomes have finished translation. At these times, the integral in the denominator of Equation 4 is simply 1, and so we have:

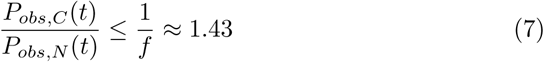

But as shown in Fig. 3C, if we consider the C-terminal peptide 274–285 and the N-terminal peptide 2–18, we observe a *P_obs,C_*(*t*)*/P_obs,N_* (*t*) ratio of approximately 2 at 60 mins, notably higher than this bound, despite the fact that during refolding, 2–18 folds more rapidly.

### A note on 274–285 vs 286–293

Let us now consider peptide 274–285 as the “N-terminal peptide” and 286–293 as the “C-terminal peptide”. We note that at all timepoints prior to the end of refolding, the analogous 286–294 peptide is *more* folded than 274–285, so for this peptide pair, the inequality in Equation 3 flips:

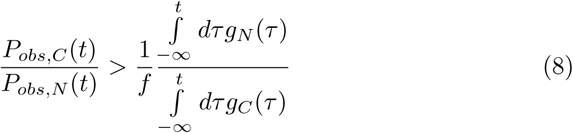

Since these peptides are adjacent, it is reasonable to assume that few ribosomes stall between them, so it is likely that f *≈* 1. Moreover, using the fact that the numerator in Equation 8 is greater than the denominator, we would thus expect:

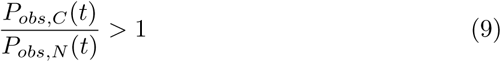

In reality, the expected lower bound would be much higher than this since 274–285 folds significantly more slowly than 286–293 during refolding. But even neglecting this, at all timepoints in our translation experiment, this ratio is actually 1, in violation of our null model.

### Null model under additional assumptions

Under a few additional reasonable assumptions, we can set even more strict bounds on the ratio of folded fractions that is expected under the null model at times shortly after the first ribosomes have completed synthesis. At these early times, for example, the 15 mins translation timepoint in the main-text replicate, it is reasonable to assume single-exponential folding kinetics, as the folding times will be much shorter than the characteristic timescales associated with any slower exponential rates in a multi-exponential process. Thus,

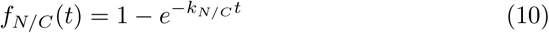

where k_N/C_ denote rate constants for N and C terminal regions, respectively. For analytical tractability, we further assume a uniform distribution of synthesis-completion times of width ⌧_f_ . That is:

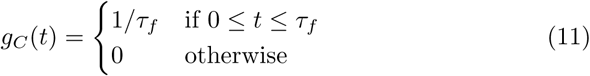

Under these conditions, we have:

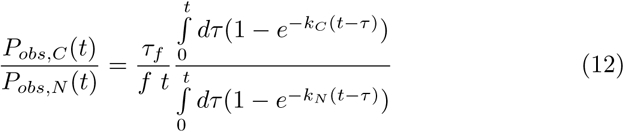

for 0 *≤* t *≤* ⌧_f_ . The integrals can be evluated and the result expressed in terms of dimensionless quantities:

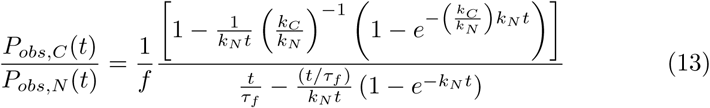

Plots of Equation 13 are shown in Fig. S7J as a function of *k_C_/k_N_* . We expect a value of approximately 0.1 from Dall et al., PNAS 2024. We further note that at 15 mins, the intensity ratio of the most intense C-terminal peptide to the least intense N-terminal peptide, normalized to this same quantity at the final timepoint, is roughly 0.4, so we set *t/*⌧*_f_* = 0.4, although we consider a range of values. Finally, we note that in our translation experiment, N-terminal peptides fold at timescales slower than 30 mins, and that at the 15 min timepoint, the fastest ribosome must have finished translation less than 5 minutes ago (since no C-terminal peptides were observed at the 10 min timepoint). We thus insist that *k_N_ t <* 5*/*30 *≈* 0.17. We use this value as an upper bound. Using values lower than this does not impact the result (Fig. S7K).

We find in Fig. S7J that over a range of *k_C_/k_N_* values around 0.1, *P_obs,C_*(*t*)*/P_obs,N_* (*t*) is no larger than *∼* 1 in our null model for a wide range of *t/*⌧*_f_* values above *∼* 0.15—we note that values lower than this are unreasonable anyways since the mass spectrometric intensity of C-terminal peptides and the intensity of the full-length western blot band at 15 mins are both inconsistent with only 15% having finished translation at this time. This is starkly inconsistent with the observation that, at 15 mins of translation, C-terminal peptides show a folded fraction *∼* 2 *—* 5 times larger than N-terminal peptides that fold significantly faster than the respective C-terminal regions during refolding from denaturant.

---

**fig. S1.**
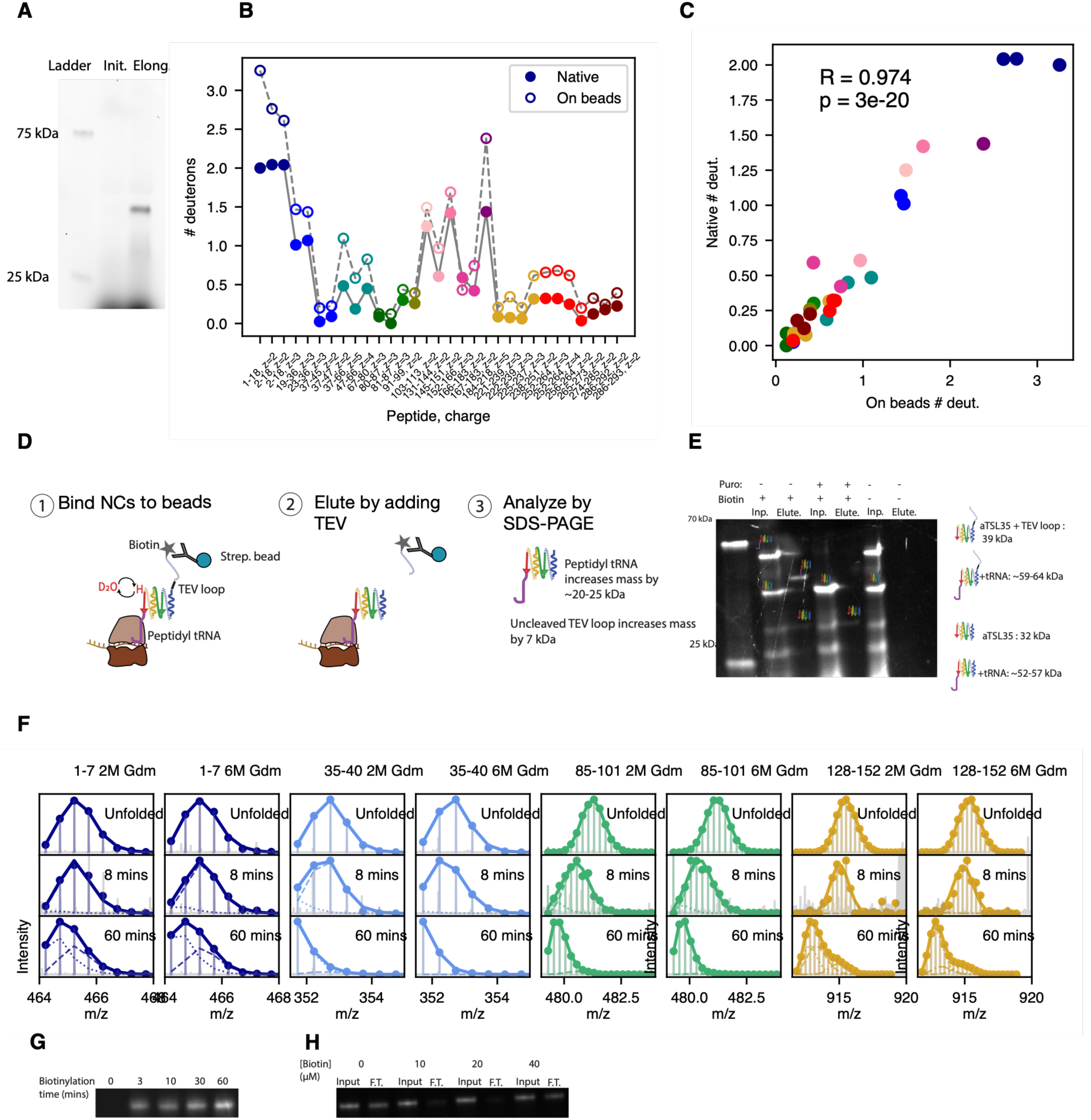
Validation of single-turnover translation and HDX-MS enrichment protocols. **(A)** Addition of 75 µM aurintricarboxylic acid (ATCA) at the elongation step leads to single-turnover translation. A synchronized translation reaction of aTS with Fluorotect GreenLys was treated with ATCA at either the initiation or elongation step, resolved via SDS-PAGE, and visualized by fluorescent imaging. Full-length aTS product is observed only when ATCA is added at the elongation step (right lane, “Elong.”) but not at the initiation step (left lane, “Init.”), indicating that ATCA blocks translation initiation without inhibiting actively elongating ribosomes. **(B)** Nascent protein enrichment coupled to HDX-MS does not globally disrupt protein structure. Pulse HDX-MS was performed on a HaloTag sample after 4 hours of elongation, enriched via biotin/streptavidin affinity capture. The number of deuterons accumulated per peptide in the protected mode (empty markers) is compared to a standard 10- second pulse-labeled HDX control on purified HaloTag protein (filled markers) in the same pH 6 buffer. **(C)** Correlation between peptide deuteration levels for the two experiments shown in (B). The high Pearson correlation coefficient and low p-value indicate a strong linear correlation. The slope of less than 1 indicates that deuterium uptake under enrichment conditions is systematically increased across the protein by a constant factor, suggesting that while the enrichment protocol (biotinylation and streptavidin bead binding) slightly destabilizes the protein and/or affects the extent of back exchange, it does not globally alter its folded structure. **(D)** Schematic of protocol for assessing nascent chain (NC) integrity following streptavidin bead binding. We translate a stalled NC construct with an N-terminal AviTag separated from the protein of interest by a linker containing a TEV loop. Following hydrogen-deuterium exchange and bead binding, TEV protease is added to elute NCs from beads, and samples are analyzed by SDS PAGE. If NCs are attached to the ribosome upon elution, then they will migrate ∼20-25 kDa larger than expected due to peptidyl tRNA attachment, a migration shift that vanishes if NCs are deliberately released from the ribosome via puromycin addition. For details, see Materials and Methods. **(E)** SDS-PAGE gel, imaged via fluorescence of NCs containing FluorTect GreenLys (see Methods), showing input (prior to bead binding) and elution (post TEV addition) samples as described in (D). Puromycin is added prior to bead binding, and biotin is included in IVT reactions, as indicated. We observe elutants migrate ∼20-25 kDa larger in the absence of puromycin than in its presence, indicating peptidyl tRNA attachment, so long as biotin is added to the translation reaction. The absence of biotin precludes streptavidin bead binding, and thus no elutants are observed. We note that elutants migrate 7 kDa faster than their respective input samples due to loss of N-terminal linker following TEV loop cleavage. **(E)** Mass spectra (represented as described in main text Fig. 2B) of various aTS-derived peptides derived from maximally deuterated control (“Unfolded”), 8 mins translation timepoint, and 60 mins translation timepoint following pulse-labeling HDX and bead binding reaction quenched with low pH buffer containing either 2M or 6M GdmCl. The high degree of similarity between trends observed in the spectra from these two conditions both during mid-translation (8 mins) and post-translation (6 mins) strongly suggests against the possibility that the observed mass spectrometric signal in our active translation experiment is biased towards spuriously released NCs due to acid-induced precipitation of peptidyl-tRNA attached species. If such precipitation were to occur, then increasing [GdmCl] from 2 to 6M would significantly reduce its extent and thus affect the observed signal. We note that the 6M GdmCl quench leads to slightly increased back exchange as observed in respective maximally deuterated control samples. (**F)** Determination of time required to biotinylate nascent chains: The protein HaloTag with an N-terminal AviTag and C-terminal 35 AA linker was translated *in vitro* in the presence of 20 µM biotin for 1.5 hours, then was supplemented with 75 µM ATCA (to prevent further initiation) and 1 µM BirA to begin biotinylation. At indicated times (including t=0, immediately prior to addition of BirA), a 4 µL reaction aliquot was drawn and translation was quenched with 18 µM puromycin at room temperature for 30 seconds, and samples were then mixed with Lamelli dye at 1X final concentration and immediately incubated at 95C for 5 mins to fully inactive BirA. All samples were then loaded on an SDS-PAGE gel and transferred to a PVDF membrane for anti-biotin western blotting with an Alexa-488 conjugated streptavidin probe. We conclude that nascent chain biotinylation is multiphasic, with a substantial degree of biotinylation occurring within a few minutes, indicating it is possible to appreciably enrich for nascent chains at early translation times. However, complete biotinylation requires hour timescales (H) Determination of optimal biotin concentration for nascent chain binding to streptavidin beads: The protein HaloTag with an N-terminal AviTag and C-terminal 35 AA linker was translated *in vitro* for 1 hour at 37C in the presence FluoroTect and variable biotin concentrations as indicated. Subsequently, 2 µL of each sample was drawn as an “input sample”, diluted in HKMT for a total volume of 8 µL, and frozen, then the remaining volume was bound for 2 hours with rotary mixing to a 15 µL aliquot of Pierce Streptavidin beads pre-equilibrated in HKMT. The flow-through was removed, and 8 µL of flow-through (F.T.) was drawn for gel analysis—we note that, if we assume no bead binding, these 8 µL aliquots would contain the same quantity of nascent chains as the input sample. All 8 µL samples were then treated with RNAse A to 1 mg/ml, followed by addition of Laemmli buffer to 1X concentration. Samples were then analyzed on a NUPAGE 4-12% Bis Tris gel and FluoroTect fluorescence was imaged. We conclude the 20 µM biotin corresponds to the optimal concentration for bead binding—increasing to 40 µM leads to reduced bead binding due to free biotin competing with streptavidin binding sites.

**fig. S2.**
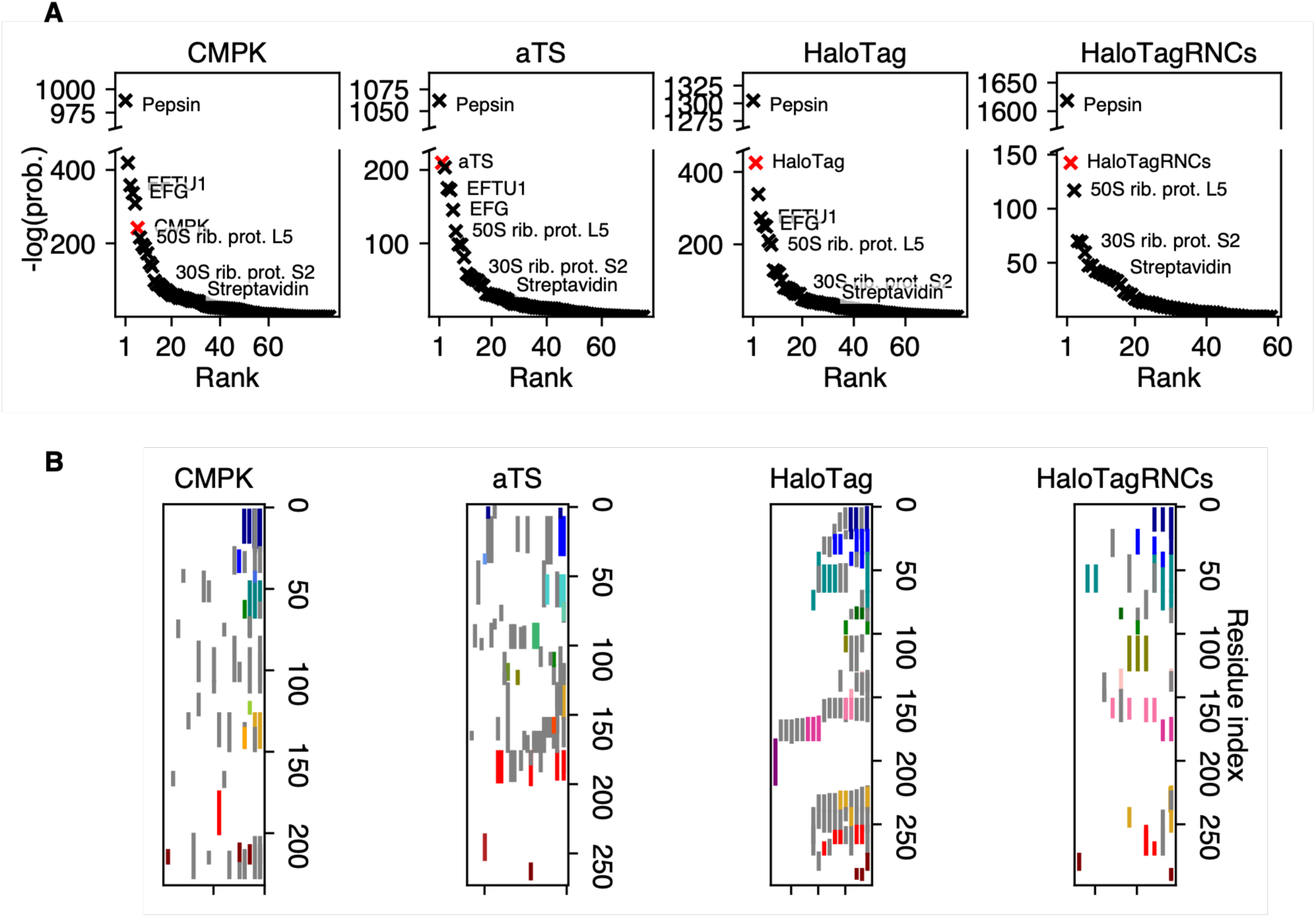
Nascent chain enrichment and coverage. **(A)** For each protein of interest that we analyze in this work (indicated above respective plots), we show confidence associated with identification of various protein species derived from our *in vitro* translation mixture, as well as proteins from our enrichment + digestion pipeline. Protein identification confidences are ranked by negative log p-values, with higher values indicating a lower probability of false assignment by the Byonic search algorithm. In each plot, the red marker indicates the protein of interest, while other proteins are shown in black. The positions corresponding to various proteins are annotated including pepsin (used for digestion), elongation factors EFTU1 and EFG, two ribosomal proteins L5 and S2, as well as streptavidin. For all proteins except CMPK, the protein of interest produces the highest confidence identification other than pepsin, which is added to bead slurry at high levels for digestion. The score corresponding to CMPK is slightly lower due to a lower digestion efficiency for this protein, but nonetheless high compared to the majority of proteins that were originally present in the non-enriched sample. HaloTagRNCs refers to a HaloTag + 34 AA C-terminal linker containing a strong SecM stalling sequence purified by sucrose cushion ultracentrifugation, which de-enriches non-ribosomal proteins from this sample. **(B)** Coverage maps indicating high confidence peptides detected from the indicated proteins. Each vertical bar represents a unique peptide + charge state combination that spans the residues indicated on the right axis provided its corresponding Pep2D value (a confidence metric used by Byonic) is less than 0.1, and colored bars represent high-quality peptides used for HDX-MS analysis. We note that these figures show representative coverage maps from a single replicate; the exact coverage varies slightly from replicate to replicate.

**fig. S3.**
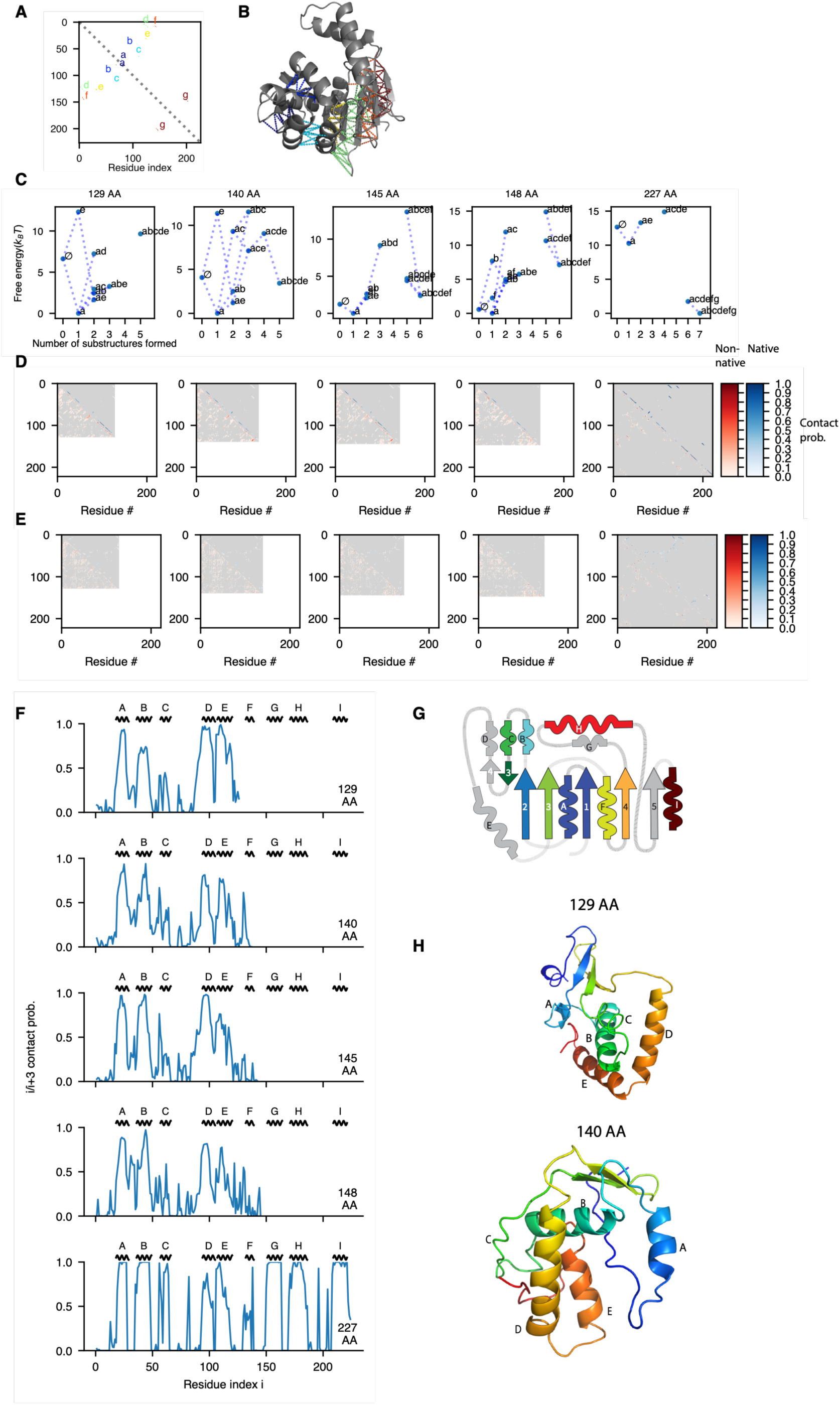
DBFOLD simulations predict that the CMPK NMP subdomain folds co-translationally while core folding is delayed until after translation Replica-exchange Monte-Carlo simulations with native-contact centric umbrella biasing using the MCPU method from reference (*29*) were reanalyzed as follows: **(A)** CMPK native contact map substructures, defined as in (*58*), shown in color and labeled. Each substructure defines a set of contiguous native contacts that is expected to form cooperatively during protein folding. For details on this substructure approach, see (*58*) **(B)** CMPK substructures mapped onto the native structure—each dashed line is a native contact belonging to a given substructure, color-coded as in (A). **(C)** Potentials of mean force (PMFs) showing free energy as a function of topological configuration—defined as a state in which a given subset of substructures is formed or absent-- for CMPK constructs in which various numbers of amino acids (AAs) have been synthesized. This analysis was performed using a slightly more lenient *f* parameter (defined as the maximum allowable ratio of the average distance between contacting residues in a given snapshot, to that same distance in the native state, such that a substructure involving those contacts is deemed present in the snapshot) than in reference (*29*) to more accurately capture the presence of dynamic inter-helix contacts. These PMFs are shown at a simulation temperature of T=0.5 (in arbitrary simulation energy units), at which point the folding free energy at full length (227 AA) is about 10 kBT, comparable to the experimentally-measured value of 6.4 kcal/mol for CMPK in (*69*) **(D)** Average alpha-carbon contact maps, computed using the MBAR method (*70*), as a function of translation length at a simulation temperature of T=0.5. Two alpha-carbons are deemed in contact in a given snapshot if their distance is within 6 Angstroms. The color shading reflects contact probability (as shown in colorbar) and blues in the upper triangular region reflect native contacts, while nonnative contacts are shown in shades of red in the lower triangular region. For details on this approach, see (*40*). This analysis reveals that native N-terminal alpha-helices form co-translationally with high probability, but native beta sheets (with the exception of beta hairpin belonging to the NMP-binding subdomain, indicated as substructure *a* in previous panels) do not form until translation is complete. **(E)** Same as (D) but now showing average side-chain contact maps, with a distance cutoff threshold of 5 Angstroms for defining a contact. These contact maps reveal a diversity of interconverting, nonnative tertiary packings at intermediate lengths, while stable, persistent native contacts do not form until full length as reflected by the dark shades of blue at length 227. **(F)** The third diagonal from the upper right triangular region in panel (D) is extracted at each chain length and plotted to reveal the probability of alpha helical contacts between each residue pair i and i+3. The locations for native alpha helices (labeled as in main text Fig. 2) are indicated above the panels. According to this analysis, DBFOLD simulations predict a high probability of NMP subdomain helix formation at intermediate chain lengths while core helices form post-translationally. The exception is core helix A, which is predicted to form co-translationally—this is in disagreement with experimental results and likely reflects an artificial over-stabilization of this helix in the simulation forcefield. **(G)** Topology map for CMPK with secondary structures labeled and color coded as in Fig. 2. **(H)** Sample simulation snapshots from indicated lengths with helices labeled. Note that these snapshots are shown in chainbow coloring that does not match the coloring shown in panel G.

**fig. S4.**
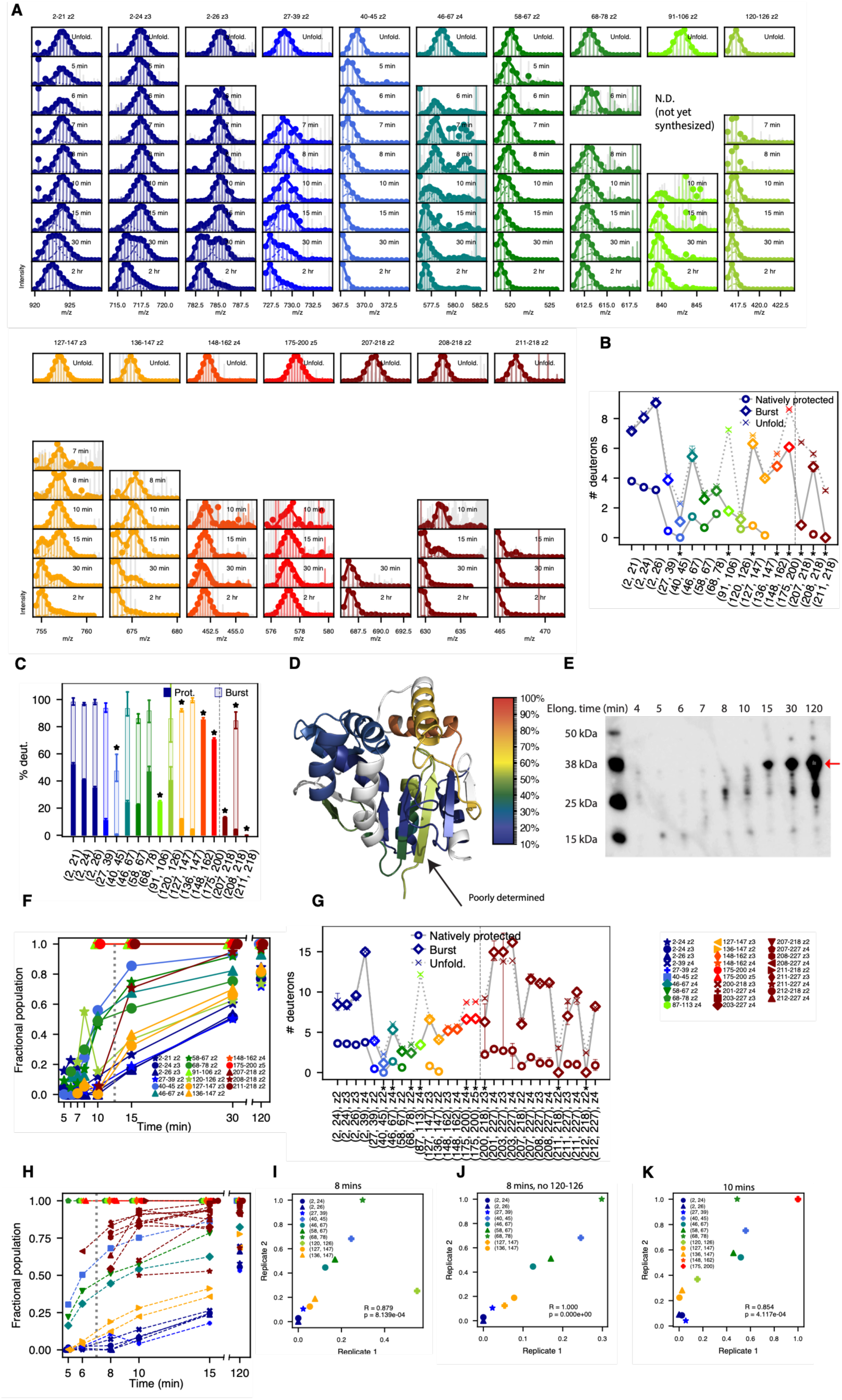
CMPK synchronized translation + pulse-labeling HDX-MS. **(A)** Mass spectra for all CMPK peptides derived from unfolded control (top row) and all elongation times, as in main text Fig. 2B. Empty panels reflect lack of peptide signal, either because the peptide is not yet synthesized or due to stochastic variabilities in mass-spectrometric detection for low-abundance peptides. **(B)** Number of deuterons associated with natively protected mode (circles), burst-phase mode (the unprotected mode observed early in translation, diamonds) and unfolded state (Xs) for each peptide. Unfolded deuteration levels are corrected for 10 second labeling (see Materials and Methods). Error bars represent 95% confidence intervals from bootstrapping and asterisks denote peptides exhibiting statistically significant burst-phase protection (see Methods). Peptides to the right of the dashed line are only detected after 10 mins (post-translation). **(C)** Same data as (B) but with deuterations expressed as a percentage of maximally deuterated value. **(D)** Percentage deuterations of all peptides are mapped onto the CMPK structure (PDB ID *2cmk*). In general, protection from deuteration correlates with secondary structure. Beta strand 1 appears to show anomalously high deuteration as it is only spanned by large peptides that encompass significant disordered regions, and hence its deuteration cannot be accurately determined. **(E)** Anti-biotin western blot (as in main text Fig. 1C.) showing progress of CMPK synchronized translation reaction as a function of time. This blot reflects the replicate that was used for pulse-labeling experiments shown in main text Fig. 2 and panel (A). Red arrow indicates full-length protein. At around 8 mins, we observe appreciable accumulation of a ∼26 kDa translation intermediate, which corresponds to translation of roughly 180 AA from CMPK (with additional ∼6 kDa due to the N terminal AviTag + linker). At this stage, the lid subdomain (residues 36-120) has fully emerged from the ribosomal exit tunnel, which typically sequesters the last 30-50 AA. Our observation that lid folding begins around this time (Fig. 2C) strongly suggests that this subdomain folds shortly after its emergence from the ribosome. **(F)** Same as main text Fig. 2C, now showing all peptides with high-quality mass spectra and replicate charge states where present **(G)** Same as (B) for replicate CMPK synchronized translation reaction. (**H)** Same as main text Fig. 2C for a replicate CMPK synchronized translation reaction. As in the first replicate, we observe that lid peptides synchronously begin acquiring protection prior to completion of synthesis, followed by post-translational core folding and rapid NMP subdomain folding. **(I)** To confirm peptides acquire protection in a reproducible sequence across replicates, we compare, for each peptide, the fractional population associated with the natively protected mode at 8 mins in both replicates. Peptides are color coded as in previous panels. A high Spearman correlation coefficient and low p value (shown on the plot) indicate a high degree of reproducibility. **(J)** Same as (I) omitting peptide 120-126 which is poorly fit in the first replicate (see panel A). **(K)** Same as (I) now shown at the 10 mins timepoint.

**fig. S5.**
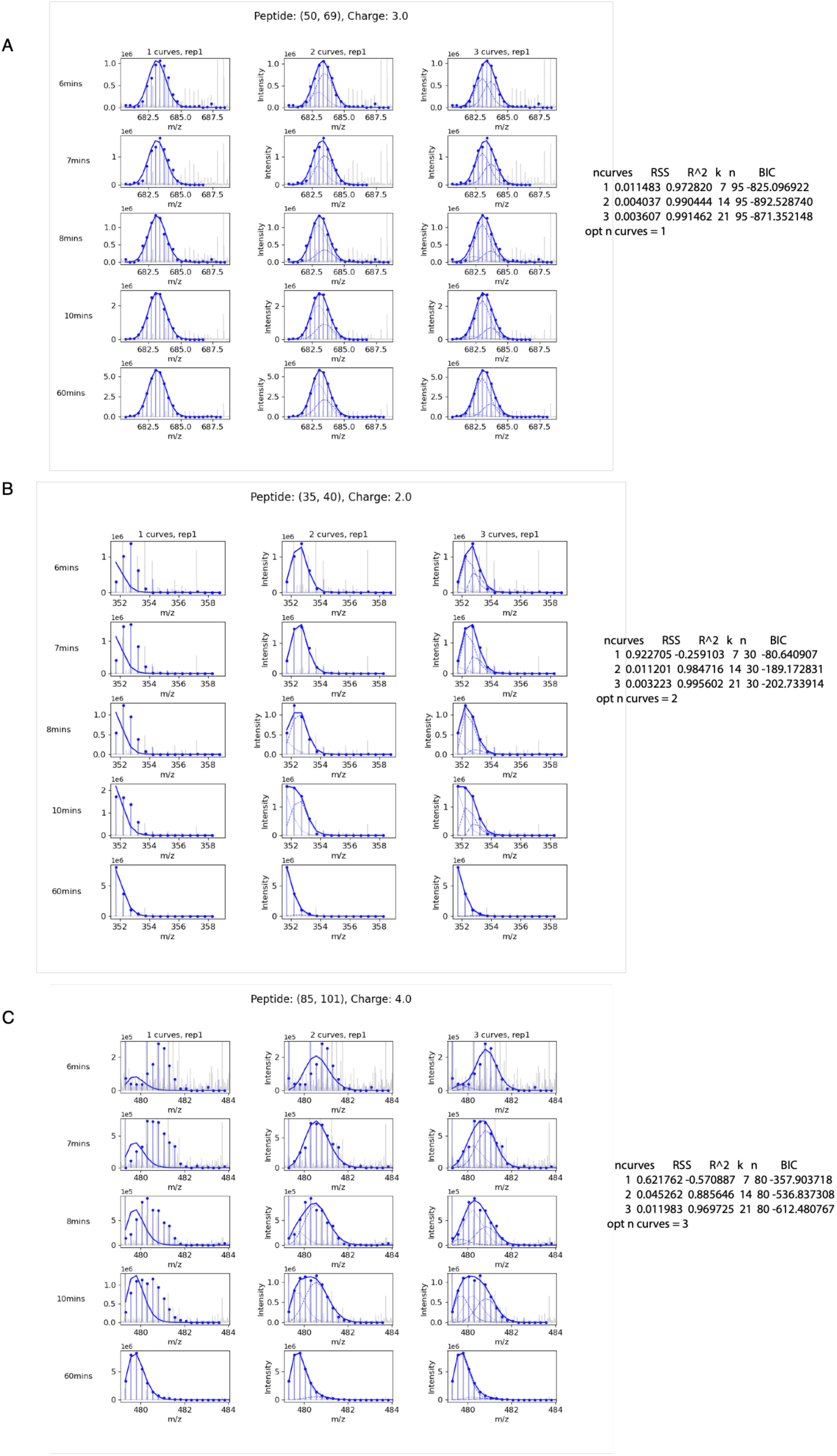
Determination of optimal number of curves for fitting aTS peptides. **(A)-(C)** For indicated peptides and charge states, we globally fit mass spectra from aTS synchronized translation experiments to either one, two or three curves. Mass spectra for each translation time and number of curves, alongside fits, are shown as in main text Fig. 2B. To the right of each set of plots, we indicate the RSS (residual sum of squares), R^2^ (goodness of fit), k (number of free fitting parameters), n (number of fitted datapoints) and BIC (Bayesian information criterion) associated with each number of curves, as well as the optimal number, determined as described in Materials and Methods.

**fig. S6.**
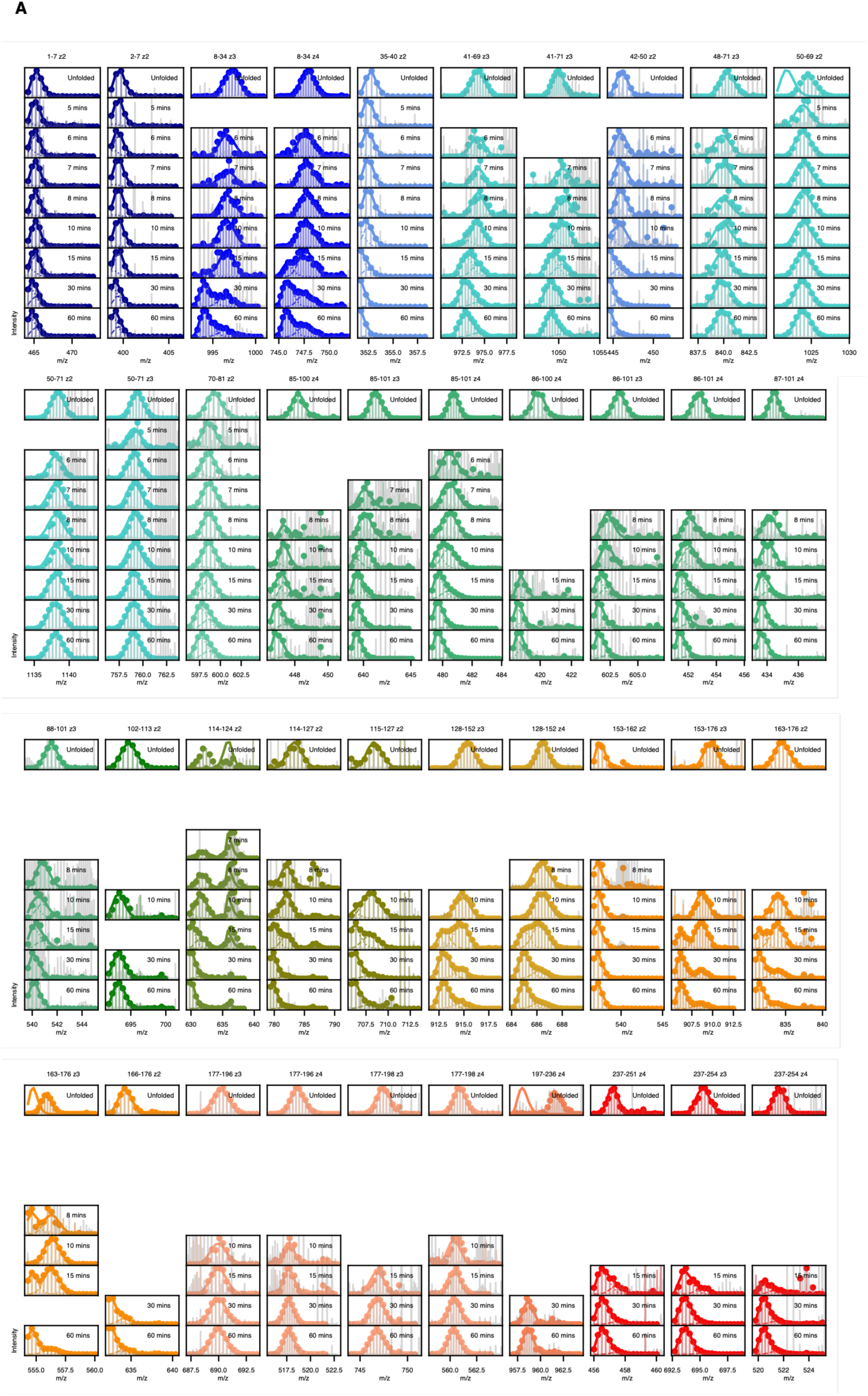
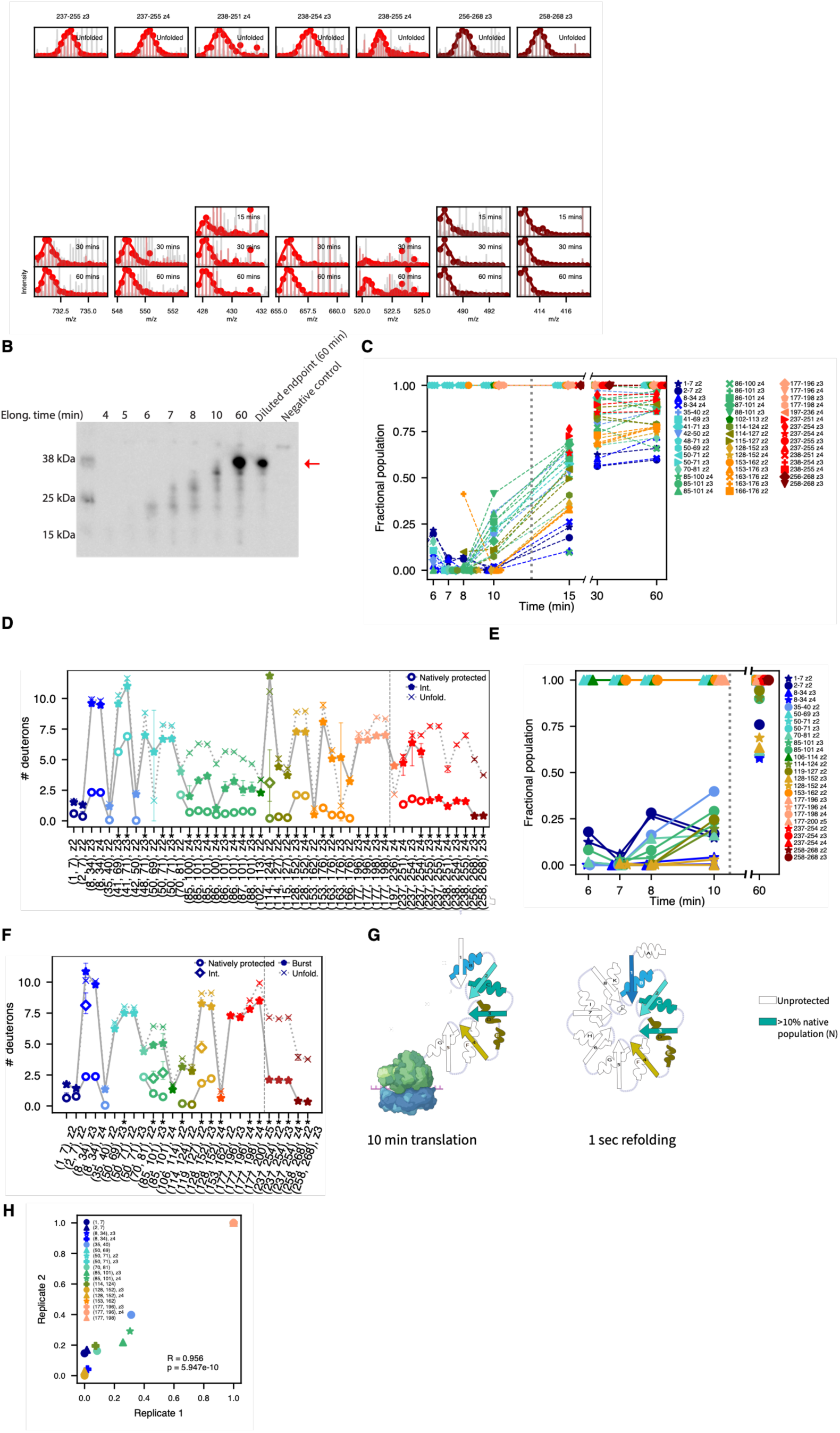
aTS synchronized translation + pulse-labeling HDX-MS. **(A)** Mass spectra for all aTS peptides derived from unfolded control (top row) and all elongation times, represented as in main text Fig. 2B. Empty panels reflect lack of peptide signal, either because the peptide is not yet synthesized or due to stochastic variabilities in mass-spectrometric detection for low-abundance peptides. **(B)** Anti-biotin western blot (as in main text Fig. 1C.) showing progress of aTS synchronized translation reaction as a function of time. This blot reflects the replicate that was used for pulse-labeling experiments shown in main text Fig. 2 and panel (A). Red arrow indicates full-length protein. (**C)** Same as main text Fig. 2G, now showing all peptides with high-quality mass spectra and replicate charge states where present. **(D)** Number of deuterons associated with natively protected mode (circles), intermediately protected mode where present (diamonds), burst-phase mode (the deprotected mode observed early in translation, asteriks) and unfolded state (Xs) for each peptide. Unfolded deuteration are corrected for 10 second labeling (see Materials and Methods). Error bars represent 95% confidence intervals from bootstrapping and asterisks denote peptides exhibiting statistically significant burst-phase protection (see Methods). Peptides to the right of the dashed line are only detected after 10 mins (post-translation). Various aTS peptides optimally fit to a single mode, For these peptides, only a star and x are shown, representing deuteration of the single global fit mode and the unfolded mode, respectively. **(E)** Same as panel C for a replicate aTS synchronized translation reaction. **(F)** Same as (D) for replicate aTS synchronized translation reaction **(G)** Comparison of folding intermediates populated during aTS translation and refolding from denaturant. Left panel shows secondary structures spanned by peptides showing a natively protected fractional population of at least 10%, provided the deuterium uptake in this native mode is no more than 50% that in the maximally deuterated control, as in main text Fig. 2H, while right panel shows structures spanned by at least one peptide showing a natively protected fractional population of at least 10% at 1 second of refolding from denaturant in the pulse-labeling HDX-MS experiment from Wintrode et. al. 2005 (*50*). For numerical values, see table S4. The co-translational and refolding intermediates largely resemble each other with the exceptions of beta strand 1, which is protected during refolding but not in the co-translational intermediate. **(H)** To confirm peptides acquire protection in a reproducible sequence across replicates, we compare, for each peptide, the fractional population associated with the natively protected mode at 10 mins in both replicates. Peptides are color coded as in previous panels. A high Spearman correlation coefficient and low p value (shown on the plot) indicate a high degree of reproducibility.

**Table S4:**
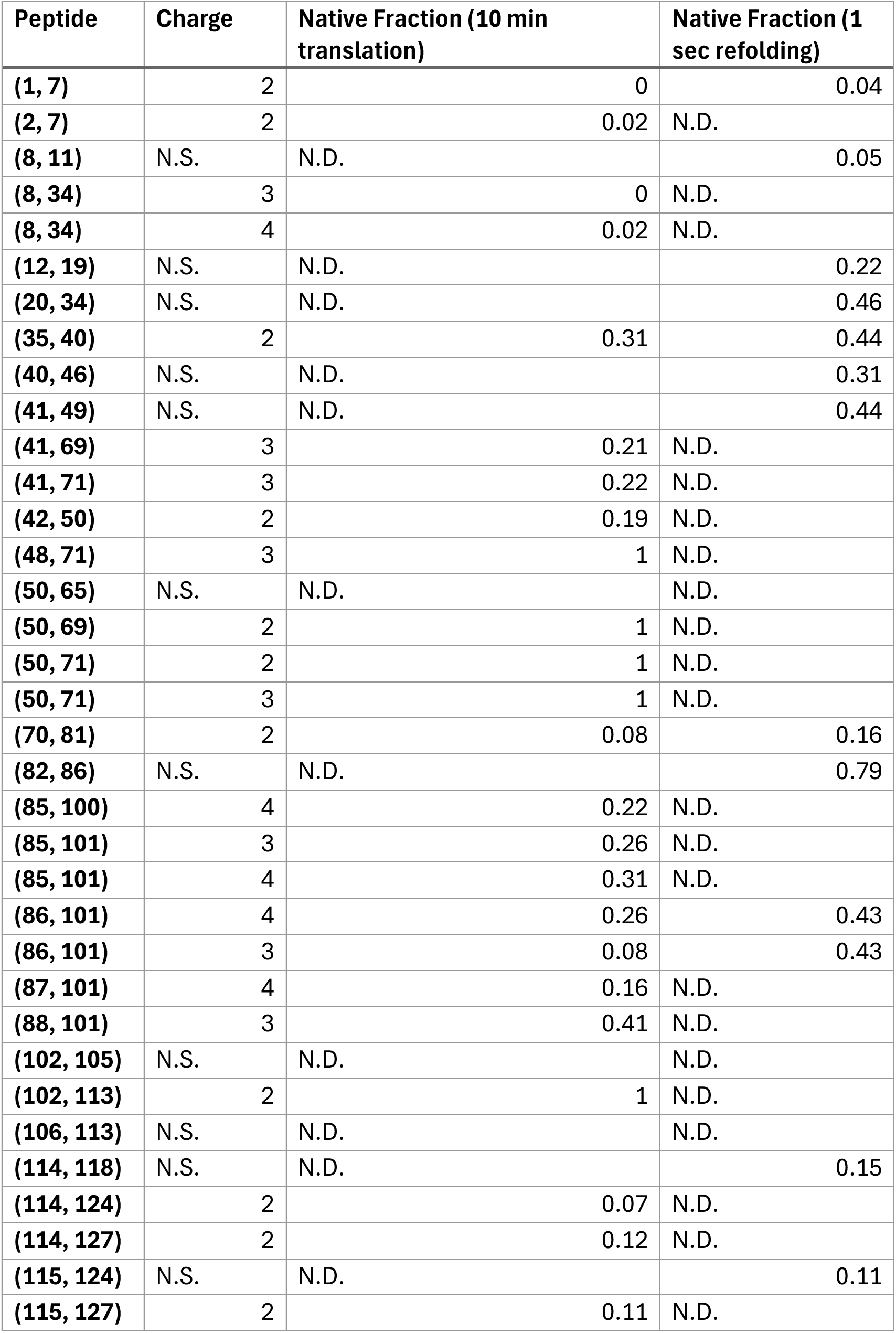
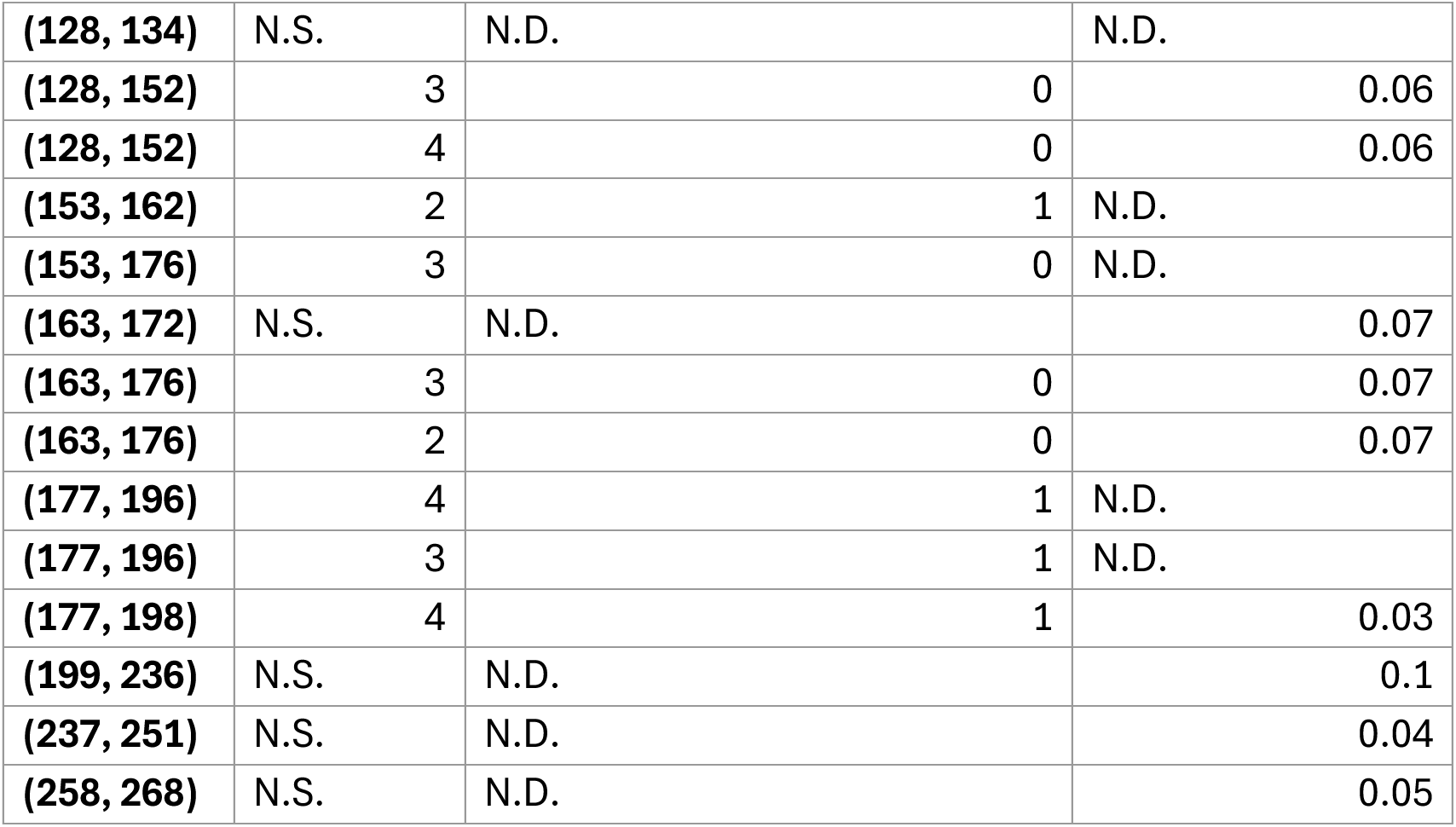
aTS co-translational folding intermediate resembles refolding intermediate from denaturant. For each peptide and charge state, we indicate the fraction protected in both the 10 minute co-translational timepoint (our experiment) and the 1 second refolding timepoint (Wintrode et. al. 2005) (*50*). N.D. indicates peptide is not detected in the respective experiment, and N.S. indicates that the charge state was not specified for peptides in Wintrode et. al. 2005.

**fig. S7:**
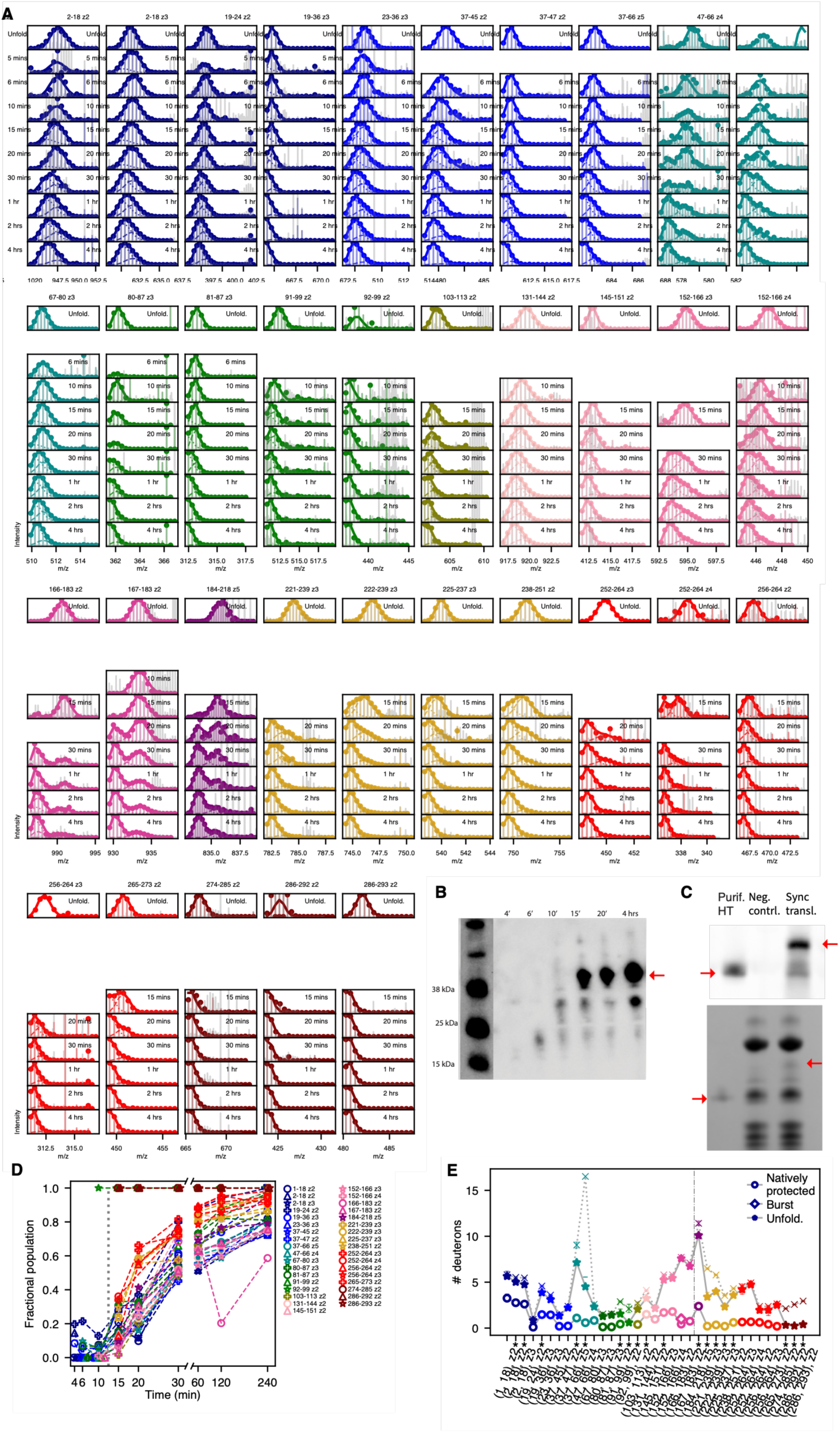
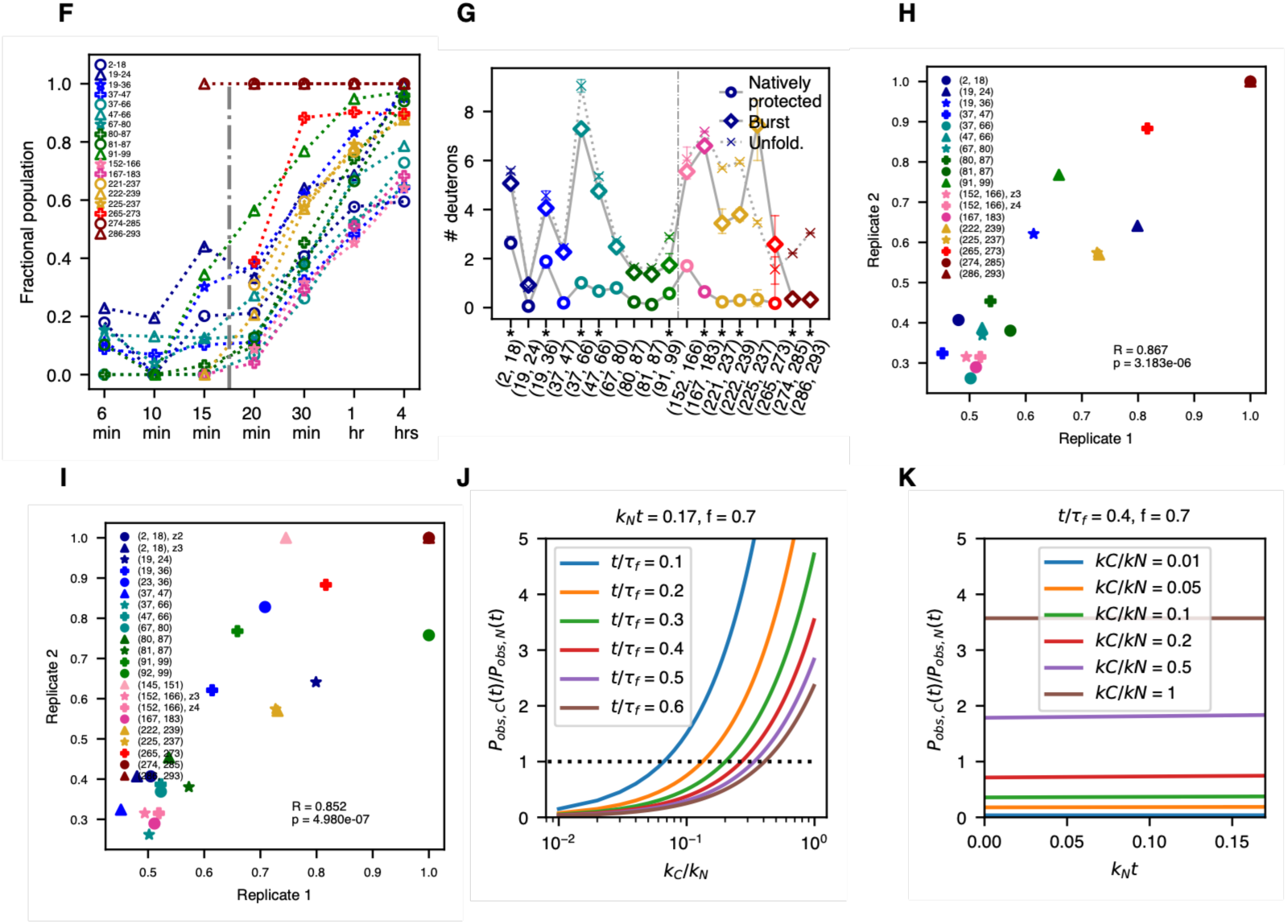
HaloTag synchronized translation + pulse-labeling HDX-MS. **(A)** Mass spectra for all HaloTag peptides derived from unfolded control (top row) and all elongation times, represented as in main text Fig. 2B. Empty panels reflect lack of peptide signal, either because the peptide is not yet synthesized or due to stochastic variabilities in mass-spectrometric detection for low-abundance peptides. **(B)** Anti-biotin western blot (as in main text Fig. 1C.) showing progress of HaloTag synchronized translation reaction as a function of time. This blot reflects the replicate that was used for pulse-labeling experiments shown in main text Fig. 2 and panel (A). Red arrow indicates full-length protein. **(C)** Top: To verify HaloTag folds into its native structure in our experimental conditions, we perform a synchronized translation reaction in the presence of 10 µM of fluorescent HaloTag ligand TMR and load the final timepoint on an SDS-PAGE gel which we fluorescently image to confirm covalent TMR bidning (middle lane). As a negative control, we perform an equivalent translation reaction without the HaloTag gene (rightmost lane), and we also load purified HaloTag that was similarly incubated with 10 µM TMR as a positive control (leftmost lane). Translated HaloTag runs larger than purified protein due to the added presence of the N-terminal AviTag and linker. Truncated fragments are also observed to bind TMR to some extend as seen previously (*9*). Bottom: Coomassie-stained gel from above. In both images, red arrows indicate HaloTag bands. (**D)** Same as main text Fig. 3C now showing all peptides with high-quality mass spectra and replicate charge states where present. **(E)** Same as main text Fig. 3D now showing all peptides with high-quality mass spectra and replicate charge states where present. **(F)** Same as main text Fig. 3C for a replicate HaloTag synchronized translation reaction. As in our first replicate, peptides from across the protein do not begin showing native protection until after translation is complete, and we observe protection in three similar stages, namely 1) rapid folding of C-terminal most helices, 2) folding of C-terminal regions of the core, with limited involvement from the N-terminus, and 3) folding of the majority of N-terminus and lid. We note that translation occurred at a slightly slower and yielded slightly reduced peptide coverage than the initial replicate, particularly near the C-terminus. **(G)** Same as main text Fig. 3C for replicate HaloTag synchronized translation reaction. As in the initial replicate, we observe subtle albeit significant burst-phase protection at various N-terminal peptides. We note that for peptide 91-99, the maximally deuterated control for this replicate gave poor spectra, and we thus used the spectra from the initial replicate instead. We validated this substitution by verifying that the maximally deuterated for this peptide when derived from stalled nascent chains showed nearly identical deuteration. (**H)** To confirm that the order of peptide protection is consistent across replicates, we compare, for each peptide, the fractional population associated with the natively protected mode at 30 mins in both replicates. The x and y axis denote the native population fraction in the two respective replicates at the 30 min timepoint. Peptides are color coded as in previous panels. A high Spearman correlation coefficient and low p value (shown on the plot) indicate a high degree of reproducibility. This plot only includes the subset of peptides with high-quality fits in both replicates that are plotted in main text Fig. 3. **(I)** Same as (H) now showing *all* peptides common to both replicates. (**J)** Plots of eq. 12 in supplementary note 3 as a function of *kC/kN* for indicated parameter values. (**K)** Same as J, now as a function of *kNt*.

**Fig. S8:**
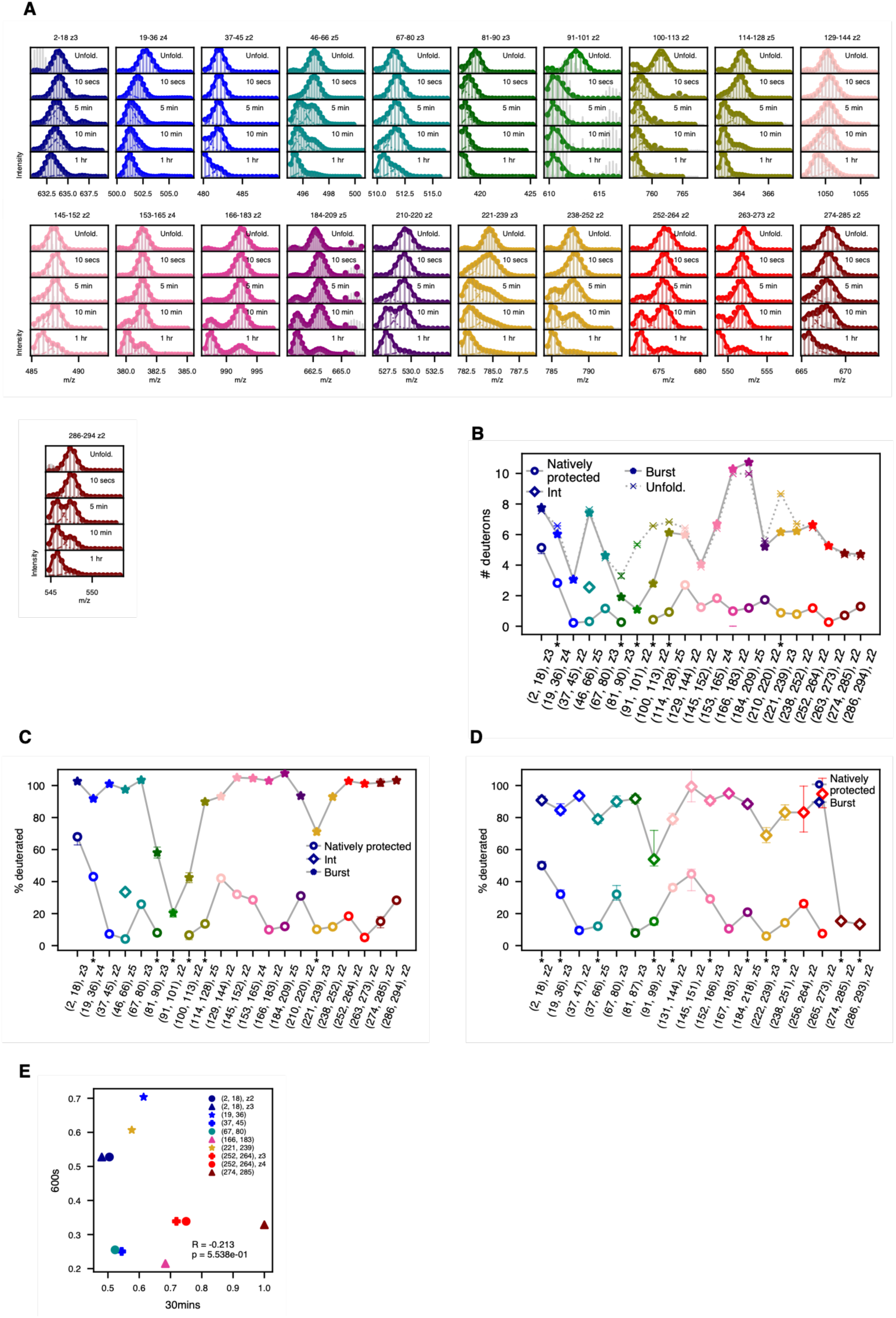
HaloTag refolding from denaturant + pulse-labeling HDX-MS. **(A)** Mass spectra for all HaloTag peptides derived from unfolded control (top row) and all timepoints during refolding from denaturant, as in main text Fig. 2B. Data were obtained from reference (*43*) and reanalyzed using the same global fitting approach applied to co-translational experiments as described in Materials and Methods . **(B)** Number of deuterons associated with natively-protected mode (circles), intermediately-protected mode where present (stars), burst-phase mode (the deprotected mode observed early in translation, diamonds) and unfolded state (Xs) for each peptide. Error bars represent 95% confidence intervals from bootstrapping and asterisks denote peptides exhibiting statistically significant burst-phase protection (see Methods). (**C)** Same as (B) with all # of deuteron values expressed as a percentage relative to the respective values in the unfolded state. (**D)** Same as main text Fig. 3D, which shows deuteration of native and burst-phase modes in active translation experiment, with all # of deuteron values expressed as a percentage relative to the respective values in the unfolded state. We note that the pattern of burst-phase protection differs from that observed in the refolding experiment, with generally higher protection seen across the N terminus. This strongly suggests that the burst-phase intermediate in co-translational folding differs from that observed in refolding. **(E)** Correlation scatterplot between natively protected fractions in 30 minutes timepoint from first replicate of co-translational folding experiment (x-axis) and 10-minute timepoint of refolding experiment (y-axis). At these timepoints, all peptides have begun to show some degree of native population in both experiments. The low Spearman correlation provides further support that these proceed through different intermediates.

**Fig. S9:**
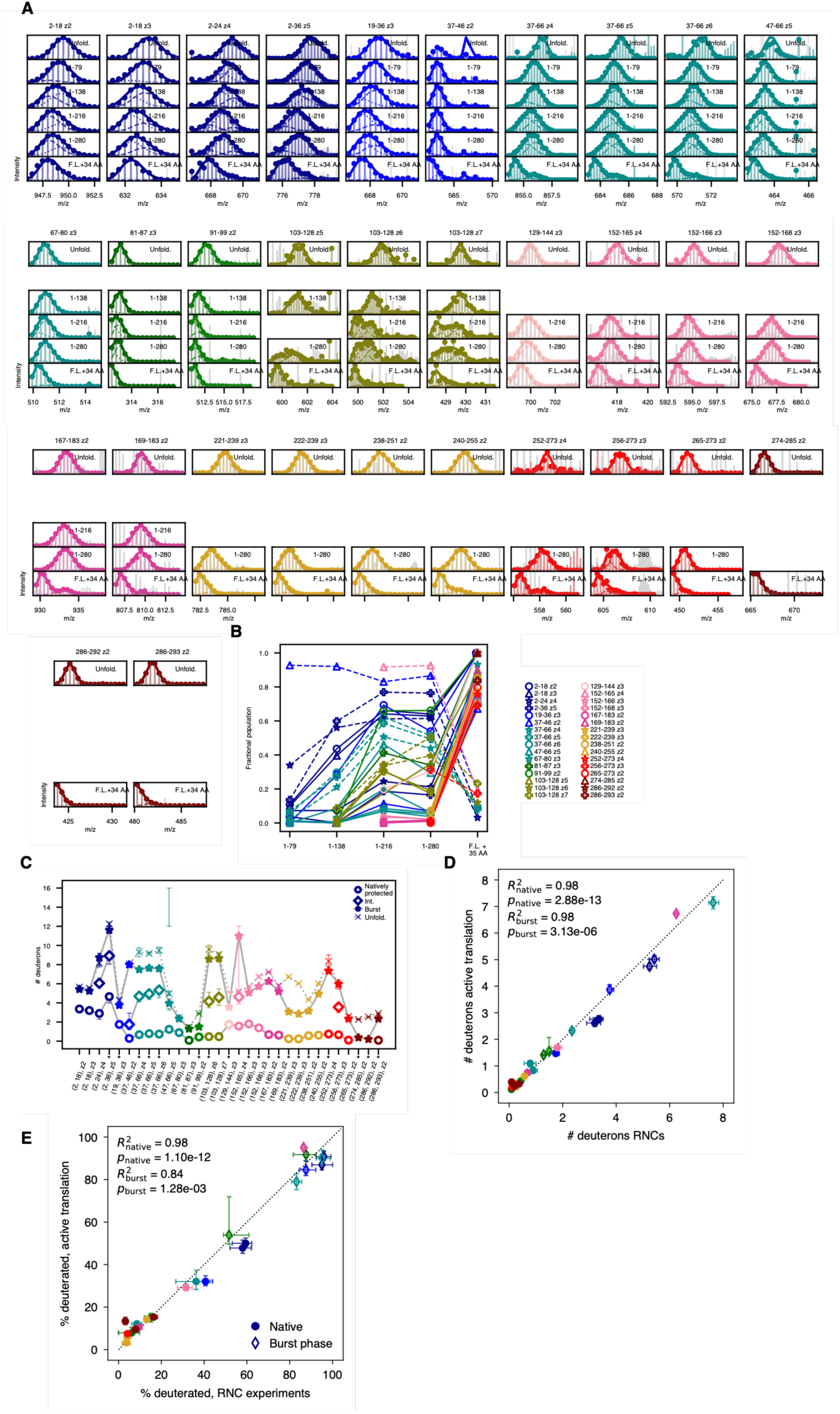
HaloTag stalled ribosomal nascent chains (RNCs) + pulse-labeling HDX-MS. (A) Mass spectra for all HaloTag RNC peptides derived from unfolded control (top row) and all translation lengths, as in main text Fig. 4B. Empty panels reflect lack of peptide signal, either because the peptide is not yet synthesized at that length or due to stochastic variabilities in mass-spectrometric detection for low-abundance peptides. **(B)** Same as main text Fig. 4E, now showing all peptides with high-quality mass spectra and replicate charge states where present. **(C)** Number of deuterons associated with natively protected mode (circles), intermediately protected mode where present (stars), burst-phase mode (the deprotected mode observed early in translation, diamonds) and unfolded state (Xs) for each peptide. Unfolded deuteration are corrected for 10 second labeling (see Materials and Methods). Error bars represent 95% confidence intervals from bootstrapping and asterisks denote peptides exhibiting statistically significant protection, relative to the maximally deuterated control corrected for 10 sec labeling, in the deprotected mode (see Materials and Methods). **(D)** Correlation between deuterium uptake associated with native (circles) and burst phase (diamonds) modes for each peptide in RNC and active translation experiments with indicated Pearson correlations. Error bars represent 95% confidence intervals from bootstrapping. **(E)** Same as (D) but with all values normalized by respective maximally deuterated control values. This normalization corrects for the confounding effect of peptide length.

## Notes

### Competing Interest Statement

The authors have declared no competing interest.

### Summary of Updates

Revised various aspects of text and figures for clarity prior to submission. Results themselves are unchanged

