## Supplementary Materials for "A detailed molecular picture of protein folding during active translation"

$$I_{m/z,t} = A \sum_{m=1}^{N_c} f_t^m \sum_{i=0}^{N} \mathrm{Binom}(i; N_{\mathrm{eff}}^m, p_m) * I_0\left(\frac{(m-i)}{z}\right)$$

Where $A$ describes the signal amplitude related to factors such as digestion and ionization efficiency, $N_c$ is the number of curves being fit, $f_t^m$ is the fraction of signal associated with mode $m$ at time $t$ (at each timepoint, these must sum to 1 over all $m$), $N$ is the total number of exchangeable amides in the peptide, $\mathrm{Binom}(i; N_{\mathrm{eff}}^m, p_m)$ refers to a normalized binomial distribution evaluated at integer $i$ with parameters $N_{\mathrm{eff}}^m$ and $p_m$ (the product of which is the average number of deuterons taken up for mode $m$), and $I_0(m/z)$ is the mass spectral intensity for the natural abundance isotopic distribution for the undeuterated peptide at m/z. In this formulation, we neglect the instrument-induced mass spectrometric peak width as well as any interference from other overlapping peptides.

$$R_{N_c}^2 = 1 - \frac{\sum_{i=1}^{n_s}\left(I_i^{N_c} - I_i\right)^2}{\sum_{i=1}^{n_s}(I_i - \bar{I})^2}$$

Where $\bar{I}$ is the average intensity over per-timepoint normalized spectra. From these values, we determine $N_R$, which is the smallest value of $N_c$ required to achieve $R_{N_c}^2 > 0.95$. Finally, the optimal number of curves is chosen as:

$$\{k_i\}_{a \dots b} = 0 \quad \text{for } i = a, a+1$$
$$k_i = k_i \quad \text{otherwise}$$

where the first two peptide residues have an effective exchange rate of zero because they back-exchange very quickly following digestion (the first residue, in fact, contains a free amine rather than an amide). We then compute the corrected deuteration for the unfolded state as:

$$D_{m,\text{corr}} = f D_{m,\text{max}}$$

where:

$$f = \frac{1}{b-a-2} \sum_{i=a+2}^{b} (1 - e^{-k_i t})$$

Where $t = 10$ s, assuming single-exponential exchange kinetics for each site, and $D_{m,\text{max}}$ is the number of deuterons exchanged in the maximally deuterated control. For most peptides, we obtain $f$ values in the range of $\sim 0.9$ to 1. We likewise apply this correction to the centroid obtained for all bootstrap iterations performed on the maximally deuterated control to obtain a corrected $P(D_{m,\text{corr}})$ for the expected deuteration of an unfolded polypeptide following 10 s of pulse labeling under our experimental conditions. Finally, we compute the p-value associated with the null hypothesis that $D_m \geq D_{m,\text{corr}}$ as the fraction of bootstrap trials for which the observed centroid is at least as large as the corrected maximally deuterated centroid as an approximation to:

$$p = \int_0^\infty d\, D_{m,\text{corr}} P(D_{m,\text{corr}}) \int_{D_{m,\text{corr}}}^\infty d\, D_m P(D_m)$$

For any peptides for which $p < 0.01$ for the least protected mode and for which $D_{m,\text{corr}} - D_m \geq 0.5$, we reject the null hypothesis and conclude that the least protected mode is significantly more protected than would be expected if it reflected the unfolded state. This same definition of statistically significant burst-phase protection is also used to designate which peptides are shown with stripped shading in Fig. 2H, with the additional requirement that this partially protected mode show no more than 85% deuterium uptake in this specific panel. For fig. S9E which shows mode deuterations normalized by the maximally deuterated value, the error bars are obtained via element-wise divided samples from the bootstrapped distribution for the respective mode and that for the corrected maximally deuterated control, respectively and then extracting the 95% confidence intervals from the resulting distributions.

We note that, in main text figures showing fractional populations as a function of time and mode protection at each peptide, a subset of high-quality peptides showing poor global fits is excluded. In addition, in cases where multiple charge states exist for a given peptide, the main text figures present only the charge state showing the stronger signal and/or higher quality fitting. In the supplementary figures, we present these graphs with all high-quality peptides and redundant charge states included. We observe reproducible behavior across charge states for a given peptide, even though these are fit and analyzed independently (see supplementary figures).

**End of Translation Determination**

As a lower-bound estimate for the first timepoint associated with the end of translation, we considered, for each timepoint $t$, the most intense amplitude $A_{C\text{max},t}$ associated with any peptide/charge state combination spanning the most C-terminal residue for which we have coverage, as well as the least intense amplitude $A_{N\text{min},t}$ associated with any peptide spanning the most N-terminal residue for which we have coverage, and computed their ratio, $R_{\text{CN},t}$. We then normalized by this intensity ratio for the exact same peptide/charge state pair at the final translation timepoint, $R_{\text{CN},t_\text{final}}$, and determined the first timepoint $t$ for which:

| | **2-21, z2** | **2-24, z3** | **2-26, z3** | **27-39, z2** | **40-45, z2** | **46-67, z4** |
|---|---|---|---|---|---|---|
| **5mins** | 3.85E+07 | 5.22E+07 | | | 1.91E+07 | |
| **6mins** | 2.31E+07 | 6.31E+07 | 1.01E+07 | | 2.59E+07 | 1.46E+07 |
| **7mins** | 7.88E+07 | 1.18E+08 | 1.64E+07 | 2.93E+07 | 7.33E+07 | 1.53E+07 |
| **8mins** | 6.89E+07 | 1.20E+08 | 1.44E+07 | 2.42E+07 | 7.70E+07 | 2.66E+07 |
| **10mins** | 7.79E+07 | 1.79E+08 | 3.14E+07 | 3.82E+07 | 1.19E+08 | 5.96E+07 |
| **15mins** | 8.88E+07 | 1.73E+08 | 3.11E+07 | 3.61E+07 | 1.05E+08 | 6.16E+07 |
| **30mins** | 1.97E+08 | 2.81E+08 | 5.81E+07 | 7.81E+07 | 2.64E+08 | 6.10E+07 |
| **2hrs** | 4.27E+08 | 6.56E+08 | 1.09E+08 | 1.49E+08 | 5.60E+08 | 1.90E+08 |

| | **58-67, z2** | **68-78, z2** | **120-126, z2** | **127-147, z3** | **136-147, z2** | **148-162, z4** |
|---|---|---|---|---|---|---|
| **5mins** | 1.52E+07 | | | | | |
| **6mins** | 1.93E+07 | 7.17E+06 | | | | |
| **7mins** | 1.10E+08 | | 1.03E+07 | 3.46E+06 | | |
| **8mins** | 1.18E+08 | 1.48E+07 | 2.45E+07 | 2.03E+07 | 2.04E+07 | |
| **10mins** | 1.65E+08 | 3.25E+07 | 7.40E+07 | 9.84E+07 | 1.09E+08 | 5.66E+06 |
| **15mins** | 1.69E+08 | 3.82E+07 | 9.68E+07 | 1.99E+08 | 1.92E+08 | 2.52E+07 |
| **30mins** | 4.58E+08 | 5.42E+07 | 2.71E+08 | 3.78E+08 | 6.80E+08 | 2.95E+07 |
| **2hrs** | 1.07E+09 | 1.52E+08 | 8.21E+08 | 7.74E+08 | 1.90E+09 | 1.19E+08 |

| | **175-200, z5** | **207-218, z2** | **208-218, z2** | **211-218, z2** |
|---|---|---|---|---|
| **5mins** | | | | |
| **6mins** | | | | |
| **7mins** | | | | |
| **8mins** | | | | |
| **10mins** | 2.04E+06 | | 1.72E+06 | |
| **15mins** | 3.40E+07 | | 3.08E+07 | 6.49E+06 |

| | | | | |
|---|---|---|---|---|
| **30mins** | 6.81E+07 | 1.93E+07 | 1.17E+08 | 4.04E+07 |
| **2hrs** | 2.24E+08 | 7.40E+07 | 4.19E+08 | 1.44E+08 |

**Table S1.**

Global fit amplitude, a quantification of overall mass spectrometric intensity, as a function of translation time for spectra corresponding to indicated CMPK peptide and charge state combinations. No value indicates the peptide was not detected at the indicated time

| | 20-36, z3 | 37-57, z3 | 139-153, z4 | 139-153, z3 | 171-179, z3 |
|---|---|---|---|---|---|
| **8mins** | 6.80E+08 | 4.93E+09 | 1.51E+09 | 7.77E+08 | 1.43E+09 |
| **10mins** | 5.40E+08 | 4.24E+09 | 1.57E+09 | 7.45E+08 | 1.48E+09 |
| **15mins** | 4.29E+08 | 3.31E+09 | 9.37E+08 | 4.61E+08 | 9.88E+08 |
| **30mins** | 4.29E+08 | 3.10E+09 | 7.49E+08 | 4.04E+08 | 9.08E+08 |

| | |
|---|---|
| **Average 10min to 15min ratio** | **1.46585922** |

**Table S2.**

Global fit amplitude, a quantification of overall mass spectrometric intensity, as a function of translation time for spectra corresponding to indicated peptide and charge state combinations from ribosomal protein L5 during translation reaction of CMPK. Only a subset of translation times is shown. Final row shows average ratio of intensity at 10 mins to that at 15 mins over all peptides.

| | 1-7, z2 | 1-7, z3 | 8-29, z4 | 8-29, z5 | 30-50, z5 | 33-50, z4 | 33-50, z5 |
|---|---|---|---|---|---|---|---|
| **8mins** | 6.30E+07 | 6.30E+07 | 8.96E+07 | 1.03E+08 | 7.84E+07 | 6.80E+07 | 4.17E+07 |
| **10mins** | 6.20E+07 | 7.17E+07 | 9.13E+07 | 1.02E+08 | 8.88E+07 | 7.60E+07 | 6.32E+07 |
| **15mins** | 5.74E+07 | 5.87E+07 | 7.19E+07 | 6.17E+07 | 7.73E+07 | 5.19E+07 | 3.16E+07 |
| **30mins** | | 6.85E+07 | 4.76E+07 | 6.08E+07 | 3.79E+07 | 5.01E+07 | 3.99E+07 |

| | |
|---|---|
| **Average 10min to 15min ratio** | **1.40545405** |

**Table S3.**
Global fit amplitude, a quantification of overall mass spectrometric intensity, as a function of translation time for spectra corresponding to indicated peptide and charge state combinations from ribosomal protein L23 during translation reaction of CMPK. Only a subset of translation times are shown. Final row shows average ratio of intensity at 10 mins to that at 15 mins over all peptides.

$$P_{obs,N}(t) = \frac{f \int_{-\infty}^{t} d\tau g_C(\tau) f_N(t-\tau)}{\int_{-\infty}^{t} d\tau g_N(\tau)} \tag{1}$$

$$P_{obs,C}(t) = \frac{\int_{-\infty}^{t} d\tau g_C(\tau) f_C(t-\tau)}{\int_{-\infty}^{t} d\tau g_C(\tau)} \tag{2}$$

Using $f_N(t) \geq f_C(t)$, we then have:

$$\frac{P_{obs,C}(t)}{P_{obs,N}(t)} \leq \frac{1}{f} \frac{\int_{-\infty}^{t} d\tau g_N(\tau)}{\int_{-\infty}^{t} d\tau g_C(\tau)} \tag{3}$$

Let us further assume that all ribosomes complete synthesis of the N-terminal region before any ribosome finishes translation of the full protein, in which case the integral in the numerator of Equation 3 is 1 for all $t$ of interest. Although this assumption may not be true in general, making this assumption allows us to set the strictest possible upper bound on $P_{obs,C}(t)/P_{obs,N}(t)$. In this case, we have:

$$\frac{P_{obs,C}(t)}{P_{obs,N}(t)} \leq \frac{1}{f}\frac{1}{\int_{-\infty}^{t} d\tau g_C(\tau)} \tag{4}$$

Or inverting, we have

$$\frac{P_{obs,N}(t)}{P_{obs,C}(t)} \geq f \int_{-\infty}^{t} d\tau g_C(\tau) \tag{5}$$

In other words, the observed ratio of the N-terminal folded fraction to the C-terminal folded fraction must be at least as large as the total fraction of ribosomes that has finished translated at time t.

$$\frac{P_{obs,C}(t)}{P_{obs,N}(t)} > \frac{1}{f}\frac{\int_{-\infty}^{t} d\tau g_N(\tau)}{\int_{-\infty}^{t} d\tau g_C(\tau)} \tag{8}$$

Since these peptides are adjacent, it is reasonable to assume that few ribosomes stall between them, so it is likely that $f \approx 1$. Moreover, using the fact that the numerator in Equation 8 is greater than the denominator, we would thus expect:

$$\frac{P_{obs,C}(t)}{P_{obs,N}(t)} > 1 \tag{9}$$

In reality, the expected lower bound would be much higher than this since 274–285 folds significantly more slowly than 286–293 during refolding. But even neglecting this, at all timepoints in our translation experiment, this ratio is actually 1, in violation of our null model.

### Null model under additional assumptions

Under a few additional reasonable assumptions, we can set even more strict bounds on the ratio of folded fractions that is expected under the null model at times shortly after the first ribosomes have completed synthesis. At these early times, for example, the 15 mins translation timepoint in the main-text replicate, it is reasonable to assume single-exponential folding kinetics, as the folding times will be much shorter than the characteristic timescales associated with any slower exponential rates in a multi-exponential process. Thus,

$$f_{N/C}(t) = 1 - e^{-k_{N/C}t} \tag{10}$$

where $k_{N/C}$ denote rate constants for N and C terminal regions, respectively. For analytical tractability, we further assume a uniform distribution of synthesis-completion times of width $\tau_f$. That is:

$$g_C(t) = \begin{cases} 1/\tau_f & \text{if } 0 \leq t \leq \tau_f \\ 0 & \text{otherwise} \end{cases} \tag{11}$$

Under these conditions, we have:

$$\frac{P_{obs,C}(t)}{P_{obs,N}(t)} = \frac{\tau_f}{f\ t}\frac{\int_{0}^{t} d\tau(1 - e^{-k_C(t-\tau)})}{\int_{0}^{t} d\tau(1 - e^{-k_N(t-\tau)})} \tag{12}$$

for $0 \leq t \leq \tau_f$. The integrals can be evluated and the result expressed in terms of dimensionless quantities:

$$\frac{P_{obs,C}(t)}{P_{obs,N}(t)} = \frac{1}{f} \frac{\left[1 - \frac{1}{k_N t}\left(\frac{k_C}{k_N}\right)^{-1}\left(1 - e^{-\left(\frac{k_C}{k_N}\right)k_N t}\right)\right]}{\frac{t}{\tau_f} - \frac{(t/\tau_f)}{k_N t}\left(1 - e^{-k_N t}\right)} \tag{13}$$

Plots of Equation 13 are shown in Fig. S7J as a function of $k_C/k_N$. We expect a value of approximately 0.1 from Dall et al., PNAS 2024. We further note that at 15 mins, the intensity ratio of the most intense C-terminal peptide to the least intense N-terminal peptide, normalized to this same quantity at the final timepoint, is roughly 0.4, so we set $t/\tau_f = 0.4$, although we consider a range of values. Finally, we note that in our translation experiment, N-terminal peptides fold at timescales slower than 30 mins, and that at the 15 min timepoint, the fastest ribosome must have finished translation less than 5 minutes ago (since no C-terminal peptides were observed at the 10 min timepoint). We thus insist that $k_N t < 5/30 \approx 0.17$. We use this value as an upper bound. Using values lower than this does not impact the result (Fig. S7K).

We find in Fig. S7J that over a range of $k_C/k_N$ values around 0.1, $P_{obs,C}(t)/P_{obs,N}(t)$ is no larger than $\sim 1$ in our null model for a wide range of $t/\tau_f$ values above $\sim 0.15$—we note that values lower than this are unreasonable anyways since the mass spectrometric intensity of C-terminal peptides and the intensity of the full-length western blot band at 15 mins are both inconsistent with only 15% having finished translation at this time. This is starkly inconsistent with the observation that, at 15 mins of translation, C-terminal peptides show a folded fraction $\sim 2 - 5$ times larger than N-terminal peptides that fold significantly faster than the respective C-terminal regions during refolding from denaturant.

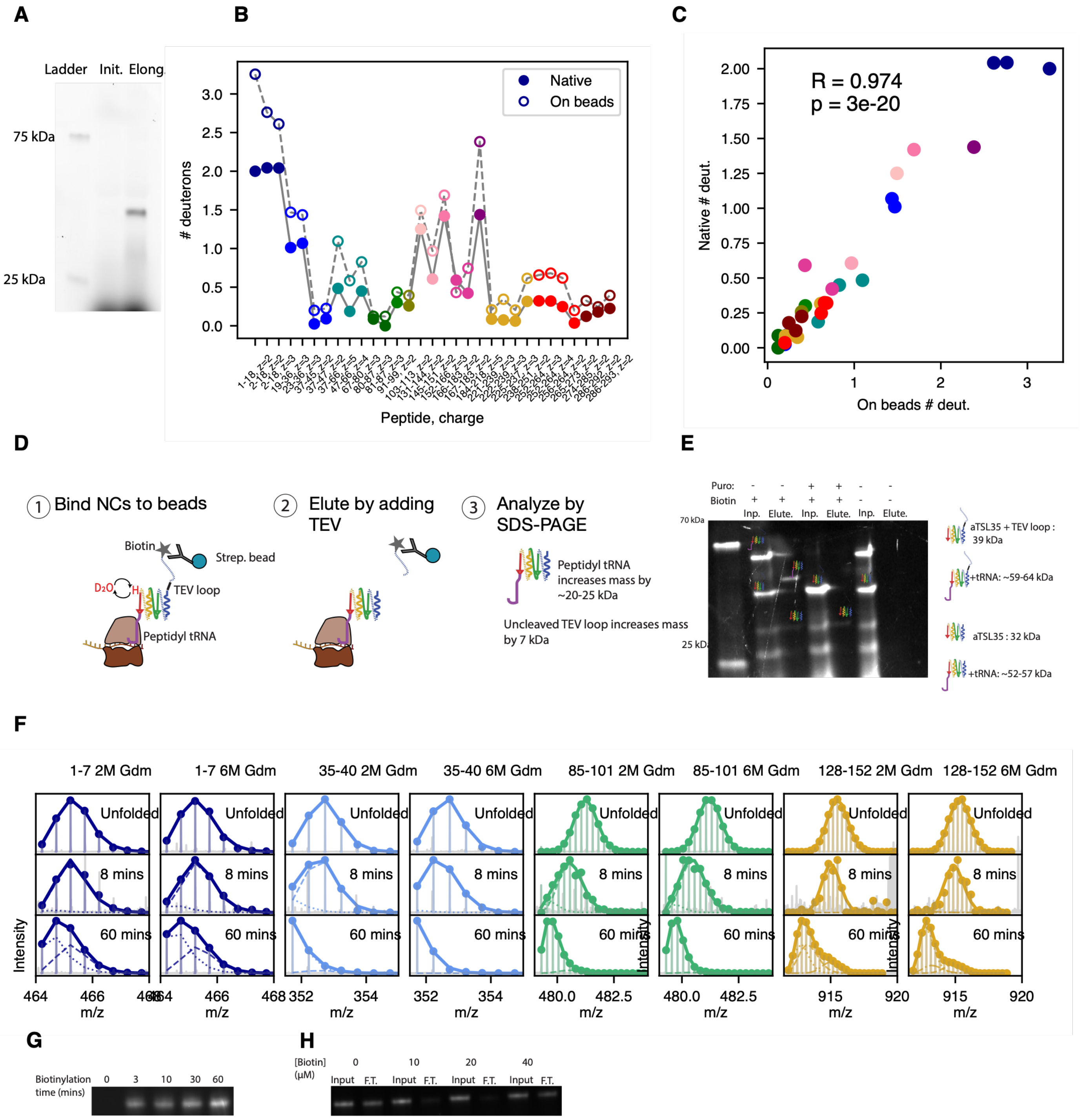
A
B
C
Ladder
Init.
Elong.
75 kDa
25 kDa
### deuterons
Native
On beads
Peptide, charge
R = 0.974
p = 3e-20
Native # deut.
On beads # deut.
D
E
1 Bind NCs to beads
2 Elute by adding TEV
3 Analyze by SDS-PAGE
Biotin
Strep. bead
TEV loop
Peptidyl tRNA
Peptidyl tRNA increases mass by ~20-25 kDa
Uncleaved TEV loop increases mass by 7 kDa
Puro:
Biotin
Inp.
Elute.
70 kDa
25 kDa
aTSL35 + TEV loop : 39 kDa
+tRNA: ~59-64 kDa
aTSL35 : 32 kDa
+tRNA: ~52-57 kDa
F
1-7 2M Gdm
1-7 6M Gdm
35-40 2M Gdm
35-40 6M Gdm
85-101 2M Gdm
85-101 6M Gdm
128-152 2M Gdm
128-152 6M Gdm
Unfolded
8 mins
60 mins
Intensity
m/z
G
H
Biotinylation time (mins)
[Biotin] (μM)
Input
F.T.

**fig. S1.** Validation of single-turnover translation and HDX-MS enrichment protocols

**(A)** Addition of 75 µM aurintricarboxylic acid (ATCA) at the elongation step leads to single-turnover translation. A synchronized translation reaction of aTS with Fluorotect GreenLys was treated with ATCA at either the initiation or elongation step, resolved via SDS-PAGE, and visualized by fluorescent imaging. Full-length aTS product is observed only when ATCA is added at the elongation step (right lane, "Elong.") but not at the initiation step (left lane, "Init."), indicating that ATCA blocks translation initiation without inhibiting actively elongating ribosomes. **(B)** Nascent protein enrichment coupled to HDX-MS does not globally disrupt protein structure. Pulse HDX-MS was performed on a HaloTag sample after 4 hours of elongation, enriched via biotin/streptavidin affinity capture. The number of deuterons accumulated per peptide in the protected mode (empty markers) is compared to a standard 10-second pulse-labeled HDX control on purified HaloTag protein (filled markers) in the same pH 6 buffer. **(C)** Correlation between peptide deuteration levels for the two experiments shown in (B). The high Pearson correlation coefficient and low p-value indicate a strong linear correlation. The slope of less than 1 indicates that deuterium uptake under enrichment conditions is systematically increased across the protein by a constant factor, suggesting that while the enrichment protocol (biotinylation and streptavidin bead binding) slightly destabilizes the protein and/or affects the extent of back exchange, it does not globally alter its folded structure. **(D)** Schematic of protocol for assessing nascent chain (NC) integrity following streptavidin bead binding. We translate a stalled NC construct with an N-terminal AviTag separated from the protein of interest by a linker containing a TEV loop. Following hydrogen-deuterium exchange and bead binding, TEV protease is added to elute NCs from beads, and samples are analyzed by SDS PAGE. If NCs are attached to the ribosome upon elution, then they will migrate ~20-25 kDa larger than expected due to peptidyl tRNA attachment, a migration shift that vanishes if NCs are deliberately released from the ribosome via puromycin addition. For details, see Materials and Methods. **(E)** SDS-PAGE gel, imaged via fluorescence of NCs containing FluorTect GreenLys (see Methods), showing input (prior to bead binding) and elution (post TEV addition) samples as described in (D). Puromycin is added prior to bead binding, and biotin is included in IVT reactions, as indicated. We observe elutants migrate ~20-25 kDa larger in the absence of puromycin than in its presence, indicating peptidyl tRNA attachment, so long as biotin is added to the translation reaction. The absence of biotin precludes streptavidin bead binding, and thus no elutants are observed. We note that elutants migrate 7 kDa faster than their respective input samples due to loss of N-terminal linker following TEV loop cleavage. **(E)** Mass spectra (represented as described in main text Fig. 2B) of various aTS-derived peptides derived from maximally deuterated control (“Unfolded”), 8 mins translation timepoint, and 60 mins translation timepoint following pulse-labeling HDX and bead binding reaction quenched with low pH buffer containing either 2M or 6M GdmCl. The high degree of similarity between trends observed in the spectra from these two conditions both during mid-translation (8 mins) and post-translation (6 mins) strongly suggests against the possibility that the observed mass spectrometric signal in our active translation experiment is biased towards spuriously released NCs due to acid-induced precipitation of peptidyl-tRNA attached species. If such precipitation were to occur, then increasing [GdmCl] from 2 to 6M would significantly reduce its extent and thus affect the observed signal. We note that the 6M GdmCl quench leads to slightly increased back exchange as observed in respective maximally deuterated control samples. (**F)** Determination of time required to biotinylate nascent chains: The protein HaloTag with an N-terminal AviTag and C-terminal 35 AA linker was translated *in vitro* in the presence of 20 µM biotin for 1.5 hours, then was supplemented with 75 µM ATCA (to prevent further initiation) and 1 µM BirA to begin biotinylation. At indicated times (including t=0, immediately prior to addition of BirA), a 4 µL reaction aliquot was drawn and translation was quenched with 18 µM puromycin at room temperature for 30 seconds, and samples were then mixed with Lamelli dye at 1X final concentration and immediately incubated at 95C for 5 mins to fully inactive BirA. All samples were then loaded on an SDS-PAGE gel and transferred to a PVDF membrane for anti-biotin western blotting with an Alexa-488 conjugated streptavidin probe. We conclude that nascent chain biotinylation is multiphasic, with a substantial degree of biotinylation occurring within a few minutes, indicating it is possible to appreciably enrich for nascent chains at early translation times. However, complete biotinylation requires hour timescales (H) Determination of optimal biotin concentration for nascent chain binding to streptavidin beads: The protein HaloTag with an N-terminal AviTag and C-terminal 35 AA linker was translated *in vitro* for 1 hour at 37C in the presence FluoroTect and variable biotin concentrations as indicated. Subsequently, 2 µL of each sample was drawn as an "input sample", diluted in HKMT for a total volume of 8 µL, and frozen, then the remaining volume was bound for 2 hours with rotary mixing to a 15 µL aliquot of Pierce Streptavidin beads pre-equilibrated in HKMT. The flow-through was removed, and 8 µL of flow-through (F.T.) was drawn for gel analysis—we note that, if we assume no bead binding, these 8 µL aliquots would contain the same quantity of nascent chains as the input sample. All 8 µL samples were then treated with RNAse A to 1 mg/ml, followed by addition of Laemmli buffer to 1X concentration. Samples were then analyzed on a NUPAGE 4-12% Bis Tris gel and FluoroTect fluorescence was imaged. We conclude the 20 µM biotin corresponds to the optimal concentration for bead binding—increasing to 40 µM leads to reduced bead binding due to free biotin competing with streptavidin binding sites.

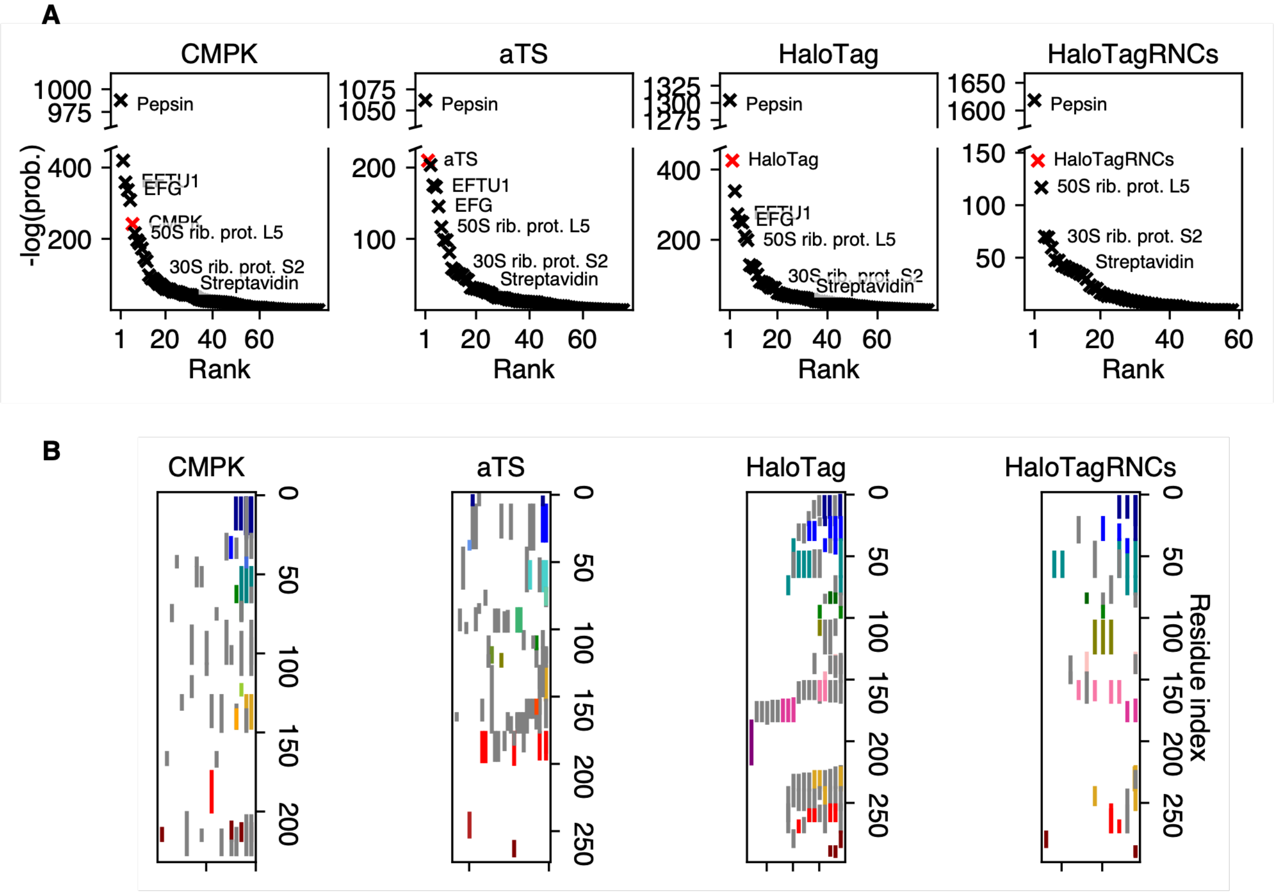


**fig. S2.** Nascent chain enrichment and coverage

**(A)** For each protein of interest that we analyze in this work (indicated above respective plots), we show confidence associated with identification of various protein species derived from our *in vitro* translation mixture, as well as proteins from our enrichment + digestion pipeline. Protein identification confidences are ranked by negative log p-values, with higher values indicating a lower probability of false assignment by the Byonic search algorithm. In each plot, the red marker indicates the protein of interest, while other proteins are shown in black. The positions corresponding to various proteins are annotated including pepsin (used for digestion), elongation factors EFTU1 and EFG, two ribosomal proteins L5 and S2, as well as streptavidin. For all proteins except CMPK, the protein of interest produces the highest confidence identification other than pepsin, which is added to bead slurry at high levels for digestion. The score corresponding to CMPK is slightly lower due to a lower digestion efficiency for this protein, but nonetheless high compared to the majority of proteins that were originally present in the non-enriched sample. HaloTagRNCs refers to a HaloTag + 34 AA C-terminal linker containing a strong SecM stalling sequence purified by sucrose cushion ultracentrifugation, which de-enriches non-ribosomal proteins from this sample. **(B)** Coverage maps indicating high confidence peptides detected from the indicated proteins. Each vertical bar represents a unique peptide + charge state combination that spans the residues indicated on the right axis provided its corresponding Pep2D value (a confidence metric used by Byonic) is less than 0.1, and colored bars represent high-quality peptides used for HDX-MS analysis. We note that these figures show representative coverage maps from a single replicate; the exact coverage varies slightly from replicate to replicate.

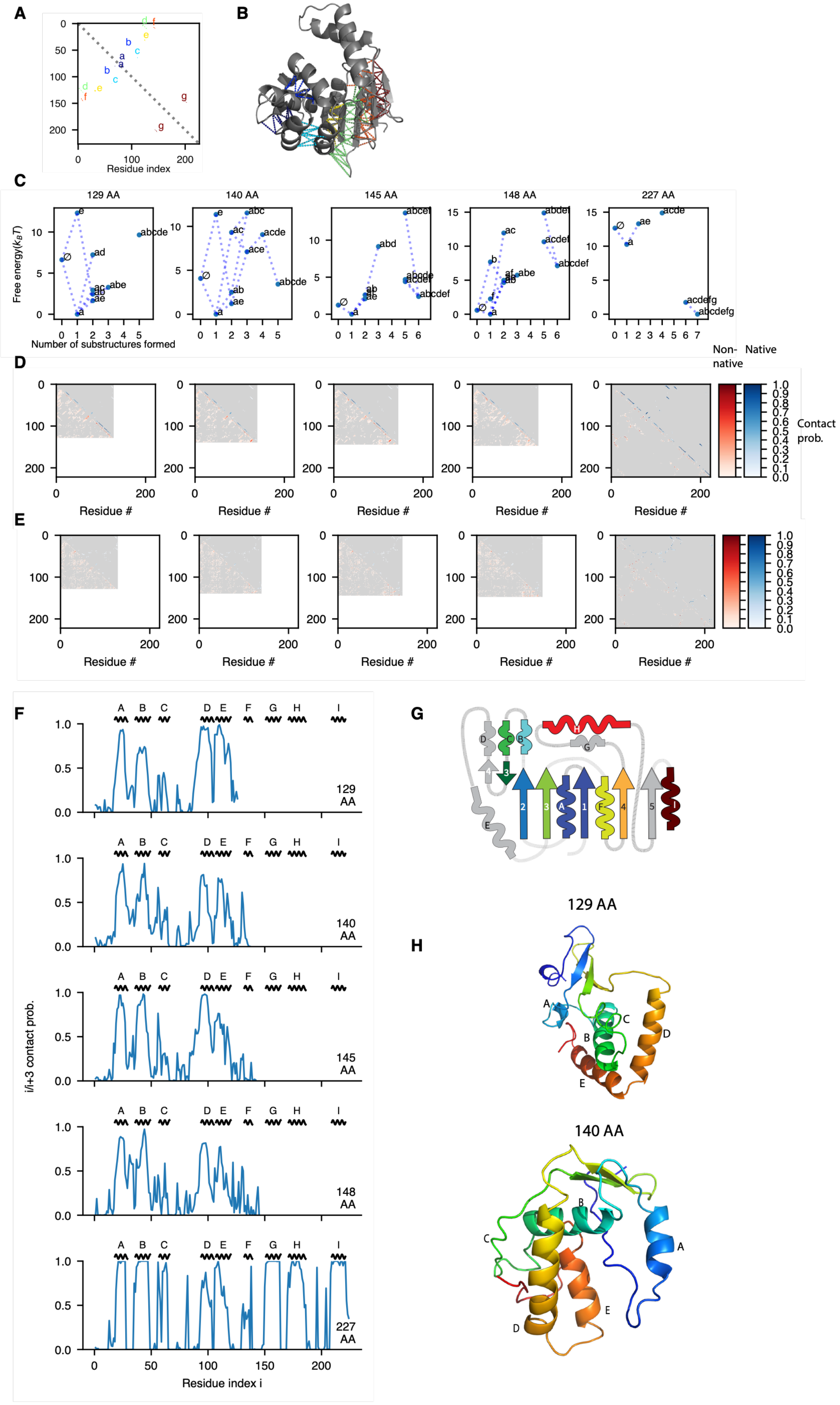
A
B
C
D
E
F
G
H
Residue index
129 AA
140 AA
145 AA
148 AA
227 AA
Free energy($k_BT$)
Number of substructures formed
Non-native
Native
Contact prob.
Residue #
i/i+3 contact prob.
Residue index i

**fig. S3** DBFOLD simulations predict that the CMPK NMP subdomain folds co-translationally while core folding is delayed until after translation

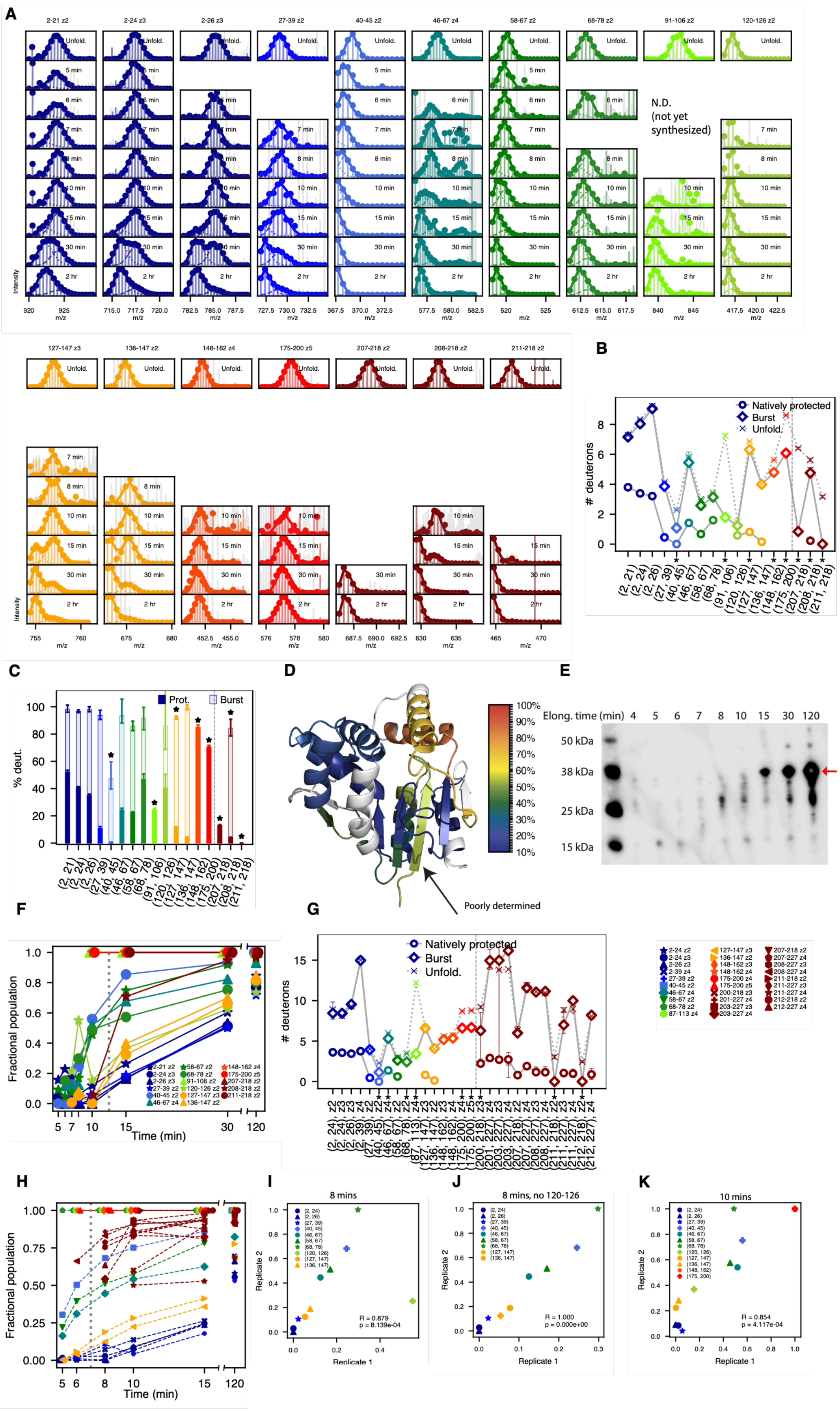
A
2-21 z2
2-24 z3
2-26 z3
27-39 z2
40-45 z2
46-67 z4
58-67 z2
68-78 z2
91-106 z2
120-126 z2
Unfold.
5 min
6 min
7 min
8 min
10 min
15 min
30 min
2 hr
N.D.
(not yet
synthesized)
Intensity
m/z
127-147 z3
136-147 z2
148-162 z4
175-200 z5
207-218 z2
208-218 z2
211-218 z2
B
Natively protected
Burst
Unfold.
### deuterons
C
Prot.
Burst
% deut.
D
Poorly determined
100%
90%
80%
70%
60%
50%
40%
30%
20%
10%
E
Elong. time (min) 4 5 6 7 8 10 15 30 120
50 kDa
38 kDa
25 kDa
15 kDa
F
Fractional population
Time (min)
G
Natively protected
Burst
Unfold.
### deuterons
H
Fractional population
Time (min)
I
8 mins
R = 0.879
p = 8.139e-04
Replicate 1
Replicate 2
J
8 mins, no 120-126
R = 1.000
p = 0.000e+00
Replicate 1
Replicate 2
K
10 mins
R = 0.854
p = 4.117e-04
Replicate 1
Replicate 2

**fig. S4** CMPK synchronized translation + pulse-labeling HDX-MS
**(A)** Mass spectra for all CMPK peptides derived from unfolded control (top row) and all elongation times, as in main text Fig. 2B. Empty panels reflect lack of peptide signal, either because the peptide is not yet synthesized or due to stochastic variabilities in mass-spectrometric detection for low-abundance peptides. **(B)** Number of deuterons associated with natively protected mode (circles), burst-phase mode (the unprotected mode observed early in translation, diamonds) and unfolded state (Xs) for each peptide. Unfolded deuteration levels are corrected for 10 second labeling (see Materials and Methods). Error bars represent 95% confidence intervals from bootstrapping and asterisks denote peptides exhibiting statistically significant burst-phase protection (see Methods). Peptides to the right of the dashed line are only detected after 10 mins (post-translation). **(C)** Same data as (B) but with deuterations expressed as a percentage of maximally deuterated value. **(D)** Percentage deuterations of all peptides are mapped onto the CMPK structure (PDB ID *2cmk*). In general, protection from deuteration correlates with secondary structure. Beta strand 1 appears to show anomalously high deuteration as it is only spanned by large peptides that encompass significant disordered regions, and hence its deuteration cannot be accurately determined. **(E)** Anti-biotin western blot (as in main text Fig. 1C.) showing progress of CMPK synchronized translation reaction as a function of time. This blot reflects the replicate that was used for pulse-labeling experiments shown in main text Fig. 2 and panel (A). Red arrow indicates full-length protein. At around 8 mins, we observe appreciable accumulation of a ~26 kDa translation intermediate, which corresponds to translation of roughly 180 AA from CMPK (with additional ~6 kDa due to the N terminal AviTag + linker). At this stage, the lid subdomain (residues 36-120) has fully emerged from the ribosomal exit tunnel, which typically sequesters the last 30-50 AA. Our observation that lid folding begins around this time (Fig. 2C) strongly suggests that this subdomain folds shortly after its emergence from the ribosome. **(F)** Same as main text Fig. 2C, now showing all peptides with high-quality mass spectra and replicate charge states where present **(G)** Same as (B) for replicate CMPK synchronized translation reaction. (**H)** Same as main text Fig. 2C for a replicate CMPK synchronized translation reaction. As in the first replicate, we observe that lid peptides synchronously begin acquiring protection prior to completion of synthesis, followed by post-translational core folding and rapid NMP subdomain folding. **(I)** To confirm peptides acquire protection in a reproducible sequence across replicates, we compare, for each peptide, the fractional population associated with the natively protected mode at 8 mins in both replicates. Peptides are color coded as in previous panels. A high Spearman correlation coefficient and low p value (shown on the plot) indicate a high degree of reproducibility. **(J)** Same as (I) omitting peptide 120-126 which is poorly fit in the first replicate (see panel A). **(K)** Same as (I) now shown at the 10 mins timepoint.

A

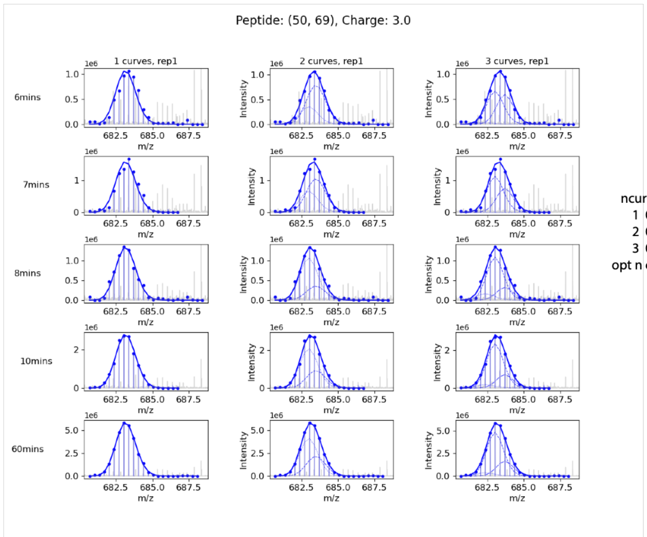


| ncurves | RSS | R^2 | k | n | BIC |
|---|---|---|---|---|---|
| 1 | 0.011483 | 0.972820 | 7 | 95 | -825.096922 |
| 2 | 0.004037 | 0.990444 | 14 | 95 | -892.528740 |
| 3 | 0.003607 | 0.991462 | 21 | 95 | -871.352148 |

opt n curves = 1

B

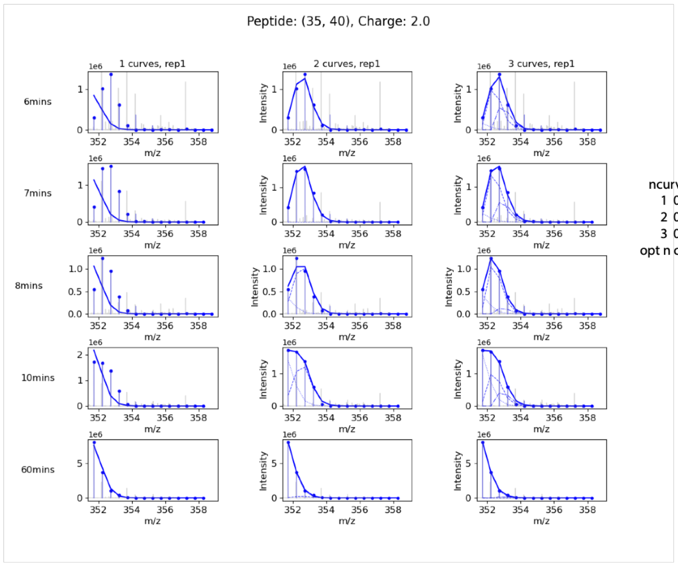


| ncurves | RSS | R^2 | k | n | BIC |
|---|---|---|---|---|---|
| 1 | 0.922705 | -0.259103 | 7 | 30 | -80.640907 |
| 2 | 0.011201 | 0.984716 | 14 | 30 | -189.172831 |
| 3 | 0.003223 | 0.995602 | 21 | 30 | -202.733914 |

opt n curves = 2

C

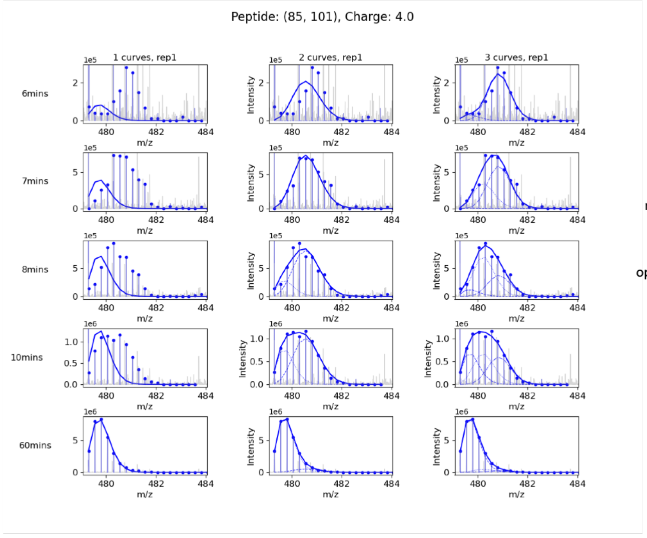


| ncurves | RSS | R^2 | k | n | BIC |
|---|---|---|---|---|---|
| 1 | 0.621762 | -0.570887 | 7 | 80 | -357.903718 |
| 2 | 0.045262 | 0.885646 | 14 | 80 | -536.837308 |
| 3 | 0.011983 | 0.969725 | 21 | 80 | -612.480767 |

opt n curves = 3

**fig. S5** Determination of optimal number of curves for fitting aTS peptides. **(A)-(C)** For indicated peptides and charge states, we globally fit mass spectra from aTS synchronized translation experiments to either one, two or three curves. Mass spectra for each translation time and number of curves, alongside fits, are shown as in main text Fig. 2B. To the right of each set of plots, we indicate the RSS (residual sum of squares), $R^2$ (goodness of fit), k (number of free fitting parameters), n (number of fitted datapoints) and BIC (Bayesian information criterion) associated with each number of curves, as well as the optimal number, determined as described in Materials and Methods.

**A**

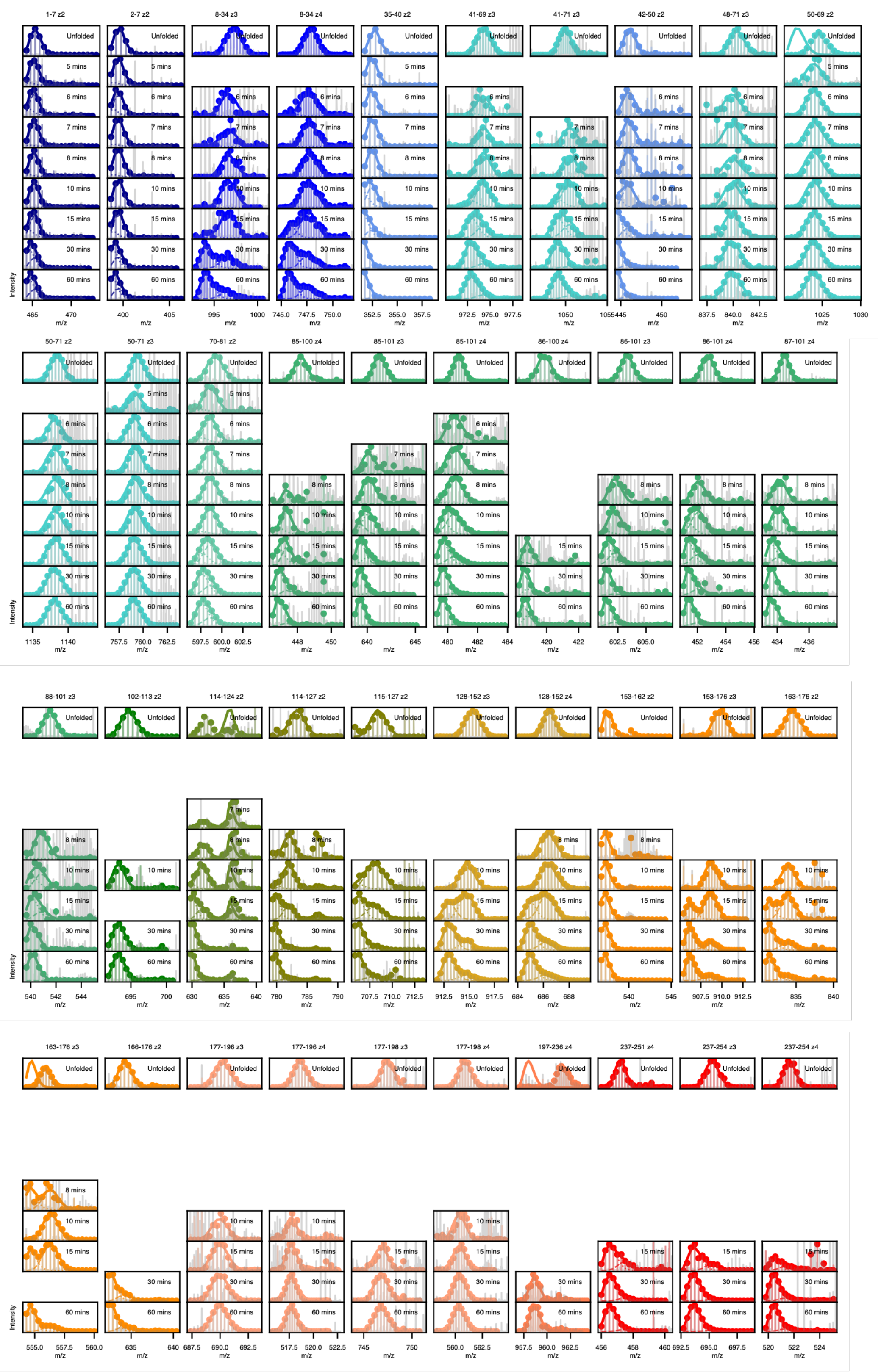
1-7 z2
2-7 z2
8-34 z3
8-34 z4
35-40 z2
41-69 z3
41-71 z3
42-50 z2
48-71 z3
50-69 z2
50-71 z2
50-71 z3
70-81 z2
85-100 z4
85-101 z3
85-101 z4
86-100 z4
86-101 z3
86-101 z4
87-101 z4
88-101 z3
102-113 z2
114-124 z2
114-127 z2
115-127 z2
128-152 z3
128-152 z4
153-162 z2
153-176 z3
163-176 z2
163-176 z3
166-176 z2
177-196 z3
177-196 z4
177-198 z3
177-198 z4
197-236 z4
237-251 z4
237-254 z3
237-254 z4
Unfolded
5 mins
6 mins
7 mins
8 mins
10 mins
15 mins
30 mins
60 mins
Intensity
m/z

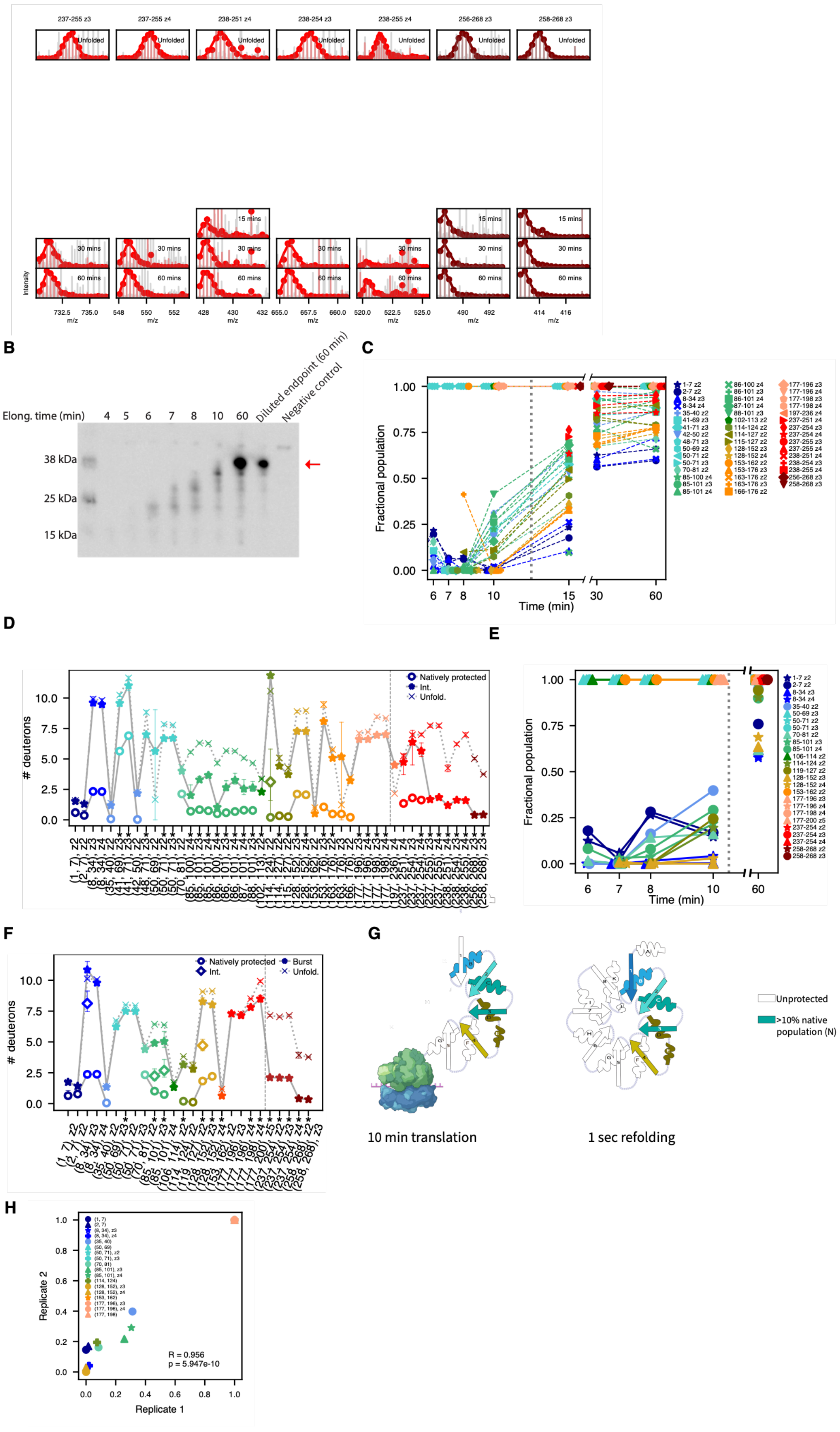
237-255 z3
237-255 z4
238-251 z4
238-254 z3
238-255 z4
256-268 z3
258-268 z3
Unfolded
15 mins
30 mins
60 mins
Intensity
m/z
B
Elong. time (min)
4
5
6
7
8
10
60
Diluted endpoint (60 min)
Negative control
38 kDa
25 kDa
15 kDa
C
Fractional population
Time (min)
D
### deuterons
Natively protected
Int.
Unfold.
E
Fractional population
Time (min)
F
### deuterons
Natively protected
Int.
Burst
Unfold.
G
10 min translation
1 sec refolding
Unprotected
>10% native population (N)
H
Replicate 2
Replicate 1
R = 0.956
p = 5.947e-10

**fig. S6** aTS synchronized translation + pulse-labeling HDX-MS
**(A)** Mass spectra for all aTS peptides derived from unfolded control (top row) and all elongation times, represented as in main text Fig. 2B. Empty panels reflect lack of peptide signal, either because the peptide is not yet synthesized or due to stochastic variabilities in mass-spectrometric detection for low-abundance peptides. **(B)** Anti-biotin western blot (as in main text Fig. 1C.) showing progress of aTS synchronized translation reaction as a function of time. This blot reflects the replicate that was used for pulse-labeling experiments shown in main text Fig. 2 and panel (A). Red arrow indicates full-length protein. (**C)** Same as main text Fig. 2G, now showing all peptides with high-quality mass spectra and replicate charge states where present. **(D)** Number of deuterons associated with natively protected mode (circles), intermediately protected mode where present (diamonds), burst-phase mode (the deprotected mode observed early in translation, asteriks) and unfolded state (Xs) for each peptide. Unfolded deuteration are corrected for 10 second labeling (see Materials and Methods). Error bars represent 95% confidence intervals from bootstrapping and asterisks denote peptides exhibiting statistically significant burst-phase protection (see Methods). Peptides to the right of the dashed line are only detected after 10 mins (post-translation). Various aTS peptides optimally fit to a single mode, For these peptides, only a star and x are shown, representing deuteration of the single global fit mode and the unfolded mode, respectively. **(E)** Same as panel C for a replicate aTS synchronized translation reaction. **(F)** Same as (D) for replicate aTS synchronized translation reaction **(G)** Comparison of folding intermediates populated during aTS translation and refolding from denaturant. Left panel shows secondary structures spanned by peptides showing a natively protected fractional population of at least 10%, provided the deuterium uptake in this native mode is no more than 50% that in the maximally deuterated control, as in main text Fig. 2H, while right panel shows structures spanned by at least one peptide showing a natively protected fractional population of at least 10% at 1 second of refolding from denaturant in the pulse-labeling HDX-MS experiment from Wintrode et. al. 2005 *(50).* For numerical values, see table S4. The co-translational and refolding intermediates largely resemble each other with the exceptions of beta strand 1, which is protected during refolding but not in the co-translational intermediate. **(H)** To confirm peptides acquire protection in a reproducible sequence across replicates, we compare, for each peptide, the fractional population associated with the natively protected mode at 10 mins in both replicates. Peptides are color coded as in previous panels. A high Spearman correlation coefficient and low p value (shown on the plot) indicate a high degree of reproducibility.

| Peptide | Charge | Native Fraction (10 min translation) | Native Fraction (1 sec refolding) |
|---|---|---|---|
| **(1, 7)** | 2 | 0 | 0.04 |
| **(2, 7)** | 2 | 0.02 | N.D. |
| **(8, 11)** | N.S. | N.D. | 0.05 |
| **(8, 34)** | 3 | 0 | N.D. |
| **(8, 34)** | 4 | 0.02 | N.D. |
| **(12, 19)** | N.S. | N.D. | 0.22 |
| **(20, 34)** | N.S. | N.D. | 0.46 |
| **(35, 40)** | 2 | 0.31 | 0.44 |
| **(40, 46)** | N.S. | N.D. | 0.31 |
| **(41, 49)** | N.S. | N.D. | 0.44 |
| **(41, 69)** | 3 | 0.21 | N.D. |
| **(41, 71)** | 3 | 0.22 | N.D. |
| **(42, 50)** | 2 | 0.19 | N.D. |
| **(48, 71)** | 3 | 1 | N.D. |
| **(50, 65)** | N.S. | N.D. | N.D. |
| **(50, 69)** | 2 | 1 | N.D. |
| **(50, 71)** | 2 | 1 | N.D. |
| **(50, 71)** | 3 | 1 | N.D. |
| **(70, 81)** | 2 | 0.08 | 0.16 |
| **(82, 86)** | N.S. | N.D. | 0.79 |
| **(85, 100)** | 4 | 0.22 | N.D. |
| **(85, 101)** | 3 | 0.26 | N.D. |
| **(85, 101)** | 4 | 0.31 | N.D. |
| **(86, 101)** | 4 | 0.26 | 0.43 |
| **(86, 101)** | 3 | 0.08 | 0.43 |
| **(87, 101)** | 4 | 0.16 | N.D. |
| **(88, 101)** | 3 | 0.41 | N.D. |
| **(102, 105)** | N.S. | N.D. | N.D. |
| **(102, 113)** | 2 | 1 | N.D. |
| **(106, 113)** | N.S. | N.D. | N.D. |
| **(114, 118)** | N.S. | N.D. | 0.15 |
| **(114, 124)** | 2 | 0.07 | N.D. |
| **(114, 127)** | 2 | 0.12 | N.D. |
| **(115, 124)** | N.S. | N.D. | 0.11 |
| **(115, 127)** | 2 | 0.11 | N.D. |

| | | | |
|---|---|---|---|
| **(128, 134)** | N.S. | N.D. | N.D. |
| **(128, 152)** | 3 | 0 | 0.06 |
| **(128, 152)** | 4 | 0 | 0.06 |
| **(153, 162)** | 2 | 1 | N.D. |
| **(153, 176)** | 3 | 0 | N.D. |
| **(163, 172)** | N.S. | N.D. | 0.07 |
| **(163, 176)** | 3 | 0 | 0.07 |
| **(163, 176)** | 2 | 0 | 0.07 |
| **(177, 196)** | 4 | 1 | N.D. |
| **(177, 196)** | 3 | 1 | N.D. |
| **(177, 198)** | 4 | 1 | 0.03 |
| **(199, 236)** | N.S. | N.D. | 0.1 |
| **(237, 251)** | N.S. | N.D. | 0.04 |
| **(258, 268)** | N.S. | N.D. | 0.05 |

**Table S4:** aTS co-translational folding intermediate resembles refolding intermediate from denaturant. For each peptide and charge state, we indicate the fraction protected in both the 10 minute co-translational timepoint (our experiment) and the 1 second refolding timepoint (Wintrode et. al. 2005) *(50)*. N.D. indicates peptide is not detected in the respective experiment, and N.S. indicates that the charge state was not specified for peptides in Wintrode et. al. 2005.

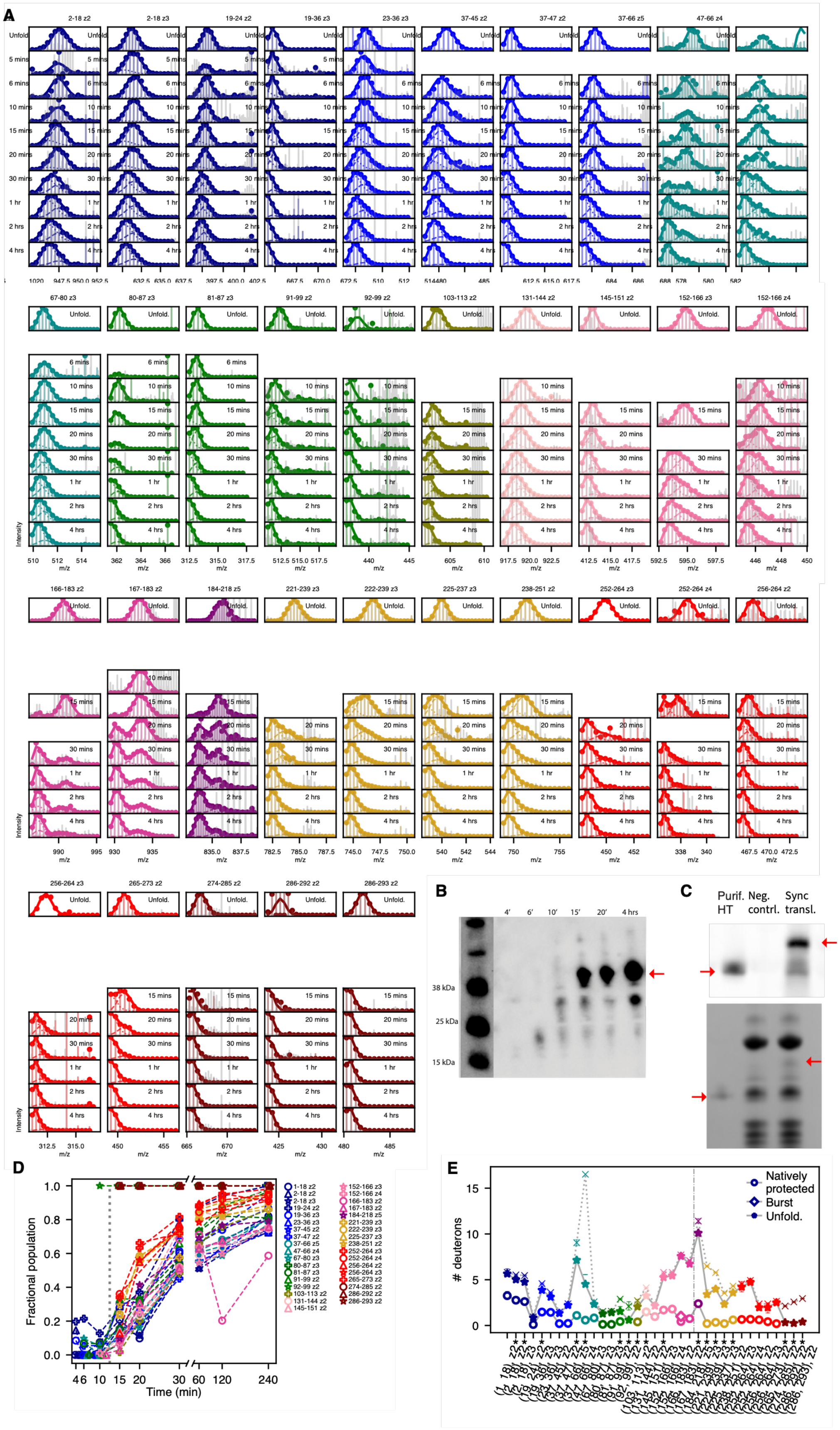
A
B
C
D
E
4'
6'
10'
15'
20'
4 hrs
38 kDa
25 kDa
15 kDa
Purif. HT
Neg. contrl.
Sync transl.
Fractional population
Time (min)
### deuterons
Natively protected
Burst
Unfold.
m/z
Intensity

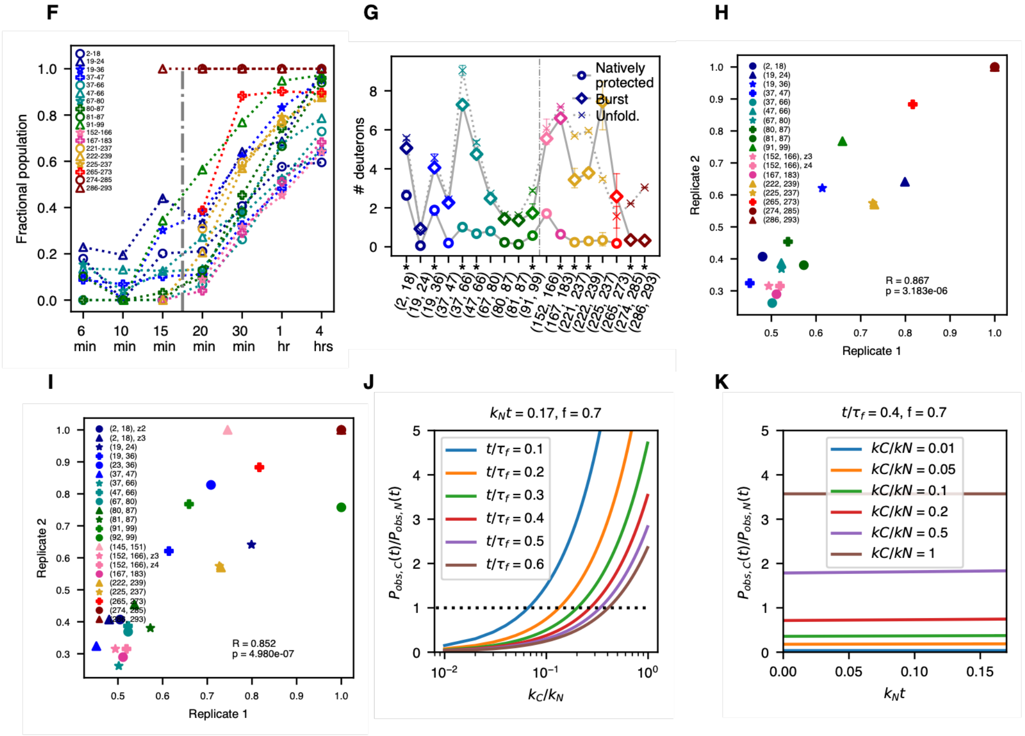


**fig. S7:** HaloTag synchronized translation + pulse-labeling HDX-MS
**(A)** Mass spectra for all HaloTag peptides derived from unfolded control (top row) and all elongation times, represented as in main text Fig. 2B. Empty panels reflect lack of peptide signal, either because the peptide is not yet synthesized or due to stochastic variabilities in mass-spectrometric detection for low-abundance peptides. **(B)** Anti-biotin western blot (as in main text Fig. 1C.) showing progress of HaloTag synchronized translation reaction as a function of time. This blot reflects the replicate that was used for pulse-labeling experiments shown in main text Fig. 2 and panel (A). Red arrow indicates full-length protein. **(C)** Top: To verify HaloTag folds into its native structure in our experimental conditions, we perform a synchronized translation reaction in the presence of 10 μM of fluorescent HaloTag ligand TMR and load the final timepoint on an SDS-PAGE gel which we fluorescently image to confirm covalent TMR bidning (middle lane). As a negative control, we perform an equivalent translation reaction without the HaloTag gene (rightmost lane), and we also load purified HaloTag that was similarly incubated with 10 μM TMR as a positive control (leftmost lane). Translated HaloTag runs larger than purified protein due to the added presence of the N-terminal AviTag and linker. Truncated fragments are also observed to bind TMR to some extend as seen previously *(9)*. Bottom: Coomassie-stained gel from above. In both images, red arrows indicate HaloTag bands. **(D)** Same as main text Fig. 3C now showing all peptides with high-quality mass spectra and replicate charge states where present. **(E)** Same as main text Fig. 3D now showing all peptides with high-quality mass spectra and replicate charge states where present. **(F)** Same as main text Fig. 3C for a replicate HaloTag synchronized translation reaction. As in our first replicate, peptides from across the protein do not begin showing native protection until after translation is complete, and we observe protection in three similar stages, namely 1) rapid folding of C-terminal most helices,

2) folding of C-terminal regions of the core, with limited involvement from the N-terminus, and 3) folding of the majority of N-terminus and lid. We note that translation occurred at a slightly slower and yielded slightly reduced peptide coverage than the initial replicate, particularly near the C-terminus. **(G)** Same as main text Fig. 3C for replicate HaloTag synchronized translation reaction. As in the initial replicate, we observe subtle albeit significant burst-phase protection at various N-terminal peptides. We note that for peptide 91-99, the maximally deuterated control for this replicate gave poor spectra, and we thus used the spectra from the initial replicate instead. We validated this substitution by verifying that the maximally deuterated for this peptide when derived from stalled nascent chains showed nearly identical deuteration. **(H)** To confirm that the order of peptide protection is consistent across replicates, we compare, for each peptide, the fractional population associated with the natively protected mode at 30 mins in both replicates. The x and y axis denote the native population fraction in the two respective replicates at the 30 min timepoint. Peptides are color coded as in previous panels. A high Spearman correlation coefficient and low p value (shown on the plot) indicate a high degree of reproducibility. This plot only includes the subset of peptides with high-quality fits in both replicates that are plotted in main text Fig. 3. **(I)** Same as (H) now showing *all* peptides common to both replicates. **(J)** Plots of eq. 12 in supplementary note 3 as a function of $k_C/k_N$ for indicated parameter values. **(K)** Same as J, now as a function of $k_N t$.

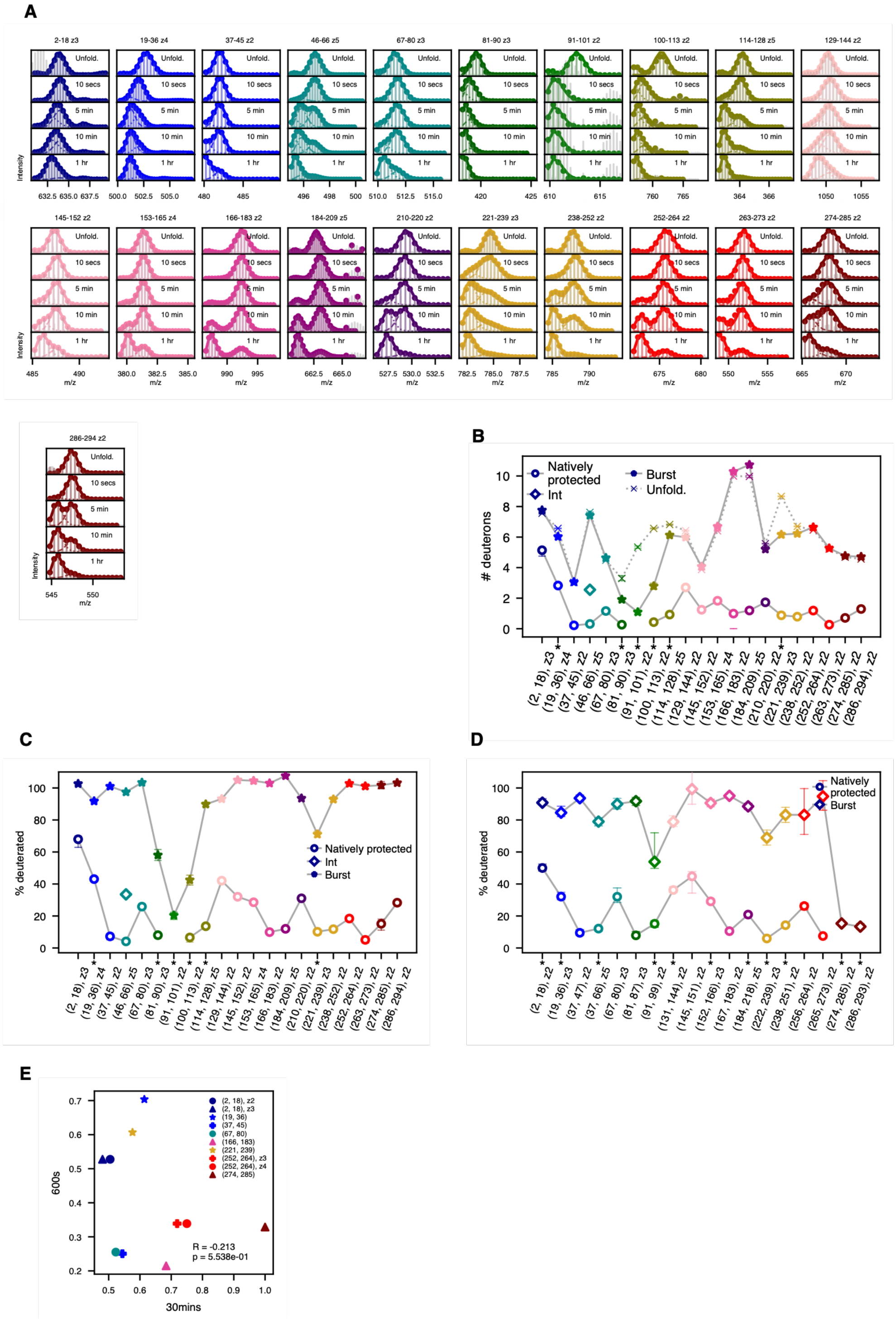


**Fig. S8:** HaloTag refolding from denaturant + pulse-labeling HDX-MS
**(A)** Mass spectra for all HaloTag peptides derived from unfolded control (top row) and all timepoints during refolding from denaturant, as in main text Fig. 2B. Data were obtained from reference *(43)* and reanalyzed using the same global fitting approach applied to co-translational

experiments as described in Materials and Methods . **(B)** Number of deuterons associated with natively-protected mode (circles), intermediately-protected mode where present (stars), burst-phase mode (the deprotected mode observed early in translation, diamonds) and unfolded state (Xs) for each peptide. Error bars represent 95% confidence intervals from bootstrapping and asterisks denote peptides exhibiting statistically significant burst-phase protection (see Methods). (**C)** Same as (B) with all # of deuteron values expressed as a percentage relative to the respective values in the unfolded state. (**D)** Same as main text Fig. 3D, which shows deuteration of native and burst-phase modes in active translation experiment, with all # of deuteron values expressed as a percentage relative to the respective values in the unfolded state. We note that the pattern of burst-phase protection differs from that observed in the refolding experiment, with generally higher protection seen across the N terminus. This strongly suggests that the burst-phase intermediate in co-translational folding differs from that observed in refolding. **(E)** Correlation scatterplot between natively protected fractions in 30 minutes timepoint from first replicate of co-translational folding experiment (x-axis) and 10-minute timepoint of refolding experiment (y-axis). At these timepoints, all peptides have begun to show some degree of native population in both experiments. The low Spearman correlation provides further support that these proceed through different intermediates.

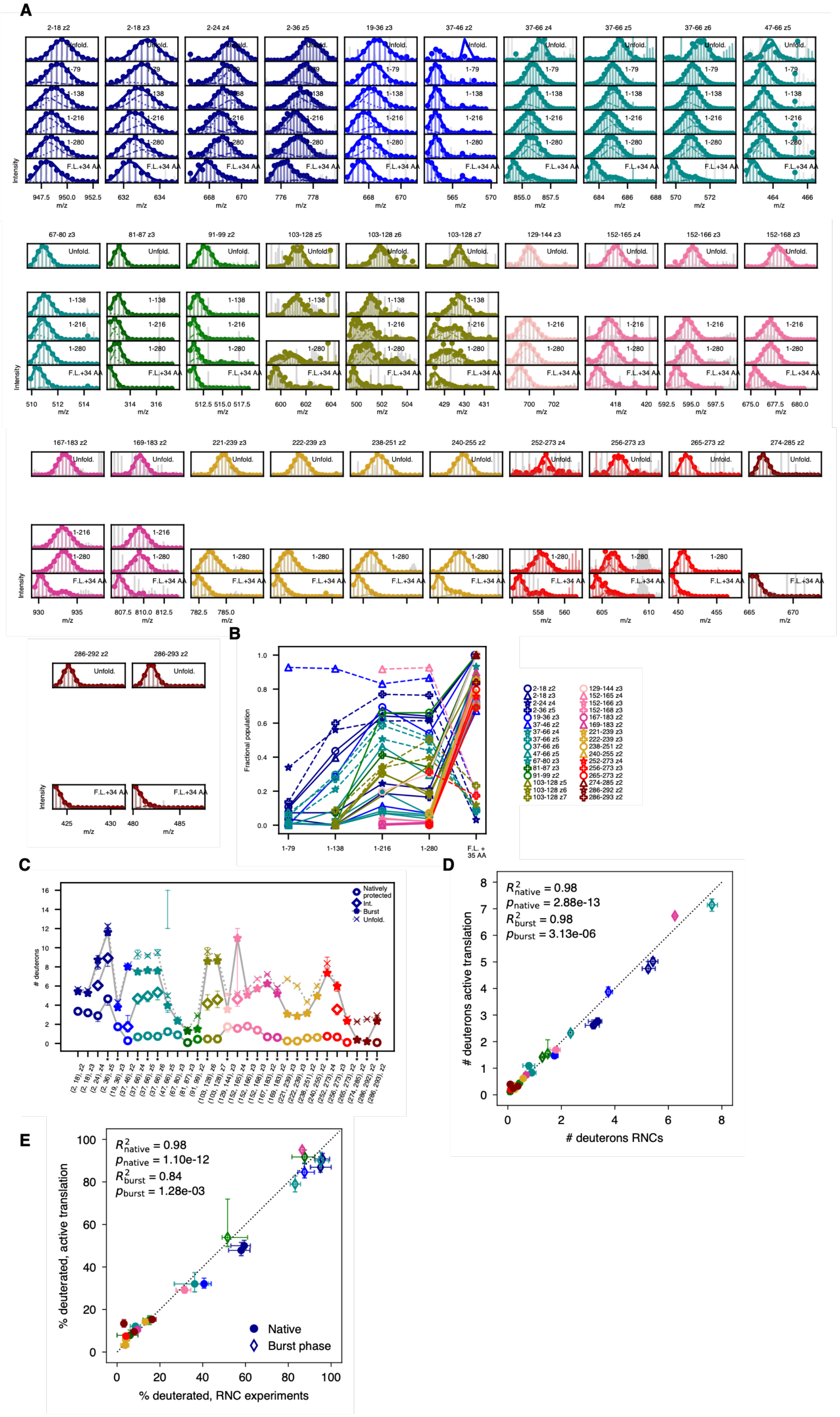
A
2-18 z2
2-18 z3
2-24 z4
2-36 z5
19-36 z3
37-46 z2
37-66 z4
37-66 z5
37-66 z6
47-66 z5
Unfold.
1-79
1-138
1-216
1-280
F.L.+34 AA
Intensity
m/z
67-80 z3
81-87 z3
91-99 z2
103-128 z5
103-128 z6
103-128 z7
129-144 z3
152-165 z4
152-166 z3
152-168 z3
167-183 z2
169-183 z2
221-239 z3
222-239 z3
238-251 z2
240-255 z2
252-273 z4
256-273 z3
265-273 z2
274-285 z2
286-292 z2
286-293 z2
B
Fractional population
1-79
1-138
1-216
1-280
F.L. + 35 AA
C
### deuterons
Natively protected
Int.
Burst
Unfold.
D
$R^2_{native}$ = 0.98
$p_{native}$ = 2.88e-13
$R^2_{burst}$ = 0.98
$p_{burst}$ = 3.13e-06
### deuterons active translation
### deuterons RNCs
E
$R^2_{native}$ = 0.98
$p_{native}$ = 1.10e-12
$R^2_{burst}$ = 0.84
$p_{burst}$ = 1.28e-03
% deuterated, active translation
% deuterated, RNC experiments
Native
Burst phase

**Fig. S9:** HaloTag stalled ribosomal nascent chains (RNCs) + pulse-labeling HDX-MS
(A) Mass spectra for all HaloTag RNC peptides derived from unfolded control (top row) and all translation lengths, as in main text Fig. 4B. Empty panels reflect lack of peptide signal, either because the peptide is not yet synthesized at that length or due to stochastic variabilities in mass-spectrometric detection for low-abundance peptides. **(B)** Same as main text Fig. 4E, now showing all peptides with high-quality mass spectra and replicate charge states where present. **(C)** Number of deuterons associated with natively protected mode (circles), intermediately protected mode where present (stars), burst-phase mode (the deprotected mode observed early in translation, diamonds) and unfolded state (Xs) for each peptide. Unfolded deuteration are corrected for 10 second labeling (see Materials and Methods). Error bars represent 95% confidence intervals from bootstrapping and asterisks denote peptides exhibiting statistically significant protection, relative to the maximally deuterated control corrected for 10 sec labeling, in the deprotected mode (see Materials and Methods). **(D)** Correlation between deuterium uptake associated with native (circles) and burst phase (diamonds) modes for each peptide in RNC and active translation experiments with indicated Pearson correlations. Error bars represent 95% confidence intervals from bootstrapping. **(E)** Same as (D) but with all values normalized by respective maximally deuterated control values. This normalization corrects for the confounding effect of peptide length.
